# Learning mutational graphs of individual tumour evolution from single-cell and multi-region sequencing data

**DOI:** 10.1101/132183

**Authors:** Daniele Ramazzotti, Alex Graudenzi, Luca De Sano, Marco Antoniotti, Giulio Caravagna

## Abstract

**Background:** A large number of algorithms is being developed to reconstruct evolutionary models of individual tumours from genome sequencing data. Most methods can analyze multiple samples collected either through bulk multi-region sequencing experiments or the sequencing of individual cancer cells. However, rarely the same method can support both data types.

**Results:** We introduce `TRaIT`, a computational framework to infer mutational graphs that model the accumulation of multiple types of somatic alterations driving tumour evolution. Compared to other tools, `TRaIT` supports multi-region and single-cell sequencing data within the same statistical framework, and delivers expressive models that capture many complex evolutionary phenomena. `TRaIT` improves accuracy, robustness to data-specific errors and computational complexity compared to competing methods.

**Conclusions:** We show that the application of `TRaIT` to single-cell and multi-region cancer datasets can produce accurate and reliable models of single-tumour evolution, quantify the extent of intra-tumour heterogeneity and generate new testable experimental hypotheses.

## Background

Sequencing data from multiple samples of single tumours can be used to investigate Intra-Tumor Heterogeneity (ITH) in light of evolution [1–3]. Motivated by this observation, several new methods have been developed to infer the “evolutionary history” of a tumour from sequencing data. According to Davis and Navin, there are three orthogonal ways to depict such history [4]: (*i*) with a phylogenetic tree that displays input samples as leaves [5], (*ii*) with a clonal tree of parental relations between putative cancer clones [6–9], and (*iii*) with the order of mutations that accumulated during cancer growth [10–12]. Ideally, the order of accumulating mutations should match the clonal lineage tree in order to reconcile these inferences.

Consistently with earlier works of us [13–18], we here approach the third problem (“mutational ordering”) from two types of data: multi-region bulk and single-cell sequencing.

Bulk sequencing of multiple spatially-separated tumour biopsies returns a noisy mixture of admixed lineages [19–23]. We can analyse these data by first retrieving clonal prevalences in bulk samples (subclonal deconvolution), and then by computing their evolutionary relations [24–31]. Subclonal deconvolution is usually computationally challenging, and can be avoided if we can read genotypes of individual cells via single-cell sequencing (SCS). Despite this theoretical advantage, however, current technical challenges in cell isolation and genome amplification are major bottlenecks to scale SCS to whole-exome or whole-genome assays, and the available targeted data harbours high levels of allelic dropouts, missing data and doublets [32–35]. Thus, the direct application of standard phylogenetic methods to SCS data is not straightforward, despite being theoretically viable [36].

Notice that a common feature of most methods for cancer evolution reconstruction is the employment of the Infinite Sites Assumption (ISA), together with the assumption of no back mutation [24–35], even though recent attempts (e.g., [9]) have been proposed to relax such assumption in order to model relevant phenomena, such as convergent evolutionary trajectories [37].

In this expanding field, we here introduce `TRaIT` (`Temporal oRder of Individual Tumors` – Figures 1 and 2), a new framework for the inference of models of single-tumour evolution, which can analyse, separately, multi-region bulk and single-cell sequencing data, and which allows to capture many complex evolutionary phenomena underlying cancer development.

**Figure 1:**
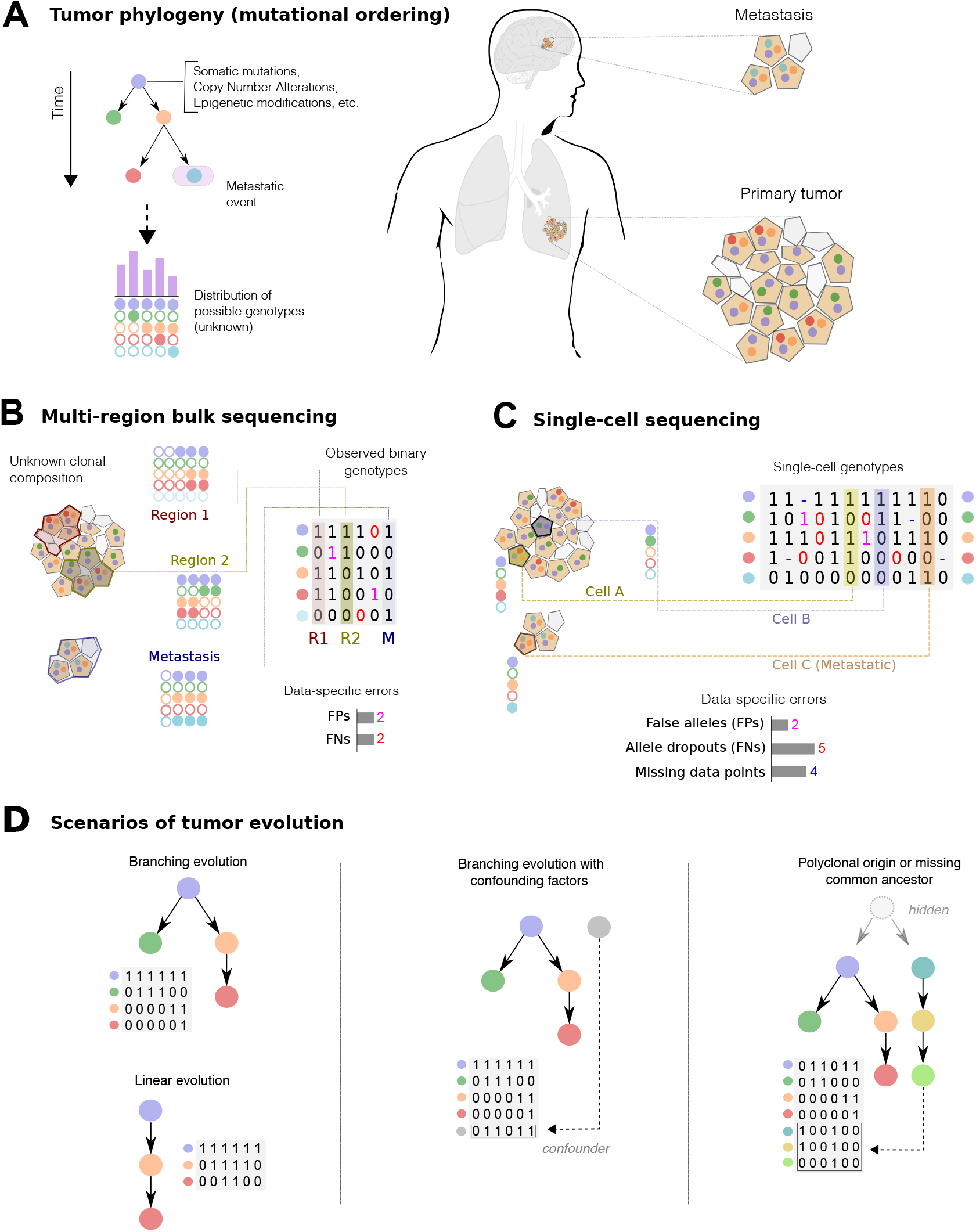
**A.** A tumour phylogeny describes the order of accumulation of somatic mutations, CNAs, epigenetic modifications, etc. in a single tumour. The model generates a set of possible genotypes, which are observed with an unknown spatial and density distribution in a tumour (primary and metastases). **B.** Multi-region bulk sequencing returns a mixed signal from different tumour subpopulations, with potential contamination of non-tumour cells (not shown) and symmetric rates of false positives and negatives in the calling. Thus, a sample will harbour lesions from different tumour lineages, creating spurious correlations in the data. **C.** If we sequence genomes of single cells we can, in principle, have a precise signal from each subpopulation. However, the inference with these data is made harder by high levels of asymmetric noise, errors in the calling and missing data. **D.** Different scenarios of tumour evolution can be investigated via `TRaIT`. (*i*) Branching evolution (which includes linear evolution), (*ii*) Branching evolution with confounding factors annotated in the data, (*iii*) Models with multiple progressions due to polyclonal tumour origination, or to the presence tumour initiating event missing from input data.

**Figure 2:**
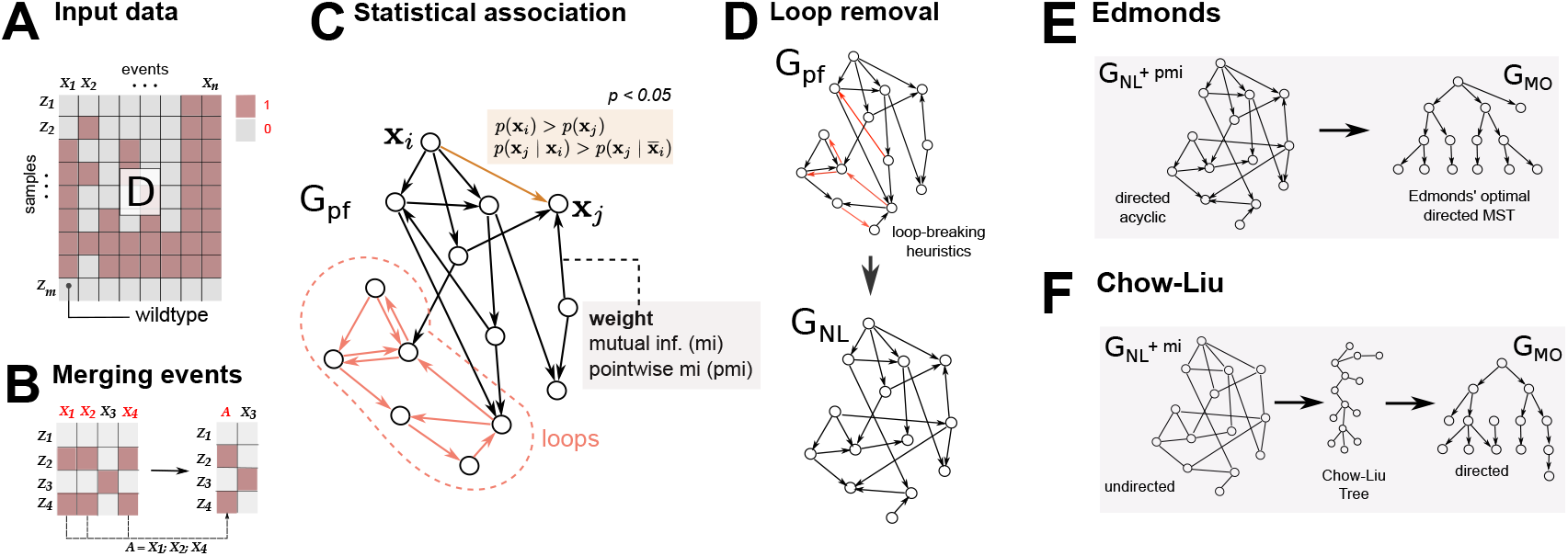
**A.** `TRaIT` processes a binary matrix *D* that stores the presence or absence of a variable in a sample (e.g., a mutation, a CNA, or a persistent epigenetic states). **B.** `TRaIT` merges the events occurring in the same samples (x_1_, x_2_ and x_4_, merged to A), as the statistical signal for their temporal ordering is undistinguishable. The final model include such aggregate events. **C.** We estimate via bootstrap the *prima facie* ordering relation that satisfies Suppes’ conditions (1) for statistical association. This induces a graph *G*_PF_ over variables x_*i*_, which is weighted by information-theoretic measures for variables’ association such as mutual information or pointwise mutual information. **D.** `TRaIT` employs heuristic strategies to remove loops from *G*_PF_ and produce a new graph *G*_NL_ [14]. **E.** `Edmonds`’s algorithm can be used to reconstruct the optimal minimum spanning tree *G*_MO_ that minimises the weights in *G*_NL_; here we use point-wise mutual information (pmi). **F.** `Chow-Liu` is a Bayesian mode-selection strategy that computes an undirected tree as a model of a joint distribution on the annotated variable. Then, we provide edge direction (temporal priority), with Suppes’ condition (1) on marginal probabilities. Therefore, confluences are possible in the output model *G*_MO_ in certain conditions

Compared to other approaches that might scale poorly for increasing sample sizes, our methods show excellent computational performance and scalability, rendering them suitable to anticipate the large amount of genomic data that is becoming increasingly available.

## Results

`TRaIT` is a computational framework that combines Suppes’ probabilistic causation [38] with information theory to infer the temporal ordering of mutations that accumulate during tumour growth, as an extension of our previous work [13–18]. The framework comprises 4 algorithms (`EDMONDS, GABOW, CHOW-LIU` and `PRIM`) designed to model different types of progressions (expressivity) and integrate various types of data, still maintaining a low burden of computational complexity (Figures 1 and 2 – see Methods for the algorithmic details).

In `TRaIT` we estimate the statistical association between a set of genomic events (i.e., mutations, copy number, etc.) annotated in sequencing data by combining optimal graph-based algorithms with bootstrap, hypothesis testing and information theory (Figure 2). `TRaIT` can reconstruct trees and forests – in general, *mutational graphs* – which in specific cases can include confluences, to account for the uncertainty on the precedence relation among certain events. Forest models (i.e., disconnected trees), in particular, can stem for possible polyclonal tumour initiation (i.e., tumours with multiple cells of origin [39]), or the presence of tumour-triggering events that are not annotated in the input data (e.g., epigenetic events) (Figure 1D).

Inputs data in `TRaIT` is represent as binary vectors, which is the standard representation for SCS sequencing and is hereby used to define a unique framework for both multi-region bulk and SCS data (Figure 1A-C). For a set of cells or regions sequenced, the input reports the presence/absence of *n* genomic events, for which `TRaIT` will layout a temporal ordering. A binary representation allows to include several types of somatic lesions in the analysis, such as somatic mutations (e.g., single-nucleotide, indels, etc.), copy number alterations, epigenetic states (e.g., methylations, chromatin modifications), etc. (see the Conclusions for a discussion on the issue of data resolution).

### Performance evaluation with synthetic simulations

We assessed the performance of `TRaIT` with both SCS and multi-region data simulated from different types of generative models.

#### Synthetic data generation

Synthetic single-cell datasets were sampled from a large number of randomly generated topologies (trees or forests) to reflect `TRaIT`’s generative model. For each generative topology, binary datasets were generated starting from the root, with a recursive procedure which we describe for the simpler case of a tree: (*i*) for the root node *x*, the corresponding variable is assigned 1 with a randomly sampled probability *p*(*x* = 1) = *r*, with *r* ~ *U*[0,1]; (*ii*) given a branching node *y* with children *y*_1_, *y*_2_,…, *y_n_*, we sample values for the *n* variables *y*_1_,*y*_2_,…,*y_n_* so that at most one randomly selected child contains 1, and the others are all 0. The recursion proceeds from the root to the leaves, and stops whenever a 0 is sampled or a leaf is reached. Note that we are simulating exclusive branching lineages, as one expects from the accumulation of mutations in single cells under the ISA.

As bulk samples usually include intermixed tumour sub-populations, we simulated bulk datasets by pooling single-cell genotypes generated as described above, and setting simulated variables (i.e., mutations) to 1 (= present) in each bulk sample if they appear in the sampled single-cell genotypes more than a certain threshold. More details on these procedures are reported in Section 2 of the Supplementary Material.

Consistently with previous studies, we also introduced noise in the true genotypes via inflated false positives and false negatives, which are assumed to have highly asymmetric rates for SCS data. For SCS data we also included missing data in a proportion of the simulated variables [11]. Notice that `TRaIT` can be provided with input noise rates, prior to the inference: therefore, in each reconstruction experiment we provided the algorithm with the noise rates used to generate the datasets, even though mild variations in such input values appear not to affect the inference accuracy – as shown in the noise robustness test presented below and in Figure 3D.

**Figure 3:**
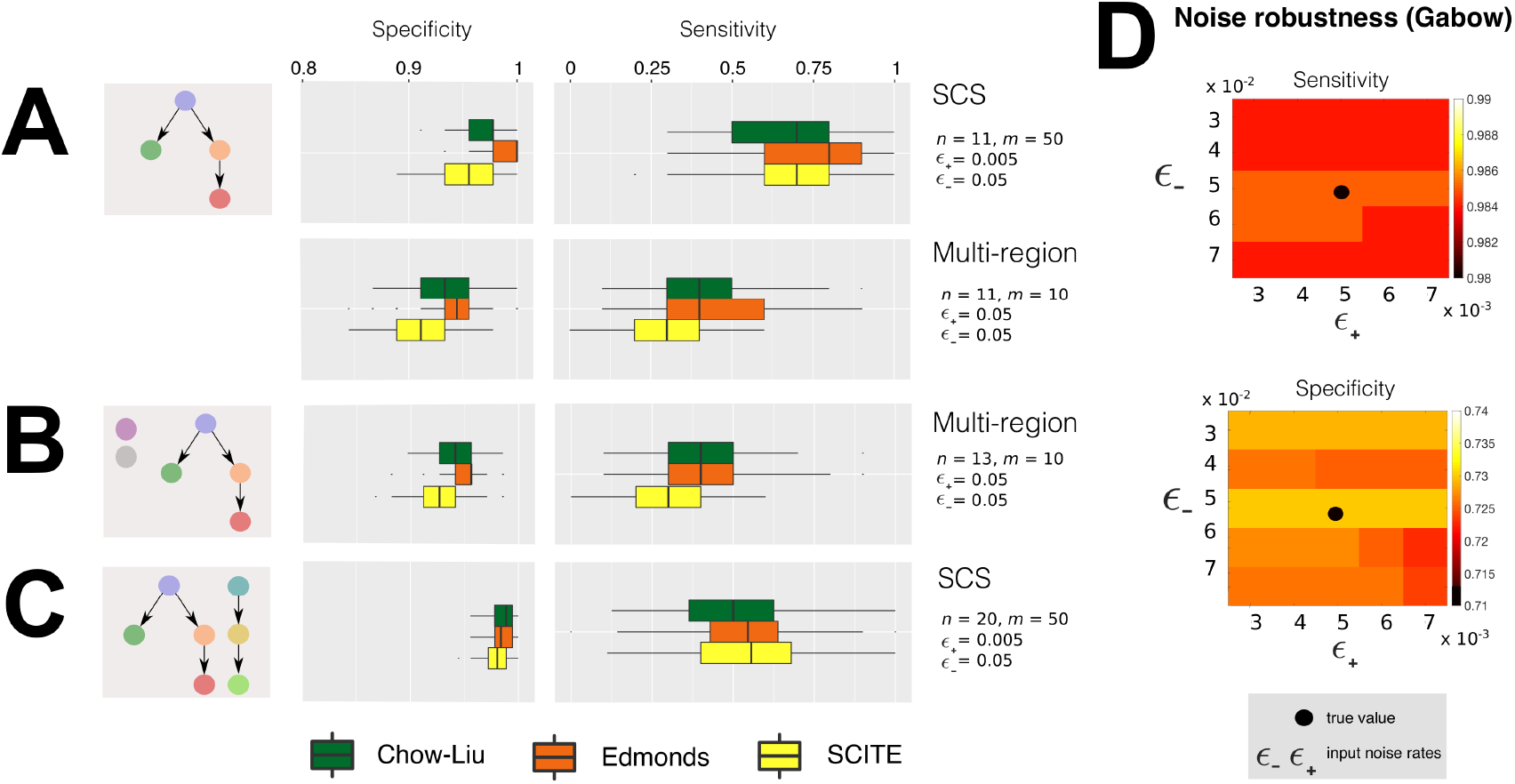
We estimate from simulations the rate of detection of true positives (sensitivity) and negatives (specificity), visualised as *box-plots* from 100 independent points each. We compare `TRaIT`’s algorithms `Edmonds` and `Chow-Liu` with `SCITE`, the state-of-the-art for mutational trees inference in a setting of mild noise in the data, and canonical sample size. In SCS data noise is *ϵ*_+_ = 5 × 10^−3^; *ϵ*_−_ = 5 × 10^−2^, in multi-region *ϵ*_−_ = 5 × 10^−2^. Extensive results for different models, data type, noise and sample size are in Supplementary Figures S3-S16. **A.** Here we use a generative model from [6] (Supplementary Figure S7-B). (left) SCS datasets with *m* = 50 single cells, for a tumour with *n* =11 mutations. (right) Multi-region datasets with *m* =10 spatially separated regions, for a tumour with *n* =11 mutations. **B.** We augment the setting in A-right with 2 random variables (with random marginal probabilty) to model confounding factors, and generated SCS data. **C.** We generated multi-region data from a tumour with *n* = 21 mutations, and a random number of 2 or 3 distinct cells of origin to model polyclonal tumour origination. **D.** Spectrum of average sensitivity and specificity for `Gabow` algorithm included in `TRaIT` (see SM) estimated from 100 independent SCS datasets sampled from the generative model in Supplementary Figure S7-B (*m* = 75, *n* =11). The true noise rates are *ϵ*_+_ = 5 × 10^−3^; *ϵ*_−_ = 5 × 10^−2^; we scan input *ϵ*_+_ and *ϵ*_−_ in the ranges: *ϵ*_+_ = (3,4,5,6,7) × 10^−3^ and 3 × 10^−2^ ≤ *ϵ*_−_ = ≤ 7 × 10^−2^.

With a total of ~140.000 distinct simulations, we could reliably estimate the ability to infer true edges (sensitivity) and discriminate false ones (specificity); further details on parameter settings are available in Section 6 of the Supplementary Material.

In particular, we compared `TRaIT`’s algorithms to `SCITE`, the state-of-the-art to infer mutational trees from SCS data [11]. We could not include `OncoNEM` [7] – the benchmark tool for clonal deconvolution – in the comparison, as its computational performance did not scale well with our large number of tests.

In the Main Text we show results for the `Edmonds` and `Chow-Liu` algorithms, included in `TRaIT`, and `SCITE`, in a selected number of relevant experimental scenarios. To improve readability of the manuscript, we leave to the Supplementary Material a comprehensive presentation of the results for `Gabow, Prim` and other approaches [13,14].

#### Results from scenario (i), branching evolution

To simulate branching evolution [19], we generated a large number of independent datasets from single-rooted tree structures. In particular, we employed three control polyclonal topologies taken from [6] (Supplementary Figure 7) and 100 randomly generated topologies, with a variable number of nodes (i.e., alterations) in the range *n* ∈ [5; 20]. Such generative models were first used to sample datasets with different number of sequenced cells (*m* = 10, 50,100). In addition to the noise-free setting, we perturbed data by introducing plausible and highly asymmetric noise rates (i.e., *ϵ*_+_ = *ϵ*_−_ = 0 (*noise-free*); *ϵ*_+_ = 0.005, *ϵ*_−_ = 0.05; *ϵ*_+_ = 0.02, *ϵ*_−_ = 0.2.). The same generative topologies were then used to sample multi-region datasets with different number of regions (*m* = 5,10, 20), and symmetric noise rates (*ϵ*_+_ = *ϵ*_−_ = 0,0.05, 0.2).

In Figure 3A we show two selected experimental settings, which are characteristic of the general trends observed on all tests. In particular, one can notice that all the techniques achieve high sensitivity and specificity with SCS data, and significantly lower scores with multi-region data from the same topology; `Edmonds` displays in general the best results with SCS data (medians ~ 0.8 and ~ 1).

From the results in all simulation settings (Supplementary Figures 8 and 9 for the SCS case; Supplementary Figures 13 and 14 for the multi-region case), we observe that the overall performance significantly improves for lower noise levels and larger datasets across for all the algorithms, a general result that is confirmed in the other experimental scenarios. In particular, with SCS data, `Edmonds` and `SCITE` display similar sensitivity, even though the latter presents (on average) lower specificity, which might point to a mild-tendency to overfit. Results on multi-region data display similar trends, with `Edmonds` showing the overall best performance and `SCITE` showing slightly lower performance, especially with small datasets and/or low noise levels.

We also specify that, as `TRaIT`’s algorithms share the same constraints in the search space and several algorithmic properties, the reduced variance observed across settings is expected.

#### Results from scenario (ii), confounding factors

To investigate the impact of possible confounding factors on inference accuracy, we introduced in the datasets from scenario (*i*) a number of random binary variables totally unrelated to the progression. More in detail, we inserted around *n* × 10% additional random columns in all datasets with *n* input variables; each additional column is a repeated sampling of a biased coin, with bias uniformly sampled among the marginals of all events.

The performance of `TRaIT` and `SCITE` in a selected setting for the multi-region case is shown in Figure 3B. Surprisingly, the introduction of confounding factors does not impact the performance significantly. In fact, despite two extra variables annotated in the data that are unrelated to the progression, most algorithms still discriminate the true generative model. Similar results are achieved in the SCS case (Supplementary Figure 10).

#### Results from scenario (iii), forest models

Forest topologies can be employed as generative models of tumours initiated by multiple cells, or of tumours whose initiation is triggered by events that are not annotated in the input data. In this test we randomly generated forests with a variable number of distinct disconnected trees, thus assuming that no mutations are shared across the trees. In detail, we generated 100 random forest topologies, with *n* = 20 nodes and *q* < 5 distinct roots (i.e., disconnected trees), both in the SCS and the multi-region case.

The performance of the tested algorithms in a selected experimental scenario with SCS is shown in Figure 3C. All algorithms display a clear decrease in sensitivity, with respect to the single-rooted case with similar values of noise and sample size. In the SCS case the performance remarkably increases with larger datasets (median values ~ 0.75 with *m* = 100 samples in the noise-free case; Supplementary Figure 11). `Edmonds` shows the best tradeoff between sensitivity and specificity, whereas `SCITE` confirms a mild tendency to overfit for small datasets, yet being very robust against noise. Results from multi-region analysis show an overall decrease in performance (Supplementary Figure 16).

#### Robustness to variations in noise input values

Similarly to other tools, e.g., [7,11], our algorithms can receive rates of false positives and negatives in the data (*ϵ*_+_ and *ϵ*_−_) as input. Thus, we analyzed the effect of miscalled rates on the overall performance. More in detail, we analyzed the variation of the performance of `Gabow` and `SCITE`, on a dataset generated from a generative tree with intermediate complexity (“*Medium*” topology in Supplementary Figure 7), with *n* =11 nodes and *m* = 75 samples, *ϵ*_+_ = 5 × 10^−3^ and *ϵ*_−_ =5 × 10^−2^. We scanned 25 possible combinations of input *ϵ*_+_ and *ϵ*_−_ in the following ranges: *ϵ*_+_ = (3, 4, 5, 6, 7) × 10^−3^ and *ϵ*_−_ = (3, 4, 5, 6, 7) × 10^−2^. Results in Figure 3D and Supplementary Tables 4 and 5 show no significant variations of the performance with different combinations of input values for *ϵ*_+_ and *ϵ*_−_, for both algorithms. This proves that the accuracy of the inference is robust to variations in the noise input values, as long as they are reasonably close to the real value. This evidence also supports our algorithmic design choice which avoids sophisticate noise-learning strategies in `TRaIT`, a further reason that speeds up computations.

#### Missing data

Significant rates of missing data are still quite common in SCS datasets, mainly due to amplification biases during library preparation. We evaluated the impact of missing data by using 20 benchmark single-cell datasets which were generated from a tree with *n* =11 nodes (Supplementary Figure 7). For every dataset we simulated the calling of mutations from *m* = 75 single sequenced cells, and in half of the cases (i.e., 10 datasets) we also imputed extra error rates in the data to model sequencing errors. In particular, we introduced false positives and false negative calls with rates *ϵ*_+_ = 0.005 and *ϵ*_−_ = 0.05. On top of this, for each of the 20 datasets we generated 5 configurations of missing data (uniformly distributed), using as measure the percentage *r* of missing data over the total number of observations. A total of 100 distinct datasets have been obtained using *r* = 0, 0.1,0.2, 0.3,0.4 (i.e., up to 40% missing data). As `SCITE` can explicitly learn parameters from missing data, we run the tool with no further parameters. Instead, for `TRaIT`’s algorithms, we performed the following procedure: for each dataset D with missing data, we imputed the missing entries via a standard Expectation-Maximization (EM) algorithm, repeating the procedure to generate 100 complete datasets (*D*_1_,…, *D*_100_). To asses the performance of each algorithm, we computed the fit to all the 100 datasets, and selected the solution that maximised the likelihood of the model.

We present in Figure 4 the results of this analysis for `Edmonds` and `Chow-Liu` algorithms included in `TRaIT`, and for `SCITE`; results for `Gabow` and `Prim` algorithms are presented in Supplementary Figure 12. In general, missing data profoundly affect the performance of all methods. `SCITE` shows overall more robust sensitivity, in spite of slightly worse specificity. The performance is always significantly improved when data do not harbour noise and, in general, is reasonably robust up to 30% missing data.

**Figure 4:**
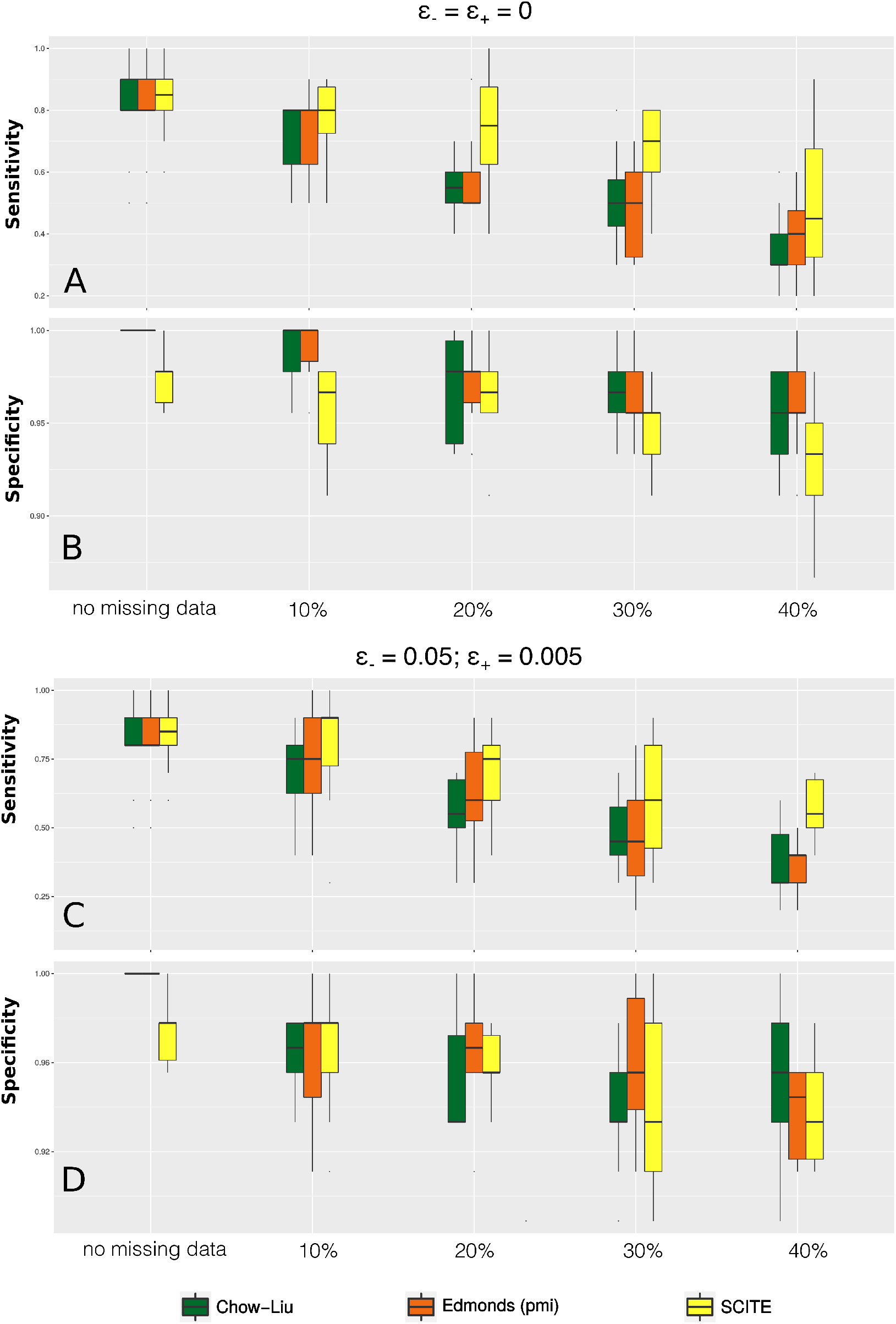
Sensitivity and specificity for different percentages *r* of missing entries, namely, *r* = (0, 0.1,0.2,0.3,0.4) as a function of the number of variables in the data, and different levels of noise: (*i*) *ϵ*_+_ = *ϵ*_−_ = 0 and (*ii*) *ϵ*_+_ = 0.005, *ϵ*_−_ = 0.05. The original dataset is generated from a tree with *n* = 11 nodes and *m* = 75 samples (Supplementary Figure 7).

#### Computational time

One of the major computational advantages of `TRaIT` is its scalability, which will be essential in anticipation of the increasingly larger SCS datasets expected in the near future. In this respect, we have observed across all tests a 3× speedup of `TRaIT`’s algorithms on standard CPUs with respect to `SCITE`, and a 40× speedup with respect to `OncoNEM` (Supplementary Table 6).

### Analysis of patient-derived multi-region data for a MSI-high colorectal cancer

We applied `TRaIT` to 47 nonsynonymous point mutations and 11 indels detected via targeted sequencing in patient `P3` of [40]. This patient has been diagnosed with a moderately-differentiated MSI-high colorectal cancer, for which 3 samples are collected from the primary tumour (`P3-1, P3-2`, and `P3-3`) and two from a right hepatic lobe metastasis `L-1` and `L-2` (Figure 5A). To prepare the data for our analyses, we first grouped mutations occurring in the same regions. We obtained: (*a*) a `clonal` group of 34 mutations detected in all samples (*b*) a `subclonal` group of 3 mutations private to the metastatic regions, and (*c*) 8 mutations with distinct mutational profiles. The clonal group contains mutations in key colorectal driver genes such as APC, KRAS, PIK3CA and `TP53` [15],

**Figure 5:**
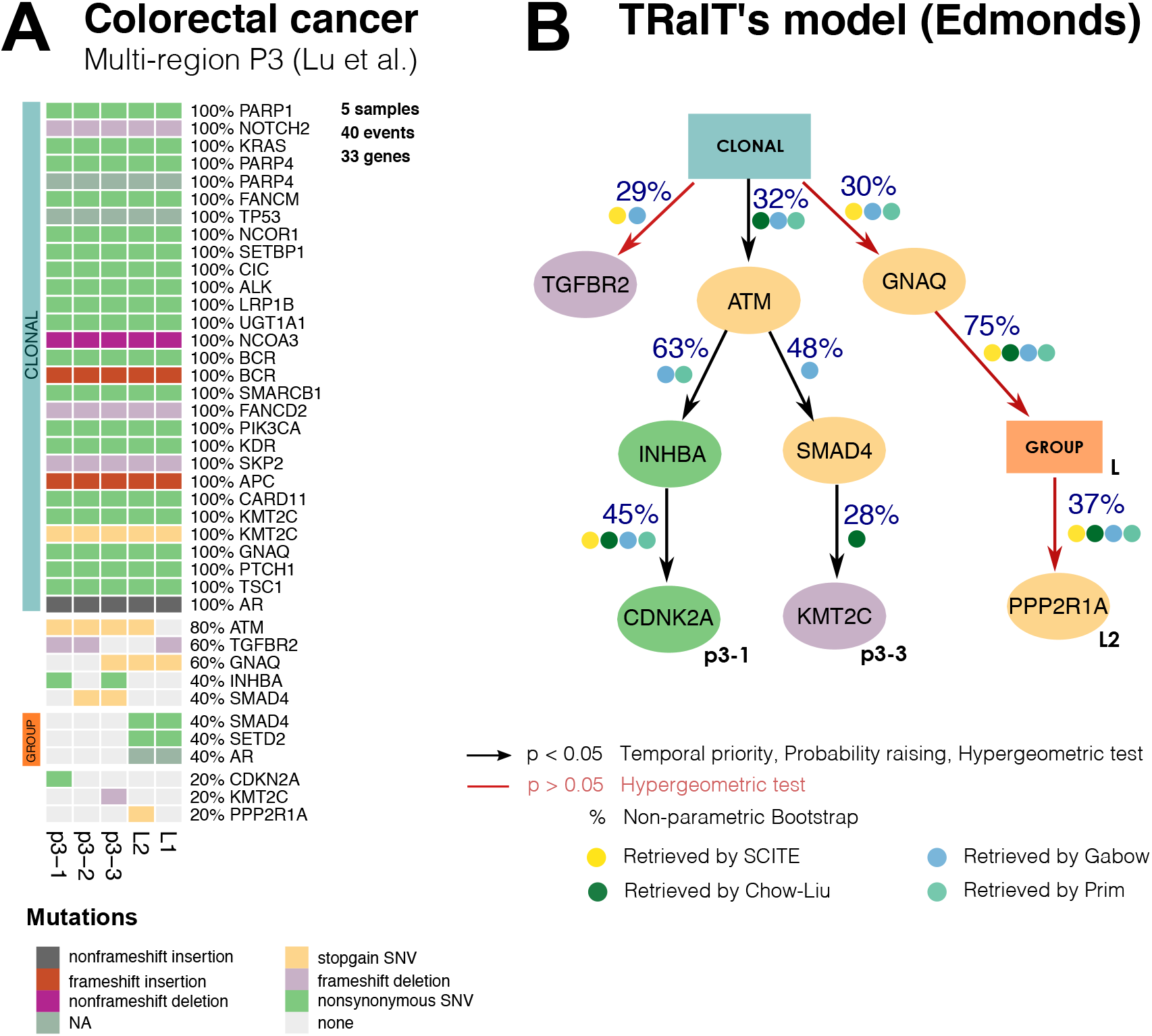
**A.** Multi-region sequencing data for a MSI-high colorectal cancer from [40], with three regions of the primary cancer: `p3-1, p3-2` and `p3-3`, and two of one metastasis: `L-1` and `L-2`. To use this data with `TRaIT` we merge mutations occur in the same samples, obtaining a clonal group of 34 mutations and a sublclonal group. **B.** The model obtained by `Edmonds` including confidence measures, and the overlap in the predicted ordering obtained by `SCITE, Chow-Liu, Gabow` and `Prim` (Supplementary Figure S21). All edges, in all models, are statistically significant for conditions (1). Four of the predicted ordering relations are consistently found across all `TRaIT`’s algorithm, which gives a high-confidence explanation for the formation of the `L2` metastasis. This finding is also in agreement with predictions by `SCITE` (Supplementary Figure S22).

`Edmonds`’s model predicts branching evolution and high levels of ITH among the subclonal populations, consistently with the original phylogenetic analysis by Lu et al. [40] (Figure 5B). In particular, the subclonal trajectory that characterizes the primary regions is initiated by a stopgain SNV in the DNA damage repair gene `ATM`, whereas the subclonal metastatic expansion seems to originate by a stopgain SNV in `GNAQ`, a gene reponsible for diffusion in many tumour types [41]. The model also pictures two distinct trajectories with different mutations in `SMAD4`: a nonsynonimous SNV in group `L`, and a stopgain SNV in two regions of the primary. Interestingly, `SMAD4` regulates cell proliferation, differentiation and apoptosis [42], and its loss is correlated with colorectal metastases [43].

We applied `SCITE` to the same data (Supplementary Figure S22), and compared it to `Edmonds`. Both models depict the same history for the metastatic branch, but different tumour initiation: `SCITE` places the `ATM` mutation on top of the clonal mutations, which appear ordered in a linear chain of 34 events. However, this ordering is uncertain because `SCITE`’s posterior is multi-modal (i.e., several orderings have the same likelihood; Supplementary Figure 22). Further comments on the results, and outputs from other algorithms are available Supplementary Material (Supplementary Figure 21).

### Analysis of patient-derived SCS data for a triple-negative breast cancer

We applied `TRaIT` to the triple-negative breast cancer patient `TNBC` of [34]. The input data consists of single-nucleus exome sequencing of 32 cells: 8 aneuploid (`A`) cells, 8 hypodiploid (`H`) cells and 16 normal cells (`N`) (Figure 6A). Wang et al considered clonal all mutations detected in a control bulk sample and in the majority of the single cells, and as subclonal those undetected in the bulk [34]; all mutations were then used to manually curate a phylogenetic tree (Figure 6B).

**Figure 6:**
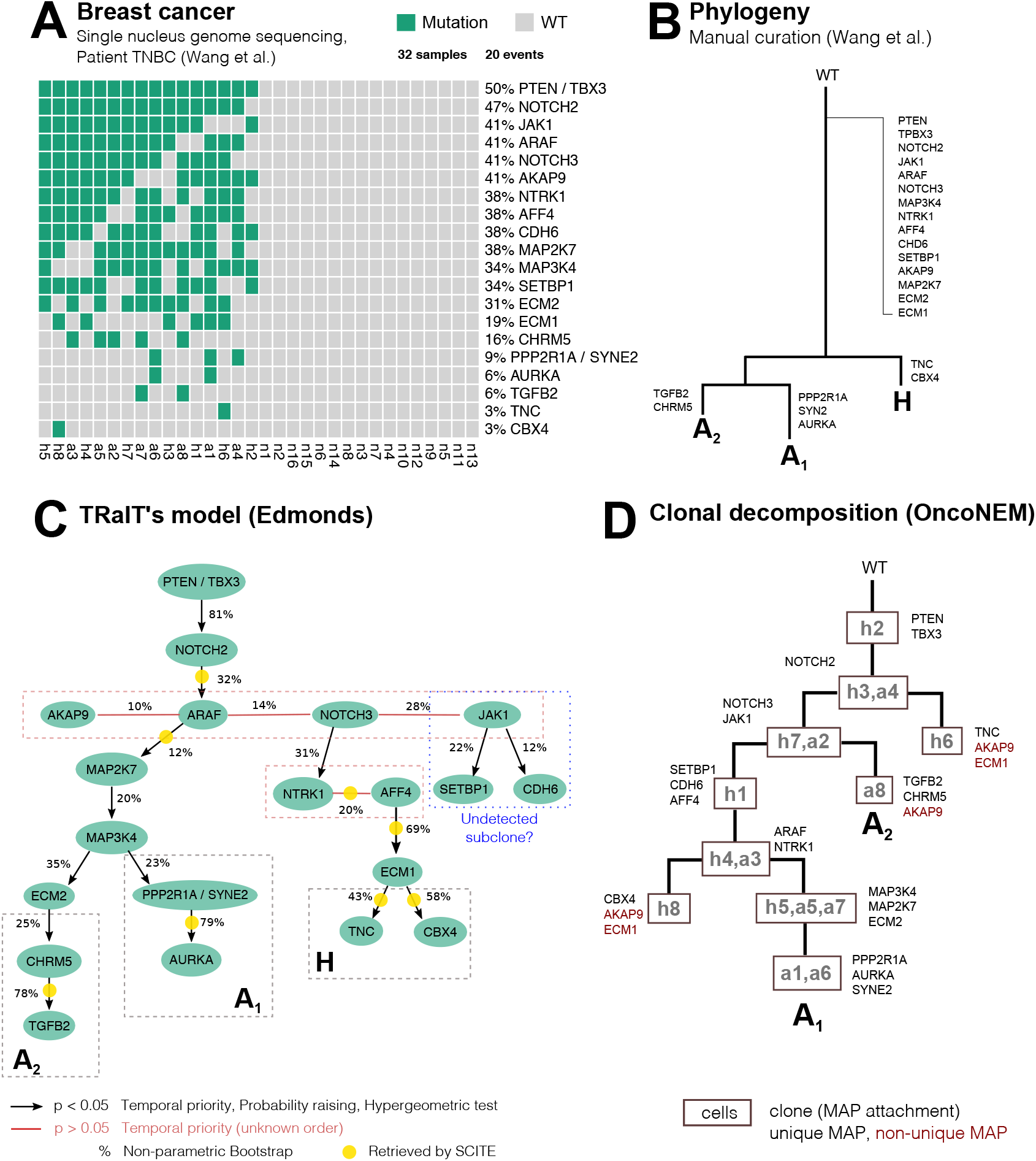
**A.** Input data from single-nucleus sequencing of 32 cells from a triple-negative breast cancer [34]. As the rate of missing values in the original data was around 1%, the authors set all missing data points equal to 0; in the dataset, allelic dropout is equal to 9.73 × 10^−2^, and false discovery equal to 1.24 × 10^−6^. **B** Phylogenetic tree manually curated in [34]. Mutations are annotated to the trunk if they are ubiquitous across cells and a bulk control sample. Subclonal mutations appearing only in more than one cell. **C.** Mutational graph obtained with `Edmonds` algorithm; p-values are obtained by 3 tests for conditions (1) and overlap (hypergeometric test), and edges annotated with a posteriori non-parametric bootstrap scores (100 estimates). For these data, all `TRaIT`’s algorithms return trees (Supplementary Figure S17-18), consistently with the manually curated phylogeny (A). Most edges are highly confident (*p* < 0.05), except for groups of variables with the same frequency which have unknown ordering (red edges). The ordering of mutations in subclones A_1_, A_2_ and tumour initiation has high bootstrap estimates (> 75%). Yellow circles mark the edges retrieved also by `SCITE`. **D.** We also performed clonal tree inference with `OncoNEM`, which predicts 10 clones. Mutations are assigned to clones via *maximum a posteriori* estimates. The mutational orderings of the early clonal expansion of the tumour and of most of the late subclonal events are consistent with `TRaIT`’s prediction.

We run `TRaIT` on all single cells, with nonsynonymous point mutations annotated in 22 genes, and set *ϵ*_+_ = 1.24 × 10^−6^ and *ϵ*_−_ = 9.73 × 10^−2^ as suggested in [34]. All `TRaIT`’s algorithms return tree topologies (Supplementary Figures 17-18); Figure 6C shows the model obtained with `Edmonds`. We integrate the analysis by applying `SCITE` to the same data, and by computing prevalence and evolutionary relations of putative clones with `OncoNEM` as well (Figure 6D).

`TRaIT` provides a finer resolution to the original analysis by Wang et al [34], and retrieves gradual accumulation of point mutations thorough tumour evolution, which highlight progressive DNA repair and replication deregulation. The model also predicts high-confidence branching evolution patterns consistent with subclones A_1_(`PPP2R1A, SYNE2` and `AURKA`), A_2_(`ECM2, CHRM5` and `TGFB2`), and `H`(`NRRK1, AFF4, ECM1, CBX4`), and provides an explicit ordering among clonal mutations in `PTEN, TBX3` and `NOTCH2`, which trigger tumour initiation. Interestingly, `TRaIT` also allows to formulate new hypotheses about a possibly undetected subclone with private mutations in `JAK1, SETBP1` and `CDH6`. Finally, we note that that temporal ordering among mutations in `ARAF`, `AKAP9`, `NOTCH3` and `JAK1` cannot be retrieved, since these events have the same marginal probability in these data.

By applying `SCITE` to these data with the same noise rates, we retrieved 10.000 equivalently optimal trees. The overlap between the first of the returned trees (Supplementary Figure S19) and ours is poor (8 out of 19 edges), and `SCITE`’s models contain a long linear chain of 13 truncal mutations. Clonal deconvolution analysis via `OncoNEM` allowed us to detect 10 clones, their lineages and evolutionary relations. This analysis is in stronger agreement with ours, and the estimated mutational ordering obtained by assigning mutations to clones (via maximum a posteriori, as suggested in [7]) largely overlaps with `TRaIT`’s predictions. This is particularly evident for early events, and for most of the late subclonal ones, exception made for subclone `H`, which is not detected by `OncoNEM`. These results prove that concerted application of tools for mutational and clonal trees inference can provide a picture of ITH at an unprecedented resolution.

## Discussion

In this paper we have introduced `TRaIT`, a computational approach for the inference of cancer evolution models in single tumours. `TRaIT`’s expressive framework allows to reconstruct models beyond standard trees, such as forests, which capture different modalities of tumour initiation (e.g., by multiple cells of origin, or by events missing in available genomic data, such as epigenetic states) and, under certain conditions of data and parameters, confluences. Future works will exploit this latter feature to define a comprehensive modelling framework that accounts for explicit violations of the ISA, in order to model further evolutionary phenomena, such as convergent (parallel) evolution and back mutations [37].

`TRaIT` is based on a binary representation of input data, for both multi-region and singlecell sequencing data. We comment on this design choice concerning the case of multi-region bulk data, because most methods that process bulk data use allelic frequencies and cancer cell fractions to deconvolve the clonal composition of a tumor (see, e.g., [29,30,44]). In this respect, allele frequency-derived inputs provide higher-resolution estimates of the temporal orderings among samples. In fact, if two mutations co-occur in the same set of samples, their relative temporal ordering cannot be determined from a binary input, while this might be possible from their cancer cell fractions. However, despite the lower resolution, a binary representation is still a viable option in multi-region analyses.

First, binary data can describe the presence or absence of a wide range of covariates, which otherwise might be difficult or impossible to represent with allele-frequencies or cancer cell fractions. These include, for instance, complex structural re-arrangements, structural variants, epigenetic modifications, over/under gene expression states and high-level pathway information. The integration of such heterogeneous data types and measurements will be essential to deliver an effective multi-level representation of the life history of individual tumours. Methods that strictly rely on allelic frequencies might need to be extended to accommodate such data types.

Second, binary inputs can be used to promptly analyse targeted sequencing panels, whereas the estimation of subclonal clusters from allele frequencies (i.e., via subclonal deconvolution) requires at least high-depth whole-exome sequencing data to produce reliable results. While it is true that whole-exome and whole-genome assays are becoming increasingly common, many large-scale genomic studies are still relying on targeted sequencing (see, e.g., [45,46]), especially in the clinical setting. A prominent example are assays for longitudinal sampling of circulating tumour DNA during therapy monitoring, which often consist of deep-sequencing target panels derived from the composition of a primary tumour (see, e.g., [47]).

Finally, binary inputs can be obtained for both bulk and single-cell sequencing data, and this in turn allows to use the same framework to study cancer evolution from both data types. This is innovative, and in the future integrative methods might draw inspiration from our approach.

## Conclusions

Intra-tumour heterogeneity is a product of the interplay arising from competition, selection and neutral evolution of cancer subpopulations, and is one of the major causes of drug resistance, therapy failure and relapse [48–52]. For this reason, the choice of the appropriate statistical approach to take full advantage of the increasing resolution of genomic data is key to produce predictive models of tumour evolution with translational relevance.

We have here introduced `TRaIT`, a framework for the efficient reconstruction of single tumour evolution from multiple-sample sequencing data. Thanks to the simplicity of the underlying theoretical framework, `TRaIT` displays significant advancements in terms of robustness, expressivity, data integration and computational complexity. `TRaIT` can process both multiregion and SCS data (separately), and its optimal algorithms maintain a low computational burden compare to alternative tools. `TRaIT`’s assumptions to model accumulation phenomena lead to accurate and robust estimate of temporal orderings, also in presence of noisy data.

We position `TRaIT` in a very precise niche in the landscape of tools for cancer evolution reconstruction, i.e., that of methods for the inference of mutational trees/graphs (not clonal or phylogenetic trees), from binary data (alteration present/absent), and supporting both multiregion bulk and single-cell sequencing data. We advocate the use of `TRaIT` as complementary to tools for clonal tree inference, in a joint effort to quantify the extent of ITH, as shown in the case study on triple negative breast cancer.

## Methods

### Input Data and Data Types

`TRaIT` processes an input binary matrix *D* with *n* columns and *m* rows. *D* stores *n* binary variables (somatic mutations, CNAs, epigenetic states, etc.) detected across *m* samples (single cells or multi-region samples) (Figure 2A). One can annotate data at different resolutions: for instance, one can distinguish mutations by type (missense vs truncating), position, or context (`G>T` vs `G>A`), or can just annotate a general “mutation” status. The same applies for copy numbers, which can be annotated at the focal, cytoband or arm-level. In general, if an entry in *D* is 1, then the associated variable is detected in the sample.

In our framework we cannot disentangle the temporal ordering between events that occur in the same set of samples. These will be grouped by `TRaIT` in a new “aggregate” node, prior to the inference (Figure 2B). `TRaIT` does not explicitly account for back mutations due to loss of heterozygosity. Yet, the information on these events can be used to prepare input data if one matches the copy number state to the presence of mutations. By merging these events we can retrieve their temporal position in the output graph (Supplementary Figure S23).

`TRaIT` supports both multi-region and SCS data. As we expect *D* to contain noisy observations of the unknown true genotypes, the algorithms can be informed of false positives and negatives rates (*ϵ*_+_ ≥ 0 and *ϵ*_−_ ≥ 0). `TRaIT` does not implement noise learning strategies, similarly to `OncoNEM` [11]. This choice is sensitive if the algorithms show stable performance for slight variations in the input noise rates, especially when reasonable estimates of *ϵ*_+_ and *ϵ*_−_ can be known a priori. This feature allows `TRaIT` to be computationally more efficient, as it avoids to include a noise learning routine in the fit. Missing data, instead, are handled by a standard Expectation Maximisation approach to impute missing values: for every complete dataset obtained, the fit is repeated and the model that maximises the likelihood across all runs is returned.

### TRaIT’s Procedure

All `TRaIT`’s algorithms can be summarised with a three-steps skeleton, where the first two steps are the same across all algorithms. Each algorithm will return a unique output model, whose post hoc confidence can be assessed via cross-validation and bootstrap [15].

#### Step 1: assessment of statistical association – Figure 2C

We estimate the statistical association between events by assessing two conditions inspired to Suppes’ theory of probabilistic causation, which is particularly sound in modelling cumulative phenomena [38].

Let *p*(·) be an empirical probability (marginal, joint, conditional etc.) estimated from dataset *D*. Conditions on (*i*) temporal direction and (*ii*) association’s strength are assessed as follows: for every pair of variables *x* and *y* in *D, x* is a plausible temporally antecedent event of *y* if

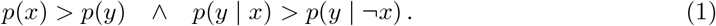

The former condition acts as the Infinite Sites Assumption (ISA), as we assume that alterations are inherited across cell divisions (i.e., somatic): thus, the comparison of marginal frequencies is a proxy to compute the relative ordering among events. The latter condition, instead, implies statistical dependence: *p*(*x,y*) > *p*(*x*)*p*(*y*) [13].

Both conditions are assessed among all variables pairs via non-parametric bootstrap and a one-tailed Mann-Whitney test: only if both conditions are statistically significant at some *α*-level (e.g., 0.05), the edge connecting the variable pair will be included in a prima-facie direct graph *G*_pf_. Edges in *G*_pf_ are candidate to be selected in the final output model, and thus we are reducing the search space via the the above conditions, which are necessary but not sufficient. These conditions have been previously used to define causal approaches for cancer progression [14,15]; see further discussion in Supplementary Material. This step has asymptotic complexity 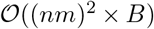 where *B* is the cost of bootstrap and hypothesis testing on each entry in *D*. Notice that this procedure can create disconnected components.

#### Step 2: loop removal – Figure 2D

*G*_PF_ can contain loops, which we have to remove to model an accumulation process. Loops may arise when an arc between a pair of nodes cannot be unequivocally directed, e.g., due to small sample size which leads to uncertain bootstrap estimations. `TRaIT` renders acyclic *G*_PF_ by using heuristic strategies that remove less confident edges (see [14]); the output produced is a new graph *G*_NL_.

#### Step 3: reconstruction of the output model – Figure 2E–F

We render *G*_NL_ a weighted graph by annotating its edges via information-theoretic measures such as point-wise mutual information and the like. Then, we can exploit 4 different off-the-shelf algorithms to reconstruct an output model *G*_MO_ from *G*_NL_. *G*_MO_ will be either a tree or a forest with multiple roots, and the complexity of this step depends on the adopted algorithm. Notably, all algorithms currently incorporated in `TRaIT` have theoretically-optimal worst-case polynomial complexity. We describe two of them (`Edmonds` and `Chow-Liu`), and leave the description of the other techniques (`Gabow` and `Prim`) to the Supplementary Material.
- `Edmonds` is an algorithm for the inference of weighted directed minimum spanning trees [53]: it scan *G*_NL_ to identify the tree that maximises the edges’ weights. Spanning trees have been previously applied to cancer [54,55]. Yet, `TRaIT` is the only framework to constraint spanning trees by condition (1);
- `Chow-Liu`’s algorithm is a method to compute a factorisation of a joint distribution over the input variables [56]. `Chow-Liu` reconstructs undirected trees by definition; we assign the direction to each edge so that the event with higher marginal probability is on top, mirroring condition (1). Confluences in *G*_MO_ can emerge under certain conditions of the observed probabilities, which account for the uncertainty on the temporal precedence among events (technically, in such cases we reconstruct direct acyclic graphs, DAGs – see the Supplementary Material for details).

In all `TRaIT`’s algorithms, if *G*_NL_ includes *k* disconnected components, then the output model *G*_MO_ will include *k* disconnected trees.

In term of complexity, we note that all `TRaIT`’s algorithms are optimal polynomial-time algorithmic solutions to each of their corresponding combinatorial problems. Thus, they scale well with sample size, a problem sometimes observed with Bayesian approaches that cannot compute a full posterior on the model parameters. Quantitative assessment of `TRaIT`’s scalability with large datasets is provided as Supplementary Material (Supplementary Table 7), where we show that many thousands of cells can be processed in a few seconds.

### Tumour evolution scenarios

`TRaIT` can infer mutational graphs in the following scenarios (see Figure 1D):

1. Branching evolution (including linear evolution as subcase): in this case `TRaIT` will return a tree with one root and zero disconnected components.
2. Presence of confounding factors in *D* (e.g., miscalled mutations): `TRaIT` will reconstruct a model with disconnected individual nodes.
3. Polyclonal origin due to multiple cells of tumour origin, or to upstream events triggering tumour development that missing in D (e.g., epigenetic events): `TRaIT` will return models with disconnected components (i.e., forests).

In general, we recommend to apply all `TRaIT`’s algorithms and to compare the output models; the creation of a consensus model is an option to rank the edges detected across several methods, as we show in the case studies.

## Abbreviations

CNA: : Copy-Number Alteration
CT: : Clonal Tree
ISA: : Infinite Sites Assumption
ITH: : Intra-Tumour Heterogeneity
MSI: : Micro-Satellite Instable
SCS: : Single-Cell Sequencing
SNV: : Single-Nucleotide Variant

## Availability of data and materials

`TRaIT` is included in the `TRONCO` R suite for TRanslational ONCOlogy, available at its webpage http://troncopackage.org and mirrored at `Bioconductor`. All data used in this paper are available from the supplementary material of [34] and [40]. We provide the source code and the input data to reproduce the case studies at: `BIMIB-DISCo/TRaIT`.

## Author’s contributions

DR, AG and GC designed the algorithmic framework. DR, LDS and GC implemented the tool. LDS carried out the simulations on synthetic data. Data gathering was performed by DR, AG, LDS and GC. All the authors analyzed the results and interpreted the models. DR, AG, MA and GC wrote the draft of the paper, which all authors reviewed and revised.

## Acknowledgements

We thank Bud Mishra for valuable insights on the effects of parallel evolution to our framework. We also thank Guido Sanguinetti and Yuanhua Huang for useful discussions on the preliminary version of this manuscript.

## References

[1] Niko Beerenwinkel, Roland F Schwarz, Moritz Gerstung, and Florian Markowetz. Cancer evolution: mathematical models and computational inference. Systematic biology, 64(1):e1–e25, 2015.

[2] Russell Schwartz and Alejandro A Schäffer. The evolution of tumour phylogenetics: principles and practice. Nature Reviews Genetics, 2017.

[3] Giulio Caravagna, Ylenia Giarratano, Daniele Ramazzotti, Ian Tomlinson, Trevor A. Graham, Guido Sanguinetti, and Andrea Sottoriva. Detecting repeated cancer evolution from multi-region tumor sequencing data. Nature Methods, 15(9):707–714, 2018.

[4] Alexander Davis and Nicholas E Navin. Computing tumor trees from single cells. Genome biology, 17(1):113, 2016.

[5] Alexandros Stamatakis. Raxml version 8: a tool for phylogenetic analysis and postanalysis of large phylogenies. Bioinformatics, 30(9):1312–1313, 2014.

[6] Ke Yuan, Thomas Sakoparnig, Florian Markowetz, and Niko Beerenwinkel. BitPhylogeny: a probabilistic framework for reconstructing intra-tumor phylogenies. Genome biology, 16(1):1, 2015.

[7] Edith M Ross and Florian Markowetz. OncoNEM: inferring tumor evolution from single-cell sequencing data. Genome biology, 17(1):1, 2016.

[8] Andrew Roth, Andrew McPherson, Emma Laks, Justina Biele, Damian Yap, Adrian Wan, Maia A Smith, Cydney B Nielsen, Jessica N McAlpine, Samuel Aparicio, Alexandre Bouchard-Côté, and Sohrab P Shah. Clonal genotype and population structure inference from single-cell tumor sequencing. Nature methods, 13(7):573–576, 2016.

[9] Hamim Zafar, Anthony Tzen, Nicholas Navin, Ken Chen, and Luay Nakhleh. SiFit: inferring tumor trees from single-cell sequencing data under finite-sites models. Genome Biology, 18(1):178, 2017.

[10] Kyung In Kim and Richard Simon. Using single cell sequencing data to model the evolutionary history of a tumor. BMC bioinformatics, 15(1):27, 2014.

[11] Katharina Jahn, Jack Kuipers, and Niko Beerenwinkel. Tree inference for single-cell data. Genome biology, 17(1):1, 2016.

[12] Sohrab Salehi, Adi Steif, Andrew Roth, Samuel Aparicio, Alexandre Bouchard-Côté, and Sohrab P Shah. ddClone: joint statistical inference of clonal populations from single cell and bulk tumour sequencing data. Genome Biology, 18(1):44, 2017.

[13] Loes Olde Loohuis, Giulio Caravagna, Alex Graudenzi, Daniele Ramazzotti, Giancarlo Mauri, Marco Antoniotti, and Bud Mishra. Inferring tree causal models of cancer progression with probability raising. PloS one, 9(10):e108358, 2014.

[14] Daniele Ramazzotti, Giulio Caravagna, Loes Olde Loohuis, Alex Graudenzi, Ilya Korsunsky, Giancarlo Mauri, Marco Antoniotti, and Bud Mishra. CAPRI: efficient inference of cancer progression models from cross-sectional data. Bioinformatics, 31(18):3016–3026, 2015.

[15] G. Caravagna, A. Graudenzi, D. Ramazzotti, R. Sanz-Pamplona, L. De Sano, G. Mauri, V. Moreno, M. Antoniotti, and B. Mishra. Algorithmic methods to infer the evolutionary trajectories in cancer progression. Proceedings of the National Academy of Sciences, 113(28):E4025–E4034, 2016.

[16] Daniele Ramazzotti. A Model of Selective Advantage for the Efficient Inference of Cancer Clonal Evolution. PhD thesis, Universitá degli Studi di Milano-Bicocca, 2017. arXiv preprint arXiv:1602.07614.

[17] Daniele Ramazzotti, Marco S Nobile, Paolo Cazzaniga, Giancarlo Mauri, and Marco Antoniotti. Parallel implementation of efficient search schemes for the inference of cancer progression models. In Computational Intelligence in Bioinformatics and Computational Biology (CIBCB), 2016 IEEE Conference on, pages 1–6. IEEE, 2016.

[18] Daniele Ramazzotti, Alex Graudenzi, Giulio Caravagna, and Marco Antoniotti. Modeling cumulative biological phenomena with suppes-bayes causal networks. Evolutionary Bioinformatics, 14:1176934318785167, 2018.

[19] Marco Gerlinger, Andrew J Rowan, Stuart Horswell, James Larkin, David Endesfelder, Eva Gronroos, Pierre Martinez, Nicholas Matthews, Aengus Stewart, Patrick Tarpey, et al. Intratumor heterogeneity and branched evolution revealed by multiregion sequencing. New England journal of medicine, 366(10):883–892, 2012.

[20] Elza C de Bruin, Nicholas McGranahan, Richard Mitter, Max Salm, David C Wedge, Lucy Yates, Mariam Jamal-Hanjani, Seema Shafi, Nirupa Murugaesu, Andrew J Rowan, et al. Spatial and temporal diversity in genomic instability processes defines lung cancer evolution. Science, 346(6206):251–256, 2014.

[21] Jianjun Zhang, Junya Fujimoto, Jianhua Zhang, David C Wedge, Xingzhi Song, Jiexin Zhang, Sahil Seth, Chi-Wan Chow, Yu Cao, Curtis Gumbs, et al. Intratumor heterogeneity in localized lung adenocarcinomas delineated by multiregion sequencing. Science, 346(6206):256–259, 2014.

[22] Lucy R Yates, Moritz Gerstung, Stian Knappskog, Christine Desmedt, Gunes Gundem, Peter Van Loo, Turid Aas, Ludmil B Alexandrov, Denis Larsimont, Helen Davies, et al. Subclonal diversification of primary breast cancer revealed by multiregion sequencing. Nature medicine, 21(7):751–759, 2015.

[23] Mariam Jamal-Hanjani, Gareth A Wilson, Nicholas McGranahan, Nicolai J Birkbak, Thomas BK Watkins, Selvaraju Veeriah, Seema Shafi, Diana H Johnson, Richard Mitter, Rachel Rosenthal, et al. Tracking the evolution of non–small-cell lung cancer. New England Journal of Medicine, 376(22):2109–2121, 2017.

[24] Layla Oesper, Ahmad Mahmoody, and Benjamin J Raphael. Inferring intra-tumor heterogeneity from high-throughput dna sequencing data. In Annual International Conference on Research in Computational Molecular Biology, pages 171–172. Springer, 2013.

[25] Francesco Strino, Fabio Parisi, Mariann Micsinai, and Yuval Kluger. TrAp: a tree approach for fingerprinting subclonal tumor composition. Nucleic acids research, 41(17):e165–e165, 2013.

[26] Andrew Roth, Jaswinder Khattra, Damian Yap, Adrian Wan, Emma Laks, Justina Biele, Gavin Ha, Samuel Aparicio, Alexandre Bouchard-Côté, and Sohrab P Shah. PyClone: statistical inference of clonal population structure in cancer. Nature methods, 11(4):396–398, 2014.

[27] Wei Jiao, Shankar Vembu, Amit G Deshwar, Lincoln Stein, and Quaid Morris. Inferring clonal evolution of tumors from single nucleotide somatic mutations. BMC bioinformatics, 15(1):1, 2014.

[28] Habil Zare, Junfeng Wang, Alex Hu, Kris Weber, Josh Smith, Debbie Nickerson, ChaoZhong Song, Daniela Witten, C Anthony Blau, and William Stafford Noble. Inferring clonal composition from multiple sections of a breast cancer. PLoS Comput Biol, 10(7):e1003703, 2014.

[29] Amit G Deshwar, Shankar Vembu, Christina K Yung, Gun Ho Jang, Lincoln Stein, and Quaid Morris. PhyloWGS: reconstructing subclonal composition and evolution from whole-genome sequencing of tumors. Genome biology, 16(1):1, 2015.

[30] Mohammed El-Kebir, Gryte Satas, Layla Oesper, and Benjamin J Raphael. Inferring the mutational history of a tumor using multi-state perfect phylogeny mixtures. Cell Systems, 3(1):43–53, 2016.

[31] Zheng Hu and Christina Curtis. Inferring Tumor Phylogenies from Multi-region Sequencing. Cell systems, 3(1):12–14, 2016.

[32] Nicholas Navin, Jude Kendall, Jennifer Troge, Peter Andrews, Linda Rodgers, Jeanne McIndoo, Kerry Cook, Asya Stepansky, Dan Levy, Diane Esposito, et al. Tumour evolution inferred by single-cell sequencing. Nature, 472(7341):90–94, 2011.

[33] Charles Gawad, Winston Koh, and Stephen R Quake. Dissecting the clonal origins of childhood acute lymphoblastic leukemia by single-cell genomics. Proceedings of the National Academy of Sciences, 111(50):17947–17952, 2014.

[34] Yong Wang, Jill Waters, Marco L Leung, Anna Unruh, Whijae Roh, Xiuqing Shi, Ken Chen, Paul Scheet, Selina Vattathil, Han Liang, et al. Clonal evolution in breast cancer revealed by single nucleus genome sequencing. Nature, 512(7513):155–160, 2014.

[35] Nicholas E Navin. The first five years of single-cell cancer genomics and beyond. Genome research, 25(10):1499–1507, 2015.

[36] Roland F Schwarz, Anne Trinh, Botond Sipos, James D Brenton, Nick Goldman, and Florian Markowetz. Phylogenetic quantification of intra-tumour heterogeneity. PLoS computational biology, 10(4):e1003535, 2014.

[37] Nicholas E Navin and Ken Chen. Genotyping tumor clones from single-cell data. Nature Methods, 13(7):555–556, 2016.

[38] Patrick Suppes. A probabilistic theory of causality. North-Holland Publishing Company Amsterdam, The Netherlands, 1970.

[39] Barbara L Parsons. Many different tumor types have polyclonal tumor origin: evidence and implications. Mutation Research/Reviews in Mutation Research, 659(3):232–247, 2008.

[40] You-Wang Lu, Hui-Feng Zhang, Rui Liang, Zhen-Rong Xie, Hua-You Luo, Yu-Jian Zeng, Yu Xu, La-Mei Wang, Xiang-Yang Kong, and Kun-Hua Wang. Colorectal cancer genetic heterogeneity delineated by multi-region sequencing. PloS one, 11(3):e0152673, 2016.

[41] Chung-Young Kim, Dae Won Kim, Kevin Kim, Jonathan Curry, Carlos Torres-Cabala, and Sapna Patel. GNAQ mutation in a patient with metastatic mucosal melanoma. BMC cancer, 14(1):516, 2014.

[42] Hafid Alazzouzi, Pia Alhopuro, Reijo Salovaara, Heli Sammalkorpi, Heikki Järvinen, Jukka-Pekka Mecklin, Akeseli Hemminki, Simo Schwartz, Lauri A Aaltonen, and Diego Arango. SMAD4 as a prognostic marker in colorectal cancer. Clinical Cancer Research, 11(7):2606–2611, 2005.

[43] Xuemei Li, Baoquan Liu, Jianbing Xiao, Ying Yuan, Jing Ma, and Yafang Zhang. Roles of VEGF-C and SMAD4 in the lymphangiogenesis, lymphatic metastasis, and prognosis in colon cancer. Journal of Gastrointestinal Surgery, 15(11):2001, 2011.

[44] Yuchao Jiang, Yu Qiu, Andy J Minn, and Nancy R Zhang. Assessing intratumor heterogeneity and tracking longitudinal and spatial clonal evolutionary history by next-generation sequencing. Proceedings of the National Academy of Sciences, 113(37):E5528–E5537, 2016.

[45] Samra Turajlic, Hang Xu, Kevin Litchfield, Andrew Rowan, Stuart Horswell, Tim Chambers, Tim OBrien, Jose I Lopez, Thomas BK Watkins, David Nicol, et al. Deterministic evolutionary trajectories influence primary tumor growth: Tracerx renal. Cell, 173(3):595–610, 2018.

[46] Samra Turajlic, Hang Xu, Kevin Litchfield, Andrew Rowan, Tim Chambers, Jose I Lopez, David Nicol, Tim OBrien, James Larkin, Stuart Horswell, et al. Tracking cancer evolution reveals constrained routes to metastases: Tracerx renal. Cell, 173(3):581–594, 2018.

[47] Christopher Abbosh, Nicolai J Birkbak, Gareth A Wilson, Mariam Jamal-Hanjani, Tudor Constantin, Raheleh Salari, John Le Quesne, David A Moore, Selvaraju Veeriah, Rachel Rosenthal, et al. Phylogenetic ctdna analysis depicts early-stage lung cancer evolution. Nature, 545(7655):446, 2017.

[48] Serena Nik-Zainal, Peter Van Loo, David C Wedge, Ludmil B Alexandrov, Christopher D Greenman, King Wai Lau, Keiran Raine, David Jones, John Marshall, Manasa Ramakrishna, et al. The life history of 21 breast cancers. Cell, 149(5):994–1007, 2012.

[49] Robert J Gillies, Daniel Verduzco, and Robert A Gatenby. Evolutionary dynamics of carcinogenesis and why targeted therapy does not work. Nature Reviews Cancer, 12(7):487–493, 2012.

[50] Rebecca A Burrell, Nicholas McGranahan, Jiri Bartek, and Charles Swanton. The causes and consequences of genetic heterogeneity in cancer evolution. Nature, 501(7467):338–345, 2013.

[51] Bert Vogelstein, Nickolas Papadopoulos, Victor E Velculescu, Shibin Zhou, Luis A Diaz, and Kenneth W Kinzler. Cancer genome landscapes. science, 339(6127):1546–1558, 2013.

[52] Andrea Sottoriva, Inmaculada Spiteri, Sara GM Piccirillo, Anestis Touloumis, V Peter Collins, John C Marioni, Christina Curtis, Colin Watts, and Simon Tavaré. Intratumor heterogeneity in human glioblastoma reflects cancer evolutionary dynamics. Proceedings of the National Academy of Sciences, 110(10):4009–4014, 2013.

[53] Jack Edmonds. Optimum branchings. Mathematics and the Decision Sciences, Part, 1:335–345, 1968.

[54] Mohammed El-Kebir, Layla Oesper, Hannah Acheson-Field, and Benjamin J Raphael. Reconstruction of clonal trees and tumor composition from multi-sample sequencing data. Bioinformatics, 31(12):i62–70, June 2015.

[55] Victoria Popic, Raheleh Salari, Iman Hajirasouliha, Dorna Kashef-Haghighi, Robert B West, and Serafim Batzoglou. Fast and scalable inference of multi-sample cancer lineages. Genome Biology, 16(1):795–17, May 2015.

[56] C Chow and Cong Liu. Approximating discrete probability distributions with dependence trees. IEEE transactions on Information Theory, 14(3):462–467, 1968.

