## Supplementary Materials for "Learning mutational graphs of individual tumour evolution from single-cell and multi-region sequencing data"

Supplementary Material

Daniele Ramazzotti, Alex Graudenzi, Luca De Sano, Marco Antoniotti, Giulio Caravagna

**7 Case studies** **34**

**8 Noise model** **34**

**9 Modeling back mutations** **40**

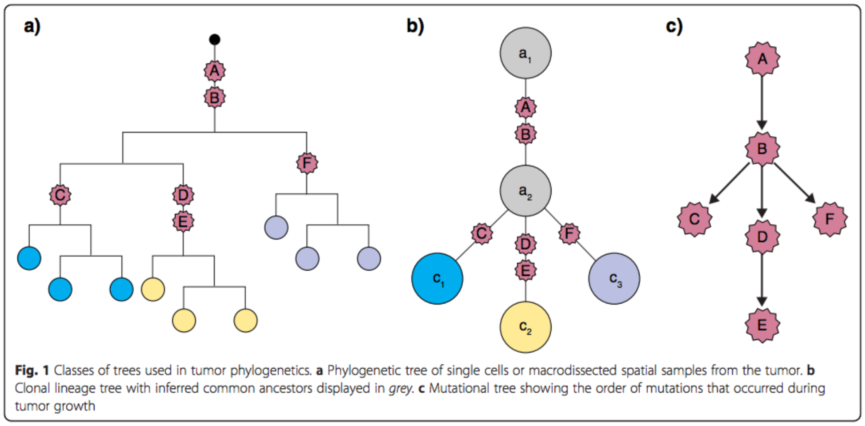

Supplementary Figure 1: The $\star$OP problems (image taken from [1]) as suggested by Davis and Navin, which define three ways to infer cancer progression from single-cell data. The POP (*phylogenetic ordering problem*), a classical phylogenetic inference problem where we display input cells as leaves of a phylogenetic tree. The COP (*clonal ordering problem*), where we identify a clonal lineage tree that models an ancestry-relation for a set of clones that we infer from data. The MOP (*mutational ordering problem*), where we find the order of mutations that accumulate during cancer progression. In this paper, we focus on the MOP problem.

### 1 A framework based on probabilistic causation

In Figure 1 of [1] Davis and Navin distinguish three different approaches to infer cancer progression models from data of individual patients. We summarize them in Supplementary Supplementary Figure 1 and call them $\star$OP, mimicking the *"$\star$-ordering problem"*. Different versions of the $\star$OP provide insights on the evolutionary aspects of cancer progression. In particular:

- when $\star$ = P, we solve a *classical phylogenetic inference problem* (PHY) and aim at displaying a set of input cells as leaves of a phylogenetic tree;
- when $\star$ = C, we seek to identify a *clonal lineage tree* (CT) that models an ancestry-relation for a set of clones that we infer from data;
- when $\star$ = M, we seek to find the order of mutations that accumulate during progression (MO).

Hopefully, results from these approaches can be somehow reconciled, as the same data type is used to approach the problem. For instance, we might argue that the order of accumulating mutations should be consistent with the clonal lineage tree, which in turn should be consistent with a phylogenetic tree of the corresponding cells that we sequence. The efforts to solve these problems are ongoing, with different techniques and tools that are becoming popular to solve the COP and the POP versions of the problem, see, e.g., [2–4]. We focus this paper on the mutational version of the problem, consistently with earlier works of us [5–10].

#### 1.1 Preliminaries

We use Suppes' framework of *probabilistic causation* as the core of our approach to cancer progression inference. We use it to derive algorithms that exploit optimal results from *minimum spanning tree reconstruction* and *Bayesian inference.* These axioms provide a *necessary but not sufficient* set of conditions to make causal claims [6–10]. In our earlier works [5–10] we considered data from multiple patients, i.e., multiple observations of the tumor progressions, to disentangle genuine from spurious causal relations. On the contrary, here we can quantify the statistical trends between mutations with Suppes' conditions, but we need to clarify that we are observing multiple measurements from the same patient (not across different patients). Thus, our claims can not be of a general causal nature, and we have to restrict to the estimation of the temporal progression in the individual tumor. This is also reflected in the spanning tree assumption of our algorithms, which implies that one unique predecessor is assigned to every considered mutation. For these reasons, the depicted relations are valid temporal orderings, even if they might depict spurious causal relations.

##### Input data

We consider a binary-valued dataset $\mathbf{D}$ with $n$ variables and $m$ observations

$$\begin{array}{c} \\ \mathbf{z}_1 \\ \mathbf{z}_2 \\ \vdots \\ \mathbf{z}_m \end{array} \begin{array}{c} \begin{array}{cccc} \mathbf{x}_1 & \mathbf{x}_2 & \cdots & \mathbf{x}_n \end{array} \\ \begin{pmatrix} x_{1,1} & x_{1,2} & \dots & x_{1,n} \\ x_{2,1} & x_{2,2} & \dots & x_{2,n} \\ \vdots & \vdots & \ddots & \vdots \\ x_{m,1} & x_{m,2} & \dots & x_{m,n} \end{pmatrix} \end{array} = \mathbf{D} \tag{1}$$

where the columns of $\mathbf{D}$ are the $n$ variables $\mathcal{X} = \{\mathbf{x}_1, \ldots, \mathbf{x}_n\}$, $\mathbf{x}_i \in \{0,1\}^m$, and $\mathbf{z}_1, \ldots, \mathbf{z}_m$ are the $m$ samples. Variables refer to genomic events detected by sequencing of cancer genomes (i.e., (epi)genomic alterations of various types such as, e.g., single nucleotide or structural variants, or copy number alterations). Value $x_{i,j} = 1$ means that event $\mathbf{x}_j$ is detected in sample $\mathbf{z}_i$.

*Assumptions.* We require each event $\mathbf{x}_i$ to measure a *somatic alteration that is persistent* during tumor evolution, e.g., a mutation or a copy number variant. Epigenetic states of expression or methylation could be used only if they fulfil this condition; this is to be verified and assessed outside of our framework. A further technical assumption, not motivated by the phenomenon of cancer progression, is that no columns of $\mathbf{D}$ are either all zeros, or ones, and that no two columns exist that are indistinguishable. For this reason in our implementation $\mathbf{D}$ is reshaped appropriately before applying our algorithms, as we discuss in the Main Text.

##### Output model

From $\mathbf{D}$, we want to estimate a *joint distribution* $p(\cdot)$ over $\mathcal{X}$ so that we can sample genotypes from our output model. In our formulation we use ideas from Bayesian Networks [11], a class of Graphical Models based on *directed acyclic graphs* augmented with parameters $\boldsymbol{\theta}$.

A partially order set (poset) of a graph $\mathcal{G}$ over $\mathcal{X}$ is defined by a *partial ordering* $\sqsubseteq$: if $\mathbf{x}_i \sqsubseteq \mathbf{x}_j$ than edge $\mathbf{x}_i \to \mathbf{x}_j$ is in $\mathcal{G}$. For $\mathcal{G}$ to be acyclic we also require the *transitive closure* of $\sqsubseteq$ to have no path that start and end in the same $\mathbf{x}_i$. We turn $\mathcal{G}$ into a generator for a *conditional probability* *distribution* by defining the parameters

$$\boldsymbol{\theta}_{\mathbf{x}_j} = \hat{p}(\mathbf{x}_j \mid \{\mathbf{x}_i \mid \mathbf{x}_i \sqsubseteq \mathbf{x}_j\}) \tag{2}$$

where $\hat{p}$ is a distribution conditioned to the incoming edges of $\mathbf{x}_j$, often called *conditional probability table* [11]. $|\boldsymbol{\theta}_{\mathbf{x}_j}|$ is exponential in the number of edges incoming to $\mathbf{x}_j$, if variables are binary.

When we infer a model of cancer progression for a patient, our output model will be the most likely *tree* (but in some cases it could be a *forest*, or a general graph), according to a measure of ti. A graph $\mathcal{G}$ is a tree if ($i$) it has one *root node* $\mathbf{x}_*$ with no incoming edges, and ($ii$) all other $\mathbf{x}_j \neq \mathbf{x}_*$ have one incoming edge, i.e., $|\{\mathbf{x}_i \mid \mathbf{x}_i \sqsubseteq \mathbf{x}_j\}| = 1$. Hence, for a tree, the parameters also simplify to $\boldsymbol{\theta}_{\mathbf{x}_j} = \hat{p}(\mathbf{x}_j \mid \mathbf{x}_i)$, i.e., they become linear in size. A graph $\mathcal{G}$ is a *forest*, if it can be partitioned into a set of trees. Trees and forests are acyclic, by definition.

#### 1.2 Weighted graphs with information theory

**Suppes' conditions as prior graph structure**

We frame Suppes' *probabilistic causation* [12] within cancer progression [5, 6], to create a *partial ordering* $\sqsubseteq_{\mathsf{PF}}$ over $\mathcal{X}$. We dub it *prima facie*, and we will use it to create the final output model's structure $\sqsubseteq$. For any pair of variables $\mathbf{x}_i$ and $\mathbf{x}_j$, we define

$$\mathbf{x}_i \sqsubseteq_{\mathsf{PF}} \mathbf{x}_j \iff p(\mathbf{x}_i) > p(\mathbf{x}_j) \wedge p(\mathbf{x}_j \mid \mathbf{x}_i) > p(\mathbf{x}_j \mid \overline{\mathbf{x}}_i)\,. \tag{3}$$

In general, $\sqsubseteq_{\mathsf{PF}}$ induces a *cyclic graph*. We interpret prima facie as a necessary condition for cancer progression, along the lines of [6]. So, we consider $\sqsubseteq_{\mathsf{PF}}$ to provide us with a superset of the edges that will appear in our output models; derivation of such edges is discussed in the next sections.

To include a pair of variables in $\sqsubseteq_{\mathsf{PF}}$, we test two inequalities over distributions estimated from $\mathbf{D}$. A statistical model of those marginal and joint/ conditional distributions over $\mathcal{X}$ can be created via *non-parametric bootstrap* [6]. Then, we can carry out a Mann-Whitney U test to compute a p-value for the alternative hypothesis that the distributions have different means: $\mathbf{x}_i \sqsubseteq_{\mathsf{PF}} \mathbf{x}_j$ when both inequalities have confidence below some desired p-value (e.g., $p < 0.05$). This testing/ bootstrap schema can support prior information of noise in the data. If we are informed that $\mathbf{D}$ harbours *false positives and negatives* rates $\epsilon_+$ and $\epsilon_-$, we can correct the marginal bootstrap estimates as

$$p(\mathbf{x}_i) = \frac{n_i - \epsilon_+}{1 - \epsilon_+ - \epsilon_-}\,, \tag{4}$$

where $n_i = \dfrac{\sum_k x_{k,i}}{m}$, and proceed similarly for the joint estimates as follows

$$p(\mathbf{x}_{i,j}) = \frac{n_{i,j} - \epsilon_+[n_i + n_j - \epsilon_+]}{(1 - \epsilon_+ - \epsilon_-)^2}\,, \tag{5}$$

where $n_{i,j} = \dfrac{\sum_k x_{k,i} x_{k,j}}{m}$. See Section 8 for the complete derivation of the error model.

##### Information-theoretic measures for associations' detection

The ordering $\sqsubseteq_{\mathsf{PF}}$ is a super set of the ordering that we want to return as output; we thus need to subset $\sqsubseteq_{\mathsf{PF}}$. To rank and select pairs in $\sqsubseteq_{\mathsf{PF}}$ we can use a score function. If we interpret each $\mathbf{x}_i$ as a random variable with binary outcomes, we can compute information-theoretic measures for the detection of its degree of association to other variables [13]. For each $\mathbf{x}_i \sqsubseteq_{\mathsf{PF}} \mathbf{x}_j$, we measure the *point-wise mutual-information* (pmi)

$$\mathsf{pmi}(\mathbf{x}_i = x, \mathbf{x}_j = y) = \log \left[ \frac{p(\mathbf{x}_i = x, \mathbf{x}_j = y)}{p(\mathbf{x}_i = x) p(\mathbf{x}_j = y)} \right], \tag{6}$$

that quantifies the discrepancy between $\mathbf{x}_i$ and $\mathbf{x}_j$ for their outcomes $x$ and $y$. Here, to detect the association between alterations that accumulate during progression, we set $x = y = 1$.

In some cases, we will also use the expected value of pmi over all the possible outcomes of $\mathbf{x}_i$ and $\mathbf{x}_j$, which is the *mutual information* (mi)

$$\mathsf{mi}(\mathbf{x}_i, \mathbf{x}_j) = \sum_{x,y} p(\mathbf{x}_i = x, \mathbf{x}_j = y) \mathsf{pmi}(\mathbf{x}_i = x, \mathbf{x}_j = y) . \tag{7}$$

These measures are standard [13], and could be used to derive alternative score functions (e.g., conditional pointwise or entropy). In this work, however, we limit our scope to pmi and pmi.

#### 1.3 Strategies for structure selection and parameters' learning

The prima facie ordering $\sqsubseteq_{\mathsf{PF}}$ induces a mapping to $2^{|\sqsubseteq_{\mathsf{PF}}|}$ potential models. Some of these are not trees or might model the distribution of the data poorly. The problem of picking a particular $\sqsubseteq$ to build $\mathcal{G}$ is hence non-trivial.

By combining $\sqsubseteq_{\mathsf{PF}}$ with a pmi/mi score we have obtained a *weighted graph.* Thus, we can exploit algorithms that extract trees (or other types of models) with certain properties, from the input graph. We denote with $\sqsubseteq_{\mathsf{PF}}^{\mathsf{pmi}}$ and respectively with $\sqsubseteq_{\mathsf{PF}}^{\mathsf{mi}}$ the graphs weighted with pmi or mi. We group a number of algorithms for structure selection into two classes: *(minimum) spanning tree algorithms* and *Bayesian model selection methods.*

##### Minimum spanning tree algorithms

These are a class of algorithms that aim at detecting the subset of pairs $\sqsubseteq$ that $(i)$ minimize the total output structure's weight, and that $(ii)$ display as a tree. The total weight is defined as the summation of the weights of the pairs that are selected. This is a well-known problem in graph theory, and we can exploit optimal solutions from the literature. The approaches are different according to the graph that these algorithms are given as input.

We note that using "spanning tree" algorithms is a standard technique in the field [14,15]. Our approach is not different in spirit, while we reuse these algorithms with a different weight structure induced by Suppes' causal ordering $\sqsubseteq_{\mathsf{PF}}$.

**Edmond:** Edmonds' optimum branching algorithm is a canonical solution to the task of inferring a spanning tree of minimum weight from a weighted directed graph, given a root node in input [16]. We use this algorithm with $\sqsubseteq_{\mathsf{PF}}^{\mathsf{pmi}}$ as input, *provided that we make it acyclic.* Cycles/ loops breaking is a hard computational problem, and we resort on the heuristic that

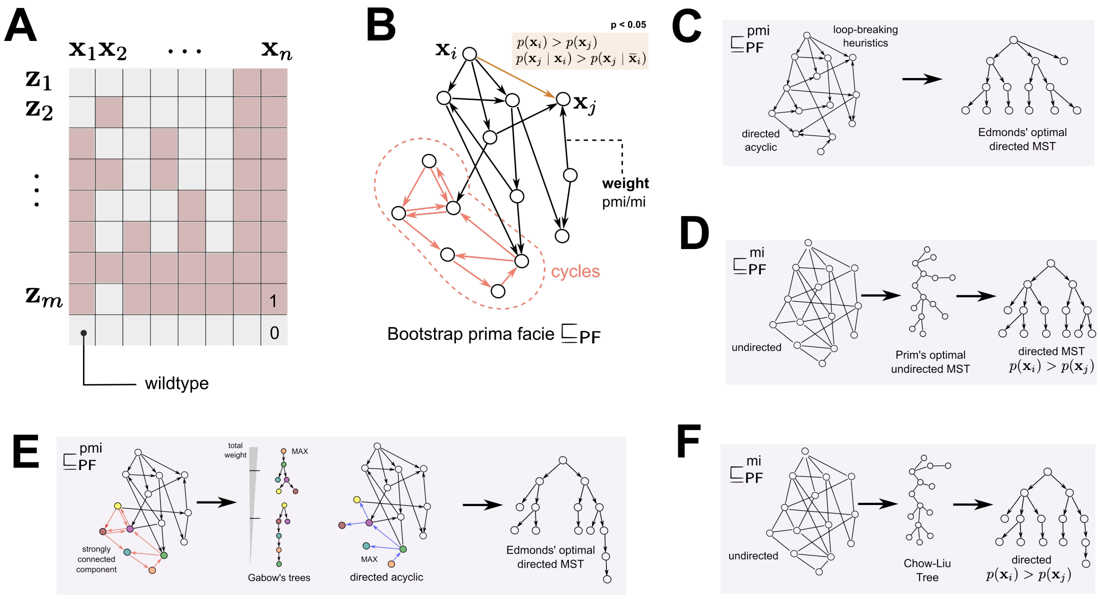

Supplementary Figure 2: **A**. The input data is a binary matrix that store the presence/absence of a variable in a sample. Some of this observations will be due to noise as false positives and negatives. **B**. We estimate via bootstrap the prima facie ordering relation $\sqsubseteq_{\mathsf{PF}}$ that satisfies Suppes' conditions for probabilistic causation. This, in turn, induces a graph over variables $\mathcal{X}$. This graph can be weighted by measuring information-theoretic measures for variables' association. **C**. If we weighted the graph with pmi, we can use an heuristic to make it acyclic, and use Edmonds' optimal solution for minimum directed spanning tree detection. **D**. If we weighted the graph with mi, we can disregard the edges' orientation, and use Prim's optimal solution for minimum undirected spanning tree detection. Then, we can orient by the marginals each edge. **E**. If we weighted the graph with pmi, we can use Gabow's optimal solution for path traversals of cyclic component and detect the best tree that makes the structure acyclic. Then, we can again use Edmonds' algorithm for spanning trees. **F**. A Bayesian optimal mode-selection strategy can compute the Chow-Liu tree that induces the distribution with minimum divergence from the true one. In this case, we weighted the graph with mi and made it undirected.

is used by the CAPRI algorithm [6]. This heuristics breaks loops according their confidence, defined as the combination of *p-values* for Suppes' conditions: edges with small confidence (i.e., high p-values) are deleted first to break loops. When $\sqsubseteq_{\mathsf{PF}}^{\mathsf{pmi}}$ is made acyclic, to use Edmonds' algorithm and maximize our pmi scores, we can change their sign. The running time of this algorithm is $\mathcal{O}(|\mathcal{X}| \cdot |\sqsubseteq_{\mathsf{PF}}^{\mathsf{pmi}}|)$, which can be optimized to $\mathcal{O}(|\mathcal{X}|\log(|\sqsubseteq_{\mathsf{PF}}|))$ for sparse[1] $\sqsubseteq_{\mathsf{PF}}^{\mathsf{pmi}}$ [17].

[1] An ordering relation is sparse if its associated matrix over $\{0,1\}^{\mathcal{X}\times\mathcal{X}}$ is sparse.

**Prim:** Prim's algorithm is the equivalent of Edmonds' for undirected tree structures [18]. We can use Prim's algorithms by disregarding the directionality of the edges in our prima facie ordering, i.e., if for every pair $\mathbf{x}_i \sqsubseteq_{\mathsf{PF}} \mathbf{x}_j$ we also force the inclusion of $\mathbf{x}_j \sqsubseteq_{\mathsf{PF}} \mathbf{x}_i$ no matter what the test statistics for Suppes' conditions. If we want to use this search strategy, however, we will use $\sqsubseteq_{\mathsf{PF}}^{\mathsf{mi}}$ as input, as we need to use a measure that is symmetric and defined over all the support of the random variables (i.e., pmi is not accounting for the cases $\mathbf{x}_i = 1 - \mathbf{x}_j = 1$ and viceversa). The complexity of this algorithm, if it is implemented by using a binary heap and an adjacency list for $\mathcal{G}$, is the same as Edmonds'. The final tree returned by this strategy is undirected, and so we orient it according to the marginal frequencies of the events. That is, for every final pair $\mathbf{x}_i \sqsubseteq_{\mathsf{PF}} \mathbf{x}_j$ and $\mathbf{x}_j \sqsubseteq_{\mathsf{PF}} \mathbf{x}_i$ we select $\mathbf{x}_i \to \mathbf{x}_j$ if $p(\mathbf{x}_i) > p(\mathbf{x}_j)$. For this reason, the final model could contain confluent structures such as $\mathbf{x}_i \sqsubseteq_{\mathsf{PF}} \mathbf{x}_j$ *and* $\mathbf{x}_k \sqsubseteq_{\mathsf{PF}} \mathbf{x}_j$ – i.e., a model with two edges confluent in $\mathbf{x}_j$: $\mathbf{x}_i \to \mathbf{x}_j$ *and* $\mathbf{x}_k \to \mathbf{x}_j$. We observe that, when this happen, the final model is not a tree as $\mathbf{x}_j$ has more than one parent, but a multi-rooted directed acyclic graph (DAG). The interpretation in terms of the induced distribution is still that of a Bayesian Network.

**Gabow:** Cycles in $\sqsubseteq_{\mathsf{PF}}^{\mathsf{pmi}}$ can be handled in another search schema by exploiting Gabow's algorithm [19], before using Edmond's algorithm to maximize the weight of the final tree. Gabow's algorithm algorithm is optimal to detect the *strongly connected components* of the directed graph $\sqsubseteq_{\mathsf{PF}}^{\mathsf{pmi}}$ in time $\mathcal{O}(|\mathcal{X}| + |\sqsubseteq_{\mathsf{PF}}|)$ if the graph is represented as an adjacency list. If $\sqsubseteq_{\mathsf{PF}}^{\mathsf{pmi}}$ has cycles, we thus create, for each strongly connected component, all the possible trees associated; then we select the tree at maximum weight for each such set of candidates. We can optimize this procedure by separating acyclic sub-graphs, if any, in the very beginning. This algorithm is optimal but more expensive than Edmond, and shall be seen as an alternative way to deal with loops in the prima facie structure.

##### Bayesian model-selection strategies

**Chow-Liu:** This is an optimal method for constructing a *second-order product approximation* of the joint probability distribution over $\mathcal{X}$ [20]. It is known that the resulting tree minimizes the Kullback-Leibler distance to the actual joint distribution, and can be interpreted as a Bayesian Network. For constructing the optimal tree, at each iteration the algorithm adds the maximum mi pair to the tree. This algorithm returns an undirected structure when we run it with $\sqsubseteq_{\mathsf{PF}}^{\mathsf{mi}}$ as input; we transform it into a directed structure as we do with the Prim search strategy. The similarity between the two algorithms is evident, also in terms of complexity.

##### Learning parameters $\boldsymbol{\theta}$

Given a graph (or, as a special case a tree), we can fit its parameters using a standard technique in the Bayesian Networks approach, by maximum likelihood estimation from $\mathbf{D}$ [11].

##### Comparison with other algorithms

As motivated in the Main Text, SCITE and OncoNEM are at the state-of-the art for two orthogonal problems in single-cell phylogenetic inference (mutational vs clonal ordering). In our simulations we compared the different model search methods just listed against SCITE, since its mutational tree is directly comparable to our models.

We note that we were not able to test all the simulation scenarios we created with OncoNEM, as, at the time of testing, its performance scaled poorly to carry out our large scale test. See for instance Supplementary Supplementary Table 6 with some example timings to run these comparisons.

### 2 Testing the framework

#### 2.1 Synthetic data and performance measures

Comparison among the algorithms is based on large synthetic tests for different combinations of model type, size, number of samples, noise etc. We describe here the details of the approach, and provide the user with its R implementation in the TRONCO tool which is available at

$\texttt{https : //sites.google.com/site/troncopackage}$

and on Bioconductor.

We devised a testing framework to gather information about the relative performance of TRaIT in a number of different scenarios.

1. Sampling from *Single-cells* (SCs) sequencing data.
2. Sampling from *Multi-region bulk* sequencing data.

In each case we take care to explore the problems induced by the presence of noise in the data.

##### Sampling from Single-cell sequencing

Genotypes from single-cell sequencing are sampled by a phylogeny. We describe the simpler case of sampling from a single tree, more general cases are trivial extensions. A cartoon is shown in Supplementary Figure 3 that shows some possible single-cell genotypes.

The following recursive procedure visits a tree, starting from its root $x_*$ (i.e., we set $\mathbf{x}_* = 1$ in the genotype), and outputs a sampled genotype according to its structure and parameters.

- If we are visiting a leaf $\mathbf{x}_l$ (i.e., a node without outgoing edges) with incoming edge $\mathbf{x}_i \to \mathbf{x}_l$, then we sample $\mathbf{x}_l = 1$ (in the genotype) with probability $\boldsymbol{\theta}_{\mathbf{x}_l} = p(\mathbf{x}_l \mid \mathbf{x}_i)$, and 0 otherwise.
- if we are visiting a branching node $\mathbf{x}_i$ (i.e., $\mathbf{x}_i = 1$ in the genotype) with children $\mathbf{x}_i \to \mathbf{x}_j$ and $\mathbf{x}_i \to \mathbf{x}_k$ we either sample only one of the children and we proceed recursively, or we stop. Notice that we forbid to sample a genotype with both children[2], i.e., $p(\mathbf{x}_j, \mathbf{x}_k \mid \mathbf{x}_i) = 0$, so

$$\begin{aligned} p(\mathbf{x}_j \mid \mathbf{x}_i) &= p(\mathbf{x}_j, \overline{\mathbf{x}}_k \mid \mathbf{x}_i) + p(\mathbf{x}_j, \mathbf{x}_k \mid \mathbf{x}_i) = p(\mathbf{x}_j, \overline{\mathbf{x}}_k \mid \mathbf{x}_i) \\ p(\mathbf{x}_k \mid \mathbf{x}_i) &= p(\overline{\mathbf{x}}_j, \mathbf{x}_k \mid \mathbf{x}_i) + p(\mathbf{x}_j, \mathbf{x}_k \mid \mathbf{x}_i) = p(\overline{\mathbf{x}}_j, \mathbf{x}_k \mid \mathbf{x}_i) \,. \end{aligned} \tag{8}$$

  Thus, genotype with $\mathbf{x}_i = \mathbf{x}_j = 1 - \mathbf{x}_k$ has probability $\boldsymbol{\theta}_{\mathbf{x}_j} = p(\mathbf{x}_j \mid \mathbf{x}_i)$, while genotype with $\mathbf{x}_i = \mathbf{x}_k = 1 - \mathbf{x}_j$ has probability $\boldsymbol{\theta}_{\mathbf{x}_k} = p(\mathbf{x}_k \mid \mathbf{x}_i)$ and genotype $\mathbf{x}_i = 1 - \mathbf{x}_j = 1 - \mathbf{x}_k$ has probability $1 - [p(\mathbf{x}_j \mid \mathbf{x}_i) + p(\mathbf{x}_k \mid \mathbf{x}_i)]$. To have consistent cell genotypes for the whole model, when we recursively proceed with $\mathbf{x}_j$ (resp. $\mathbf{x}_k$) we set $\mathbf{x}_k$ (resp. $\mathbf{x}_j$) and all its descendants equal to 0.

[2] This is analogous of saying that, at the genotype level, every branching is interpreted as an *exclusive branch*. In this case the truth table for $\mathbf{x}_i$, $\mathbf{x}_k$ and $\mathbf{x}_k$ resembles a xor-network when $\mathbf{x}_i = 1$.

Notice that, by construction, genotypes are consistent with the phylogeny of the generative model, as $p(\mathbf{x}_j, \mathbf{x}_k \mid \overline{\mathbf{x}}_i) = p(\overline{\mathbf{x}}_j, \mathbf{x}_k \mid \overline{\mathbf{x}}_i) = p(\mathbf{x}_j, \overline{\mathbf{x}}_k \mid \overline{\mathbf{x}}_i) = 0$ for any branching structure.

We observe that: (*i*) we can easily generalize this procedure to an arbitrary amount of children per branching, and that (*ii*) this procedure generates only genotypes that correspond to cancer cells (because we start with $\mathbf{x}_* = 1$). If required, we can a posteriori add wild-type genotypes to a dataset to account for contamination of normal cells.

##### Sampling from Multi-region bulk sequencing

When we collect and sequence a bulk of tumor cells we get a signal that is a mixture of alterations found in different tumor sub-populations.

In terms of induced distribution and the branching structures described in single-cell sequencing sampling, this means that data will support $p(\mathbf{x}_i = \mathbf{x}_j = \mathbf{x}_k) > 0$ as the sequenced samples will contain cells from both populations with signatures $\mathbf{x}_i = \mathbf{x}_j = 1 - \mathbf{x}_k$ and $\mathbf{x}_i = \mathbf{x}_k = 1 - \mathbf{x}_j$. To create such a signal there are different ways. On one hand, one can change the effect of branchings on the induced distribution to account for $p(\mathbf{x}_i = \mathbf{x}_j = \mathbf{x}_k) > 0$, on the other one can emulate a mixed signal by mixing a number of individual signals. The former approach requires more parameters in the generative model to account for the conditional probabilities of both children, given a parent node. We adopt the latter approach and sample $c$ genotypes from a single-cell sequencing experiment: let $\mathbf{z}_1, \ldots, \mathbf{z}_c$ be such samples, we create a sample

$$\mathbf{z}_* = \bigvee_i \mathbf{z}_i \tag{9}$$

where each component of $\mathbf{z}_*$ is 1 if at least one $\mathbf{z}_i$ is 1. Then, we repeat the procedure to produce as many samples as we need, according to the number of regions that we want to simulate.

This approach requires only one more parameter, $c$. If one interprets the $c$ samples as $c$ cells, one might be tempted to pick a very large $c$ (e.g., $c \gg 10^6$). If one does so, however, all $\mathbf{z}_*$ will be similar, as for large $c$ the proportions of the sampled genotypes will converge to the true ones. Thus, our dataset would have small variance across samples, biasing the data. Thus, to have more variance, we set $c$ to be small ($c < n$) and interpret $c$ as the probabilistic number of clones spread across the regions that we sequence.

##### Adding noise to synthetic data

When we say that observed data are obtained by adding noise to sampled genotypes, we mean the usual introduction of *false positives and negatives* with rates $\epsilon_+ \geq 0$ and $\epsilon_- \geq 0$, respectively [2,3,6]. Precisely, when we have sampled a dataset of samples $D$ according either to (8) or (9), we apply to $D$ an independent point-wise process that flips the matrix's entries according to the rates $\epsilon_+/\epsilon_-$.

To investigate the ideal performance of the algorithms we sometimes use *noise-free* data, that is $\epsilon_+ = \epsilon_- = 0$. In more realistic setting, we use different models of noise according to the type of simulate sequencing. Sequencing of single cells is characterized by distinct errors, which usually take place in the DNA amplification phase: (*i*) *allelic dropouts* and (*ii*) *false alleles*, the former occurring at a significantly higher rate, thus leading to higher rates of false negatives. Accordingly, in the generation of noisy single-cell data we expect highly asymmetric noise parameters $\epsilon_+ \ll \epsilon_-$, that we simulate by assuming one-order of magnitude in their difference. Multi-region bulk sequencing data instead harbour more balanced noise effects, in that case we set $\epsilon_+ = \epsilon_-$.

**Performance measured**

We want to measure the tendency to overfit or underfit of every algorithm, in particular circumstances of sample size, noise etc. Thus, the performances measured in each experiment are:

- the rate at which *true model edges are inferred*

$$\textit{sensitivity} \quad \frac{\mathsf{TP}}{\mathsf{TP}+\mathsf{FN}};$$

- the rate at which *false model edges are discarded*

$$\textit{specificity} \quad \frac{\mathsf{TN}}{\mathsf{TN}+\mathsf{FP}};$$

where TP, TN, FP, FN are the number of true (T)/ false (F) positives (P) and negatives (N).

**Algorithms' implementation**

We used the official release of each tool.

- CAPRI, CAPRESE and the algorithms that we describe in this paper are available in TRONCO.
- SCITE was downloaded from its Github repository.
- OncoNEM was downloaded from its Bitbucket repository.

#### 2.2 Working scenarios

We define four possible working scenarios, which represent distinct cancer evolution *modes* and related phenomena (Supplementary Supplementary Figure 3).

**Branching evolution**
In this case different cancer subclones (with distinct lineages) diverge from a common ancestor and are characterized by distinct accumulating alteration. This can be modelled via trees with distinct branches describing the subclonal trajectories. Particular instances of this scenario is *linear evolution*, in which alterations accumulate along a linear path, with no branches.

**Confounding factors**
In this case the generative models are phylogenetic trees, as above, but the observed data also include *uncorrelated random events*. This is a handle to account for possible *confounding factors*, i.e., (epi)genomic alterations that have no functional role in the progression, and that we do not know a priori.

**Multiple independent trajectories**
In this case the generative topology are multiple independent trees, grouped as a *forest*. This is a way to model tumors that originate from two or more cells, a phenomenon also known as *polyclonal tumor origin* [21], or the possible presence of *hidden* events triggering tumor development, but not annotated in the available data (e.g., methylations).

According these working scenarios, we perform a simple non-exhaustive test of the performance of the algorithms, with single-cell and multi-region data.

### 3 Preliminary tests

These tests are rather simple in size and variability of settings, and more detailed tests are carried out in the next sections. However, they still give a general idea of the general performance trend.

#### 3.1 Single-cell data

**Experiment I.**

We test a set of fixed topologies according to the scenarios in Supplementary Figure 3: (*i*) a tree with $n = 11$ nodes (*Branching evolution*), (*ii*) which we then augment with the addition of 2 disconnected nodes (*Confounding factors*), (*iii*) a forest with two distinct trees that account for $n = 7$ nodes (*Multiple independent trajectories*). . For each single test we generated 100 single-cell datasets with $m = 75$ samples and a mild noise setting, $\epsilon_+ = 0.005$ $\epsilon_- = 0.05$.

**Results: (Supplementary Figures 4, 5; Supplementary Tables 1, 2,**

1. *Branching evolution*

   The performances of all the algorithms are consistently similar and very good. Yet, Gabow and Edmond with pmi reach highest performance, while SCITE displays a larger dispersion. All algorithm but CAPRI have similar median sensitivity, suggesting a comparable capability of inferring the true relations from data. Specificity scores, instead, suggest that SCITE tends to overfit more than other approaches (i.e., lower true negative rate). CAPRI displays a very high specificity, but that is possibly due its "regularization" approach that tends to return sparse models (thus, trading specificity for sensitivity)[3].

2. *Confounding factors*

   A similar trend is observed also in this scenario, with SCITE displaying a very good sensitivity, but also the lowest specificity among all. We can observe that the best algorithm in this case seems to be Prim, as it displays the same sensitivity of SCITE, but higher specificity. CAPRI's very high specificity is still due to the regularization terms. We finally show in Supplementary Table 1 that our approach is capable to model these progressions, with the confounding factor consistently presenting a much lower trend of significance, compared to all the other events.

3. *Multiple independent trajectories*

   In this case Gabow, Edmond and SCITE present an identical performance with median values of both specificity and sensitivity equal to 100%, slightly outperforming the other algorithms. In particular, by looking at Supplementary Table 2 one can notice that all the techniques are able to retrieve the two distinct roots of the progression, with the exception of PRIM and CAPRESE in a few cases.

[3]This is a general trend of this algorithm that we expect to observe in all the experiments. Thus, we omit from commenting it any further.

Example genotypes sampled from the models below

**A** *single-cell*: $x_1x_2x_3$, $x_1x_5$, $x_1x_5x_6$
*multi-region*: $x_1x_2x_3$, $x_1x_2x_5$ (e.g., $x_1x_2/x_1x_5$), $x_1x_5x_6x_2x_4$ (e.g., $x_1x_2x_4/x_1x_5x_6$)

**B** *single-cell*: $x_1x_2x_3\underline{z_1}\underline{z_2}$, $x_1x_5\underline{z_2}$, $x_1x_5x_6$
*multi-region*: $x_1x_2x_5\underline{z_1}\underline{z_2}$ (e.g., $x_1x_2\underline{z_1}/x_1x_5\underline{z_2}$ or $x_1x_2\underline{z_1}\underline{z_2}/x_1x_5$ or ...)

**C** *single-cell*: $x_1x_2x_3$, $x_7x_8$, $x_1$, $x_7x_8x_9$
*multi-region*: $x_1x_2x_5$ (e.g., $x_1x_2/x_1x_5$), $x_1x_2x_5x_7x_8$ (e.g., $x_1x_2/x_1x_5/x_7x_8$)

Observed data obtained by applying to noise to genotypes

In **A**, the single-cell genotype $x_1x_2x_3$ might be observed as $x_1x_3$, when $x_2$ is not detected (false negative), or as $x_1x_2x_3x_4$ when $x_4$ is wrongly detected (false positive).

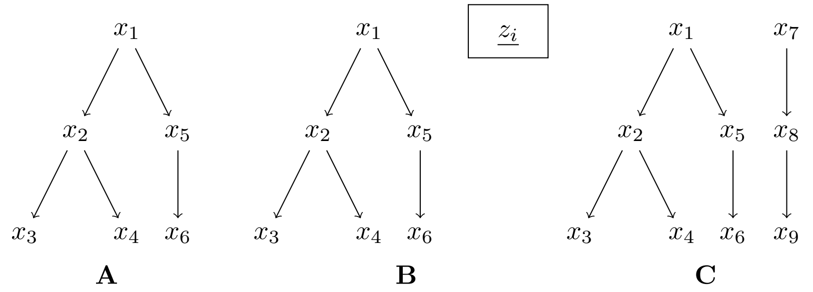

Supplementary Figure 3: **A.** Branching evolution. This phylogenetic tree has 6 nodes (one for each alteration $x_i$), and a unique variant/cell of origin, $x_1$. This is the most common case in which inference is carried out. **B.** Confounding factors. A phylogenetic model can be extended with spurious variables $\underline{z_i}$ that confound the inference problem. The inference is hindered because we have to detect the spuriousness of the association between any $\underline{z_i}$ and the true variants $x_i$. **C.** Multiple independent trajectories.A forest of phylogenetic models can describe the presence of multiple independent progressions that start from different variants ($x_1$ and $x_7$), as it might happen with tumors that start from different cells of origin, or with hidden/ not-annotated events triggering tumor development.

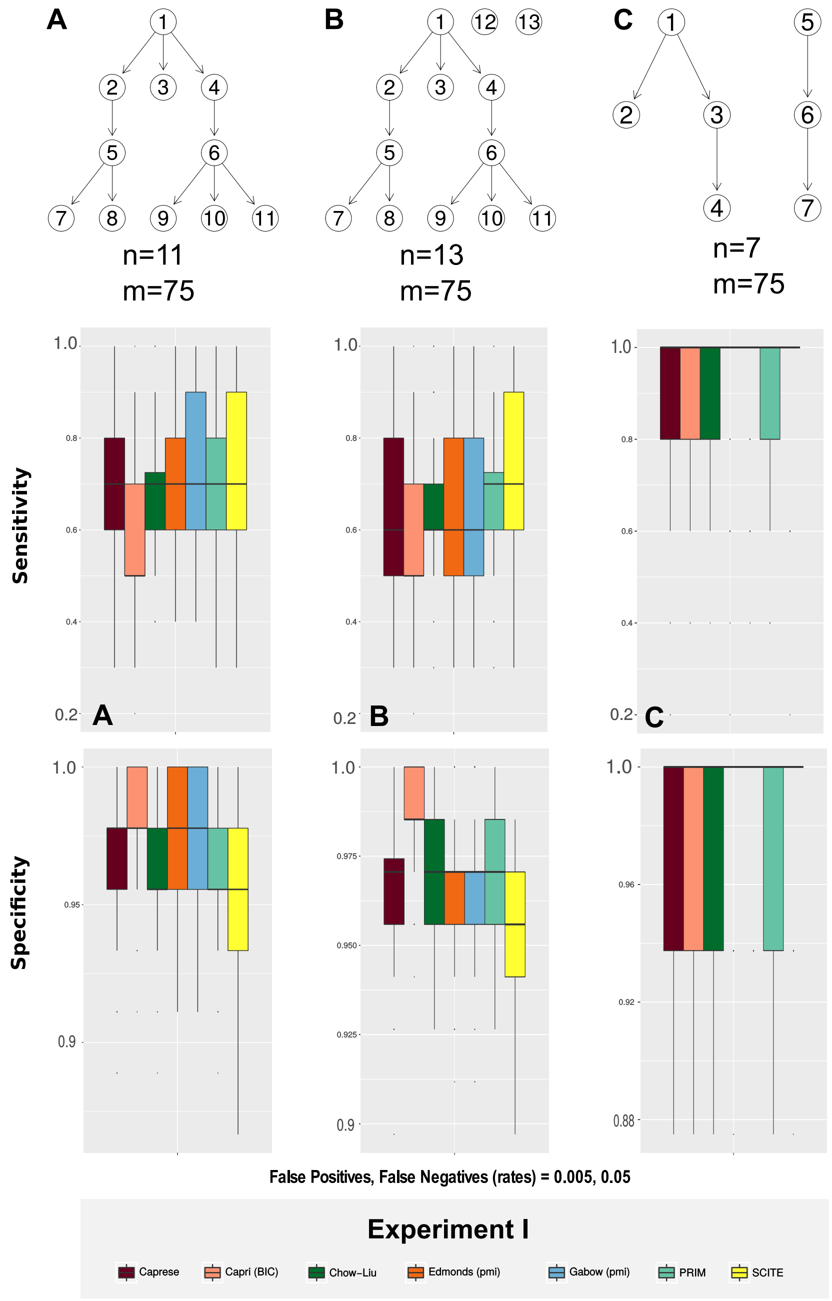

Supplementary Figure 4: Experiment I.(SCS data). We test the CAPRESE ($\lambda = 0.5$), CAPRI (with BIC regularization) and SCITE (default parameters) against our new algorithms, for model inference from single-cell data with noise $\epsilon_+/\epsilon_- = 0.005/0.05$. We use Gabow and Edmond with pointwise mutual information (pmi). The boxplots present sensitivity and specificity scores for 100 distinct datasets representing the working scenarios discussed in Section 2.2. Results are discussed in Section 2.2. Parameters are reported in Section 6. **A.** Branching evolution. **B.** Confounding factors. **C**. Multiple independent trajectories.

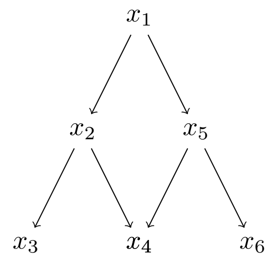

Supplementary Figure 5: **Statistical complications due to convergent trajectories.** Example of evolution in a single patient with convergent trajectories; this model does not fulfill the Infinite Sites Assumption (ISA) model. This specific case shows possible statistical issues complicating the inference, and limitations of our approach. Specifically, let us focus on a subset of 6 possible genotypes derived from the generative model above: $x_1x_2$, $x_1x_2x_3$, $x_1x_2x_4$, $x_1x_5$, $x_1x_5x_4$ and $x_1x_5x_6$. We now focus on the nodes presenting confluent trajectories, i.e., $x_2 \to x_4$ and $x_5 \to x_4$, and consider the probabilities involved in estimating the prior graph structure for these relations. Consider a fictitious dataset with 10 samples and assume that: $(i)$ genotypes $x_1x_2x_3$ and $x_1x_5x_6$ are very rare and never observed in our dataset, (i.e., with probability 0); $(ii)$ genotypes $x_1x_2x_4$ and $x_1x_5x_4$ have 4 times the probability of being observed than genotypes $x_1x_2$ and $x_1x_5$, (i.e., the former with probability 0.4 and the latter 0.1). We compute the marginal and joint probabilities for all the variables of interest: $p(x_2) = p(x_5) = 0.5$, $p(x_4) = 0.8$, $p(x_2, x_4) = p(x_4, x_5) = 0.4$. We observe that in this scenario Suppes' temporal priority is reverted; in fact, being $p(x_4) = 0.8 > 0.5 = p(x_2) = p(x_5)$, $x_4$ is estimated to be earlier in time than $x_2$ and $x_5$. At the same time, the 3 events present perfect independence – in fact, $p(x_2, x_4) = p(x_4, x_5) = p(x_2) \cdot p(x_4) = p(x_4) \cdot p(x_5) = 0.4$ – and for this reason also the probability raising condition would be violated. Although what shown here is a pretty rare and unfortunate configuration, we still point out that convergent trajectories especially in the case of hard exclusivity among the parents (such as this one), may further complicate the inference.

| | p-value poset | p-value (Edmond) | Edges (Edmond) |
|---|---|---|---|
| Node 1 | 0.05 | 1.20e-04 | 1.00 |
| Node 2 | 0.09 | 1.00e-02 | 2.00 |
| Node 3 | 0.12 | 5.06e-07 | 2.00 |
| Node 4 | 0.13 | 2.37e-06 | 1.17 |
| Node 5 | 0.11 | 2.00e-02 | 1.00 |
| Node 6 | 0.12 | 5.25e-06 | 1.01 |
| Node 7 (confounding) | **0.44** | **4.00e-01** | **0.77** |

Supplementary Table 1: Experiment I (confounding factors.) Mean p-values for probability raising, for each in/out-coming arcs in each node of Suppes' poset, and in the topology inferred by Edmond, for which we show the average number of arcs per node. The statistics are averaged over 100 SCS datasets generated from the low polyclonal tree topology with 1 confounding factor and $n = 7$ nodes (Supplementary Figure 7). We used a noise-free configuration and 100 samples.

#### 3.2 Multi-region bulk sequencing data

##### Experiment I-MR

*We reproduce* Experiment I *with multi-region data sampled from the same generative topologies of Supplementary Figure 4. The parameters are identical to* Experiment I, *with the exception of the number of samples (here biopsies, rather than cells), which is set to* $m = 20$.

**Results: (Supplementary Figure 6, Supplementary Tables 3)**

1. *Branching evolution.* All the algorithms but CAPRI and SCITE display identical median values of sensitivity and specificity, with Gabow slightly outperforming other techniques. However, all the algorithms struggle in retrieving a large number of true relations, as the median sensitivity ranges around 50% for most techniques. In this case, SCITE is the least accurate algorithm, showing a poor efficacy in retrieving both true positives and true negatives, whereas CAPRI displays high specificity and low sensitivity.

2. *Confounding factors.* The overall performance slightly worsens with confounders. In particular, the average sensitivity values are lower because the confounders introduces spurious correlations. The general trend is however preserved, with Gabow and SCITE being the most and least accurate algorithms.

3. *Multiple independent trajectories.* CAPRI, Prim and Chow-Liu show a slightly better trade-off between sensitivity and specificity, while SCITE is less accurate with this mixed signal. More in detail, most algorithms are unable to infer the two distinct roots of the progression, with the exception of CAPRESE and CAPRI which succeed in around half of the cases (Supplementary Table 3). Edmond and Gabow display a slightly better performance than the remaining techniques, yet remarkably worse than the SCS case.

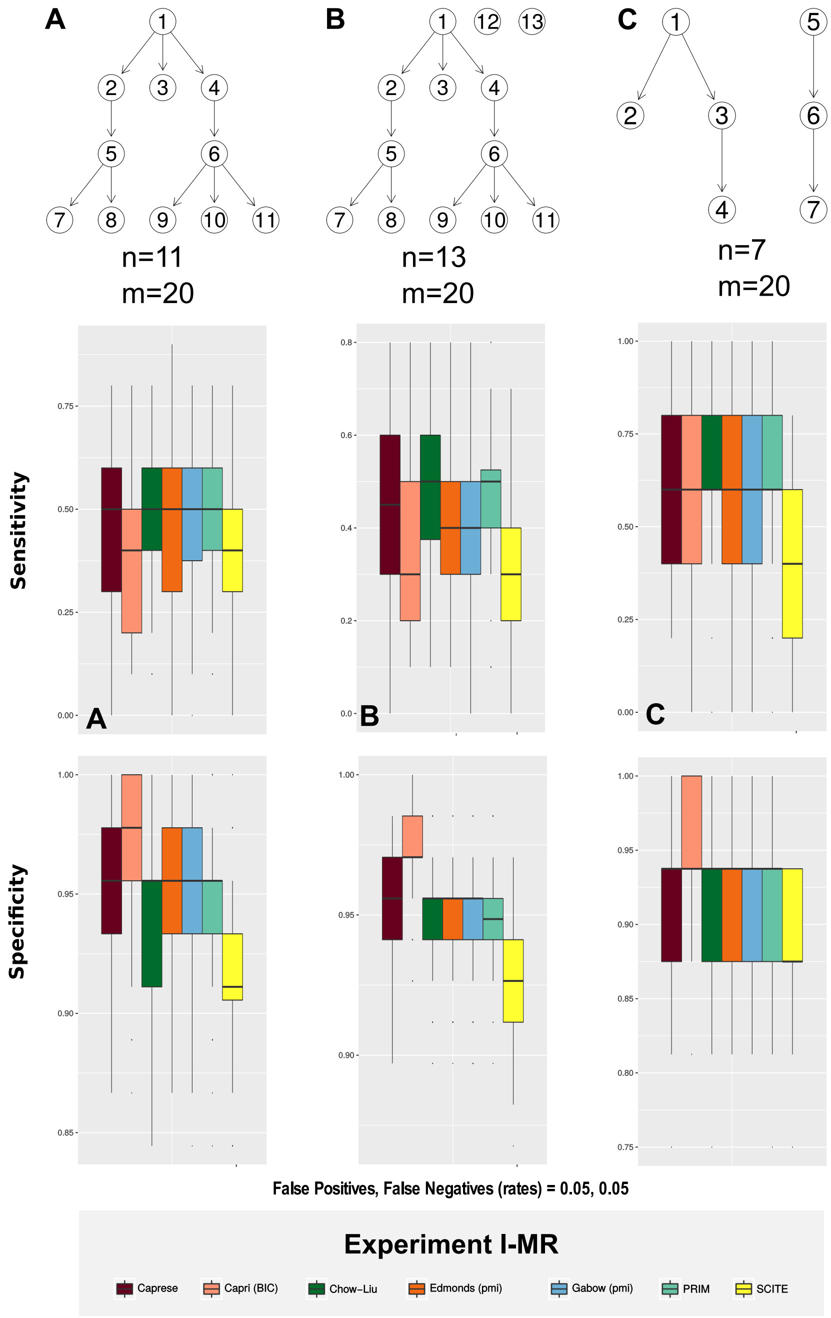

Supplementary Figure 6: Experiment I-MR (MR data). We perform the analogous of the test shown in Supplementary Figure 4, but with multi-region data. **A.** Branching evolution ($n = 11$, $m = 20$). **B.** Confounding factors. ($n = 8$, $m = 20$). **C.** Multiple independent trajectories.

| | Roots | | |
|---|---|---|---|
| | 1 | 2 | 3 |
| CAPRESE | 0 | 98 | 2 |
| CAPRI | 0 | 100 | 0 |
| CHOW-LIU | 0 | 100 | 0 |
| PRIM | 0 | 83 | 17 |
| GABOW | 0 | 100 | 0 |
| EDMONDS | 0 | 100 | 0 |
| SCITE | 0 | 100 | 0 |

Supplementary Table 2: Experiment I (multiple independent trajectories). Number of inferred models with 1, 2 or 3 distinct roots, from 100 different SCS as in Supplementary Figure 4C.

| | Roots | | | | |
|---|---|---|---|---|---|
| | 1 | 2 | 3 | 4 | 6 |
| CAPRESE | 34 | 51 | 11 | 4 | 0 |
| CAPRI | 7 | 43 | 35 | 12 | 3 |
| CHOW-LIU | 96 | 2 | 1 | 1 | 0 |
| PRIM | 96 | 2 | 1 | 1 | 0 |
| GABOW | 81 | 16 | 2 | 1 | 0 |
| EDMONDS | 82 | 15 | 2 | 1 | 0 |
| SCITE | 98 | 2 | 0 | 0 | 0 |

Supplementary Table 3: Experiment I-MR. (multiple independent trajectories). Number of inferred models with 1 to 6 roots, from 100 different multi-region datasets generated from a forest with 2 roots. Input models have $n = 7$ nodes, see Supplementary Figure 6C.

### 4 Detailed tests with single-cell data

We present results from a large-scale test with single-cell data generated (*i*) from biologically plausible phylogenetic models, and (*ii*) from a large number of randomly generated topologies. We consider three scenarios (branching evolution, confounding factors, and multiple independent trajectories).

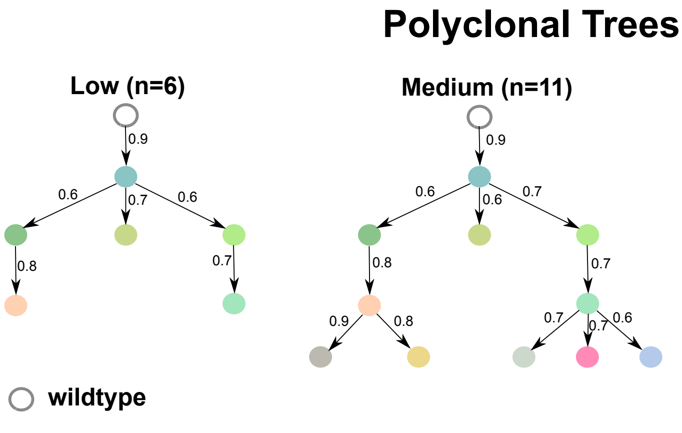

Supplementary Figure 7: Polyclonal trees used to study the performance of BitPhylogeny in [22]. These have different number $n$ of clones, which we classify as low, medium and high. We use them to generate synthetic data for some tests, e.g., in Experiment II. The wild-type is a fake node; in our models, we do not have it.

#### 4.1 Branching evolution

##### Experiment II

*We consider polyclonal tumors originating from a unique cell, in a single-cell sequencing experiment. To generate data we fix the phylogenetic trees in Supplementary Figure 7 [22]. These trees have variable number of clones ($n = 6, 11, 17$); we use them to sample different number of sequenced cells ($m = 10, 50, 100$). Besides one ideal noise-free setting, we perturb data with plausible medium and high asymmetric noise rates ($\epsilon_+/\epsilon_-$), in order to mimic characteristics errors in sampling cells and calling mutations. We compare the performance with over 4000 independent tests.*

##### Results (Supplementary Figure 8).

Reasonably, the overall performance of each algorithm is higher with lower levels of noise and larger datasets. In the ideal cases of noise-free data and 100 sampled cells, for instance, all algorithms converge to the true generative model. Noteworthy, in many realistic cases, median sensitivity and specificity measures are above 90%. The overall performance trend depends on model size ($n$), the smaller models being easier to infer as one might expect. For all algorithms, the ability to detect true relations (sensitivity) clearly drops for pathological settings (e.g., we infer 20% of the true edges for the 17-clones model, when we sequence 10 cells).

Gabow, Edmond and SCITE, display a similar superior ability to infer the true relations (i.e., high sensitivity). However, SCITE seems to overfit (i.e., with a 10% loss of specificity for the 17-clones model, when we sequence 100 cells). This is particularly evident with small datasets and models. It also persists with larger models, most likely because of the larger search space for its MCMC heuristics. CAPRI, as expected, shows very high specificity but low sensitivity, due to BIC regularization.

**Experiment III.** *We generalize Experiment II to* 100 *randomly sampled topologies with variable number of nodes* ($n = 5, 10, 20$)*. This shall avoid any bias induced by holding fixed the polyclonal topologies in Supplementary Figure 7.*

**Results (Supplementary Figure 9):** In general, the results partially reflect those of Experiment II. All the algorithms display very good performances in most settings, in many cases converging to the generative topologies (especially with small models, i.e., with $n = 5, 10$ nodes). A very similar and optimal overall performance is that obtained by Gabow, Edmond, CAPRESE and SCITE, with minor differences in the different parameter settings, yet with an evident tendency of SCITE in inferring denser models with more false positives (i.e., highlighting a lower specificity). Also in this case CAPRI show a very good specificity because of the regularization, but fails in capturing many true positives.

**Experiment IV.** *In order to assess the robustness of the inference with respect to different rates of false positives and false negatives rates provided as input to the algorithms, we investigated the variation of the performance of two selected algorithms, namely Gabow and SCITE, on a dataset generated from the* Medium *phylogenetic tree in Supplementary Figure 7, with* $n = 11$ *nodes and* $m = 75$ *samples,* $\epsilon_+ = 5 \times 10{-}3$ *and* $\epsilon_- = 5 \times 10{-}2$*, for the* 25 *possible combinations of input* $\epsilon_+$ *and* $\epsilon_-$ *in the following ranges:* $\epsilon_+ = (3, 4, 5, 6, 7) \times 10^{-3}$ *and* $\epsilon_- = (3, 4, 5, 6, 7) \times 10^{-2}$.

**Results (Supplementary Table 4 and Supplementary Table 5):** By looking at the performance with respect to the different combinations of $\epsilon_+$ and $\epsilon_-$ provided as input to the algorithms, we unexpectedly do not observe noteworthy variations, for both the algorithms. This result indicates that, if the value of noise provided as input to the algorithms is close to the real value (i.e., within a reasonable range), the inference accuracy is not remarkably perturbed. As a consequence, one might question about the usefulness of the usually computationally expensive techniques used for the inference of noise models, as done, for instance, by SCITE. We leave some further comments on this topic to the main text.

#### 4.2 Confounding Factors

Here true variables are mixed to random 0/1 variables, totally unrelated to the progression. This could be a simple model of uncertainty in the calling, where we over-call variants that are not true related to the progression at a certain error rate.

**Experiment V.** *We use data from Experiment II; to each dataset we add random binary columns. A column is a repeated sampling of a* biased coin*, with bias uniformly sampled among the marginals of all events.* $n \times 10\%$ *random columns are inserted per dataset, where* $n$ *is the true model size.*

**Results (Supplementary Figure 10):** Surprisingly, the results of this experiment reflects those of Experiment II, with minor differences in the performance of the various algorithms in the different settings. The overall performance is good, at least with sufficiently large datasets and sufficiently small levels of noise. Also in this case, SCITE tends to overfit, especially with larger models and small datasets.

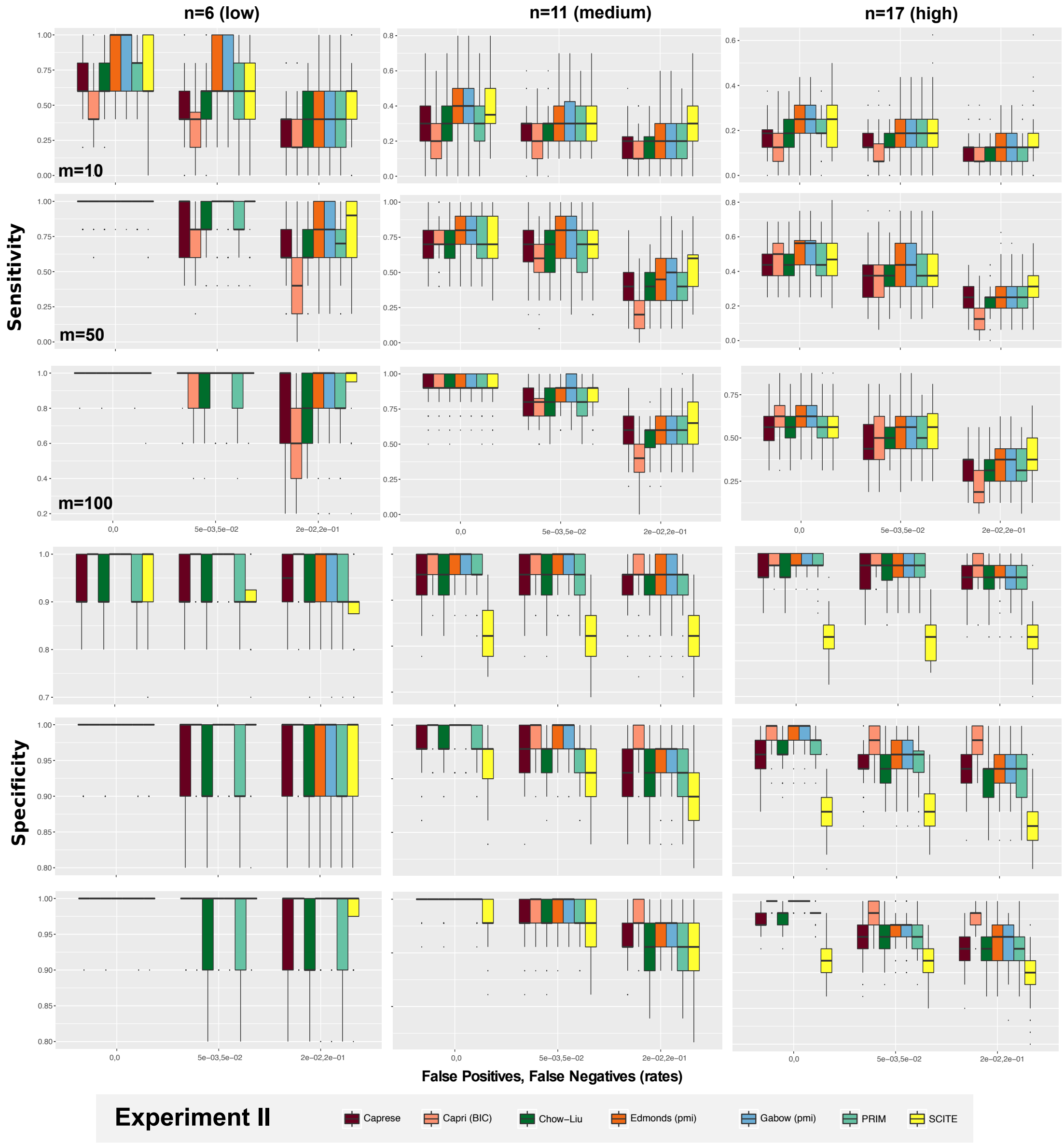

Supplementary Figure 8: Experiment II, (branching evolution, fixed topologies, SCS data). All the algorithms are tested on datasets generated from the phylogenetic trees shown in Supplementary Figure 7. Three incremental level of unbalanced noise in the data are assumed: $\epsilon_+ = \epsilon_- = 0$ (noise-free), $\epsilon_+ = 5 \cdot 10^{-3}$, $\epsilon_- = 5 \cdot 10^{-2}$ and $\epsilon_+ = 2 \cdot 10^{-2}$, $\epsilon_- = 2 \cdot 10^{-1}$. We test distinct sample set sizes ($n = 10, 50, 100$) and 100 distinct datasets for each case, reporting the distributions of sensitivity and specificity.

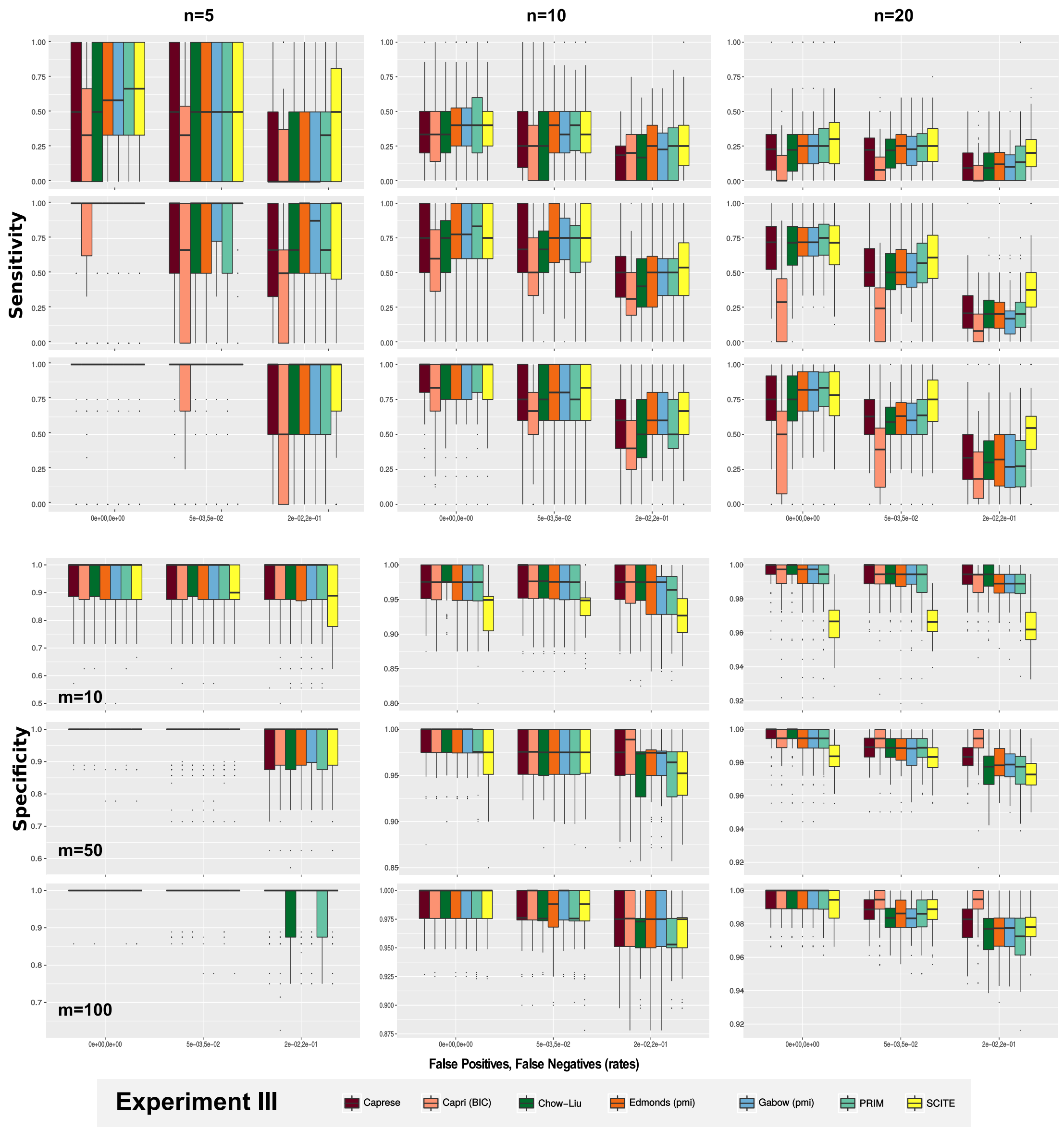

Supplementary Figure 9: Experiment III. (branching evolution, random topologies, SCS data). The algorithms are tested on SCS data generated from 100 random tree topologies, with $n = 5, 10, 20$ clones. As in Experiment II, three levels of noise ($\epsilon_+$, $\epsilon_-$) and three sample sizes ($m$) are tested.

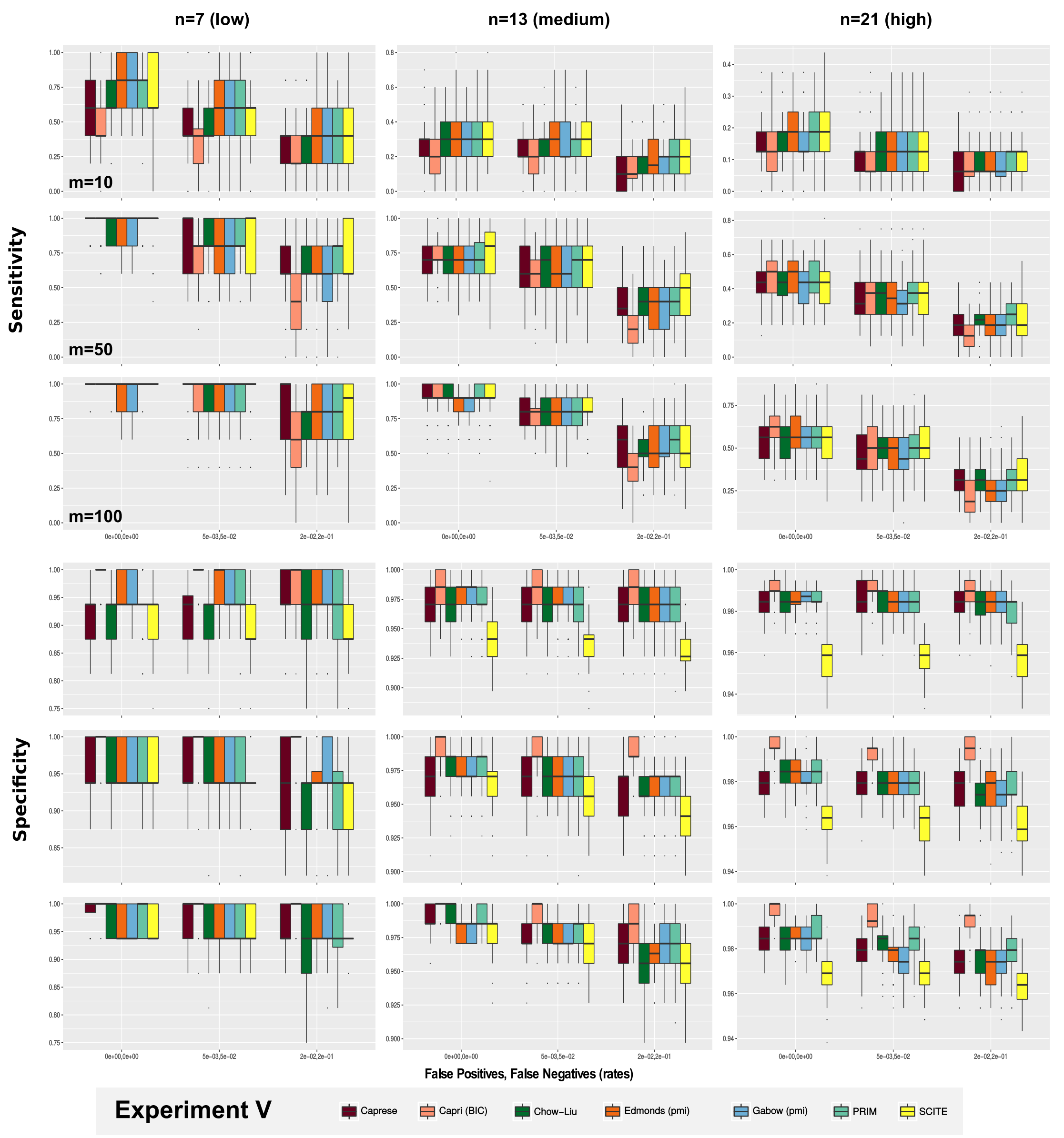

Supplementary Figure 10: Experiment V. (confounding factors, fixed topologies, SCS data). We add $n \times 10\%$ random (0/1) columns to the SCS data generated from the trees in Supplementary Figure 7. We test three levels of noise ($\epsilon_+ = \epsilon_- = 0$, $\epsilon_+ = 5 \cdot 10^{-3}$, $\epsilon_- = 5 \cdot 10^{-2}$, $\epsilon_+ = 2 \cdot 10^{-2}$, $\epsilon_- = 2 \cdot 10^{-1}$), and three sample sizes ($m = 10, 50, 100$).

#### 4.3 Multiple Independent Trajectories

In this case the signal that we detect in the data is composed from different true signals, one per population of cells. So, we need to infer a forest with a number of trees equal to the number of different progressions, which it seems reasonable to assume to be low, e.g., below 5.

**Experiment VI.** *We extend the sampling strategy in Experiment III to account for forests with fixed total number of nodes, i.e.,* $n = 20$*. We perform the same procedure of that experiment.*

**Results (Supplementary Figure 11):** All the algorithms display a very low sensitivity with small datasets (with 20% median value with $m = 10$ samples), remarkably increasing the performance with larger datasets (median values around 75% with $m = 100$ samples in the noise-free case). Gabow, Edmond and CAPRESE show a good tradeoff between sensitivity and specificity, displaying a good and similar performance, whereas SCITE confirms the tendency to overfit for small datasets, yet being the most robust algorithm against noise in the data.

#### 4.4 Inference with missing data

In addition to the false positives/negatives introduced in the data via allelic dropouts and false alleles, unobserved or missing data points represent another major problem when dealing with single-cell sequencing data. In the early works, around 60% of the data were missing due to the low quality of the sequencing technique. Even though the technology has remarkably improved during the last few years, leading to more reliable and usable data, we here investigate the influence of missing data on the inference, with respect to the considered algorithms. In particular, we performed simulated experiments with a specific generative topology and with different amounts of missing data, ranging from 10% to 40%, analyzing the variation in the inference accuracy.

**Experiment VII** *In order to evaluate the impact of missing data on the inference accuracy, we chose* 20 *benchmark single-cell datasets generated from the* Medium *phylogenetic tree in Supplementary Figure 7, with* $n = 11$ *nodes and* $m = 75$ *samples.* 10 *of these datasets were generated with* $\epsilon_+ = \epsilon_- = 0$*, and the remaining* 10 *datasets with (ii)* $\epsilon_+ = 0.005$*,* $\epsilon_- = 0.05$*. For each of the* 20 *datasets we generated* 5 *further datasets, with the following ratio of randomly included missing entries:* $r = (0, 0.1, 0.2, 0.3, 0.4)$*, for a total of* 100 *distinct datasets. As SCITE naturally deals with datasets with missing data, we performed the inference with no further parameters. Instead, in order to perform the reconstruction with the remaining algorithms, we followed this procedure. For each one of the* 80 *datasets with missing data (we did not consider the case with* $r = 100$*), we filled the missing entries via a classical Expectation Maximization (EM) algorithm, and we repeated this step to create* 100 *complete datasets (for each incomplete datasets). We then performed the inference with all the algorithms on all the* 100 *datasets in each case, selecting the model with the best likelihood score, which was then used in the performance assessment.*

**Results (Supplementary Figure 12):** As one can see in Supplementary Figure 12, the performance of all the algorithms is profoundly affected by the presence of missing data, both in the noisy and in the noise-free cases. SCITE displays an overall more robust sensitivity than the other techniques, yet in spite of a worse specificity, which would point at a tendency toward overfitting also in this

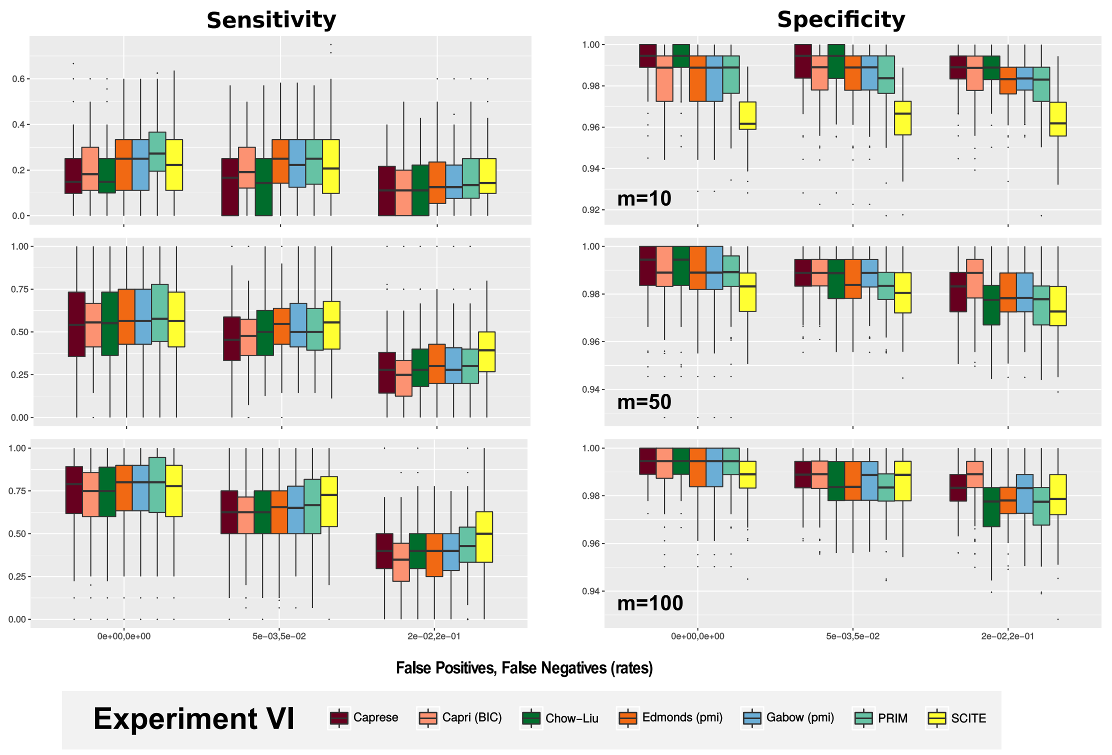

Supplementary Figure 11: Experiment VI. (multiple independent trajectories, SCS data). Single-cell datasets are generated from a large number (i.e., 100 for each case) of random forest topologies, with $n = 20$ distinct clones. Three levels of noise ($\epsilon_+ = \epsilon_- = 0$, $\epsilon_+ = 5 \cdot 10^{-3}$, $\epsilon_- = 5 \cdot 10^{-2}$, $\epsilon_+ = 2 \cdot 10^{-2}$, $\epsilon_- = 2 \cdot 10^{-1}$), and three sample sizes ($m = 10, 50, 100$) are tested.

scenario. As expected, the performance of all the techniques is significantly better in the noise-free case and, in general, is maintained at acceptable levels up to values of missing data around 20%/30% according to the cases.

### 5 Detailed tests with multi-region bulk-sequencing data

As we did for SCS data, we here present the results of detailed comparative tests of with multi-region sequencing data. The analyses are organized as for SCS data.

#### 5.1 Branching evolution

**Experiment II-MR** *Here we reproduce Experiment II in the case of multi-region data, and sample the polyclonal topologies shown in Supplementary Figure 7, with symmetric noise rates ($\epsilon_- = \epsilon_+$). We compare the boxplot performance from 100 tests.*

**Results (Supplementary Figure 13):** The accuracy of most algorithms is good in all the scenarios. They all reach high values of specificity, whereas satisfactory values of sensitivity are observed only with combination of sufficiently large datasets and sufficiently low noise. As expected, the overall performance worsens with larger and more complex generative models.

In this case Gabow and Edmond display the best efficiency in retrieving both the true positives and negatives, Edmond being slightly better in a certain number of parameter settings. Conversely, SCITE shows the worst performance, especially with small datasets and low levels of noise, yet proving a certain robustness to the increase in the noise level. CAPRI displays very good values of specificity even with these data type, yet most likely due to its regularization.

**Experiment III-MR** *This experiment reproduces Experiment III with multi-region data, hence sampling the datasets from 100 randomly generated topologies with variable number of nodes.*

**Results (Supplementary Figure 14):** The results of this experiment resemble Experiment III: overall good performance is observed, yet with low sensitivity with small datasets and noisy data. Gabow and Edmond consistently display optimal and very similar trends, with Edmond showing a better performance in a sightly larger number of settings. SCITE confirms to be less accurate with multi-region data, especially with small datasets, even when the level of noise is low. We remark that we did not include an experiment analogous to Experiment IV because with symmetrical noise rates we do not expect significant differences in accuracy.

#### 5.2 Confounding factors

**Experiment V-MR** *We reproduce Experiment V with multi-region data augmented with $n \times 10\%$ random columns. We use the three topologies shown in Supplementary Figure 7.*

**Results (Supplementary Figure 15):** The results are in accordance with those of the analogous SCS experiment, with an expected overall decrease of accuracy due to the introduction of spuriously correlated events. Edmond is the most accurate algorithm, slightly improving over Gabow; SCITE seems less efficient in retrieving both the true and the false relations, especially with small datasets and/ or low noise levels.

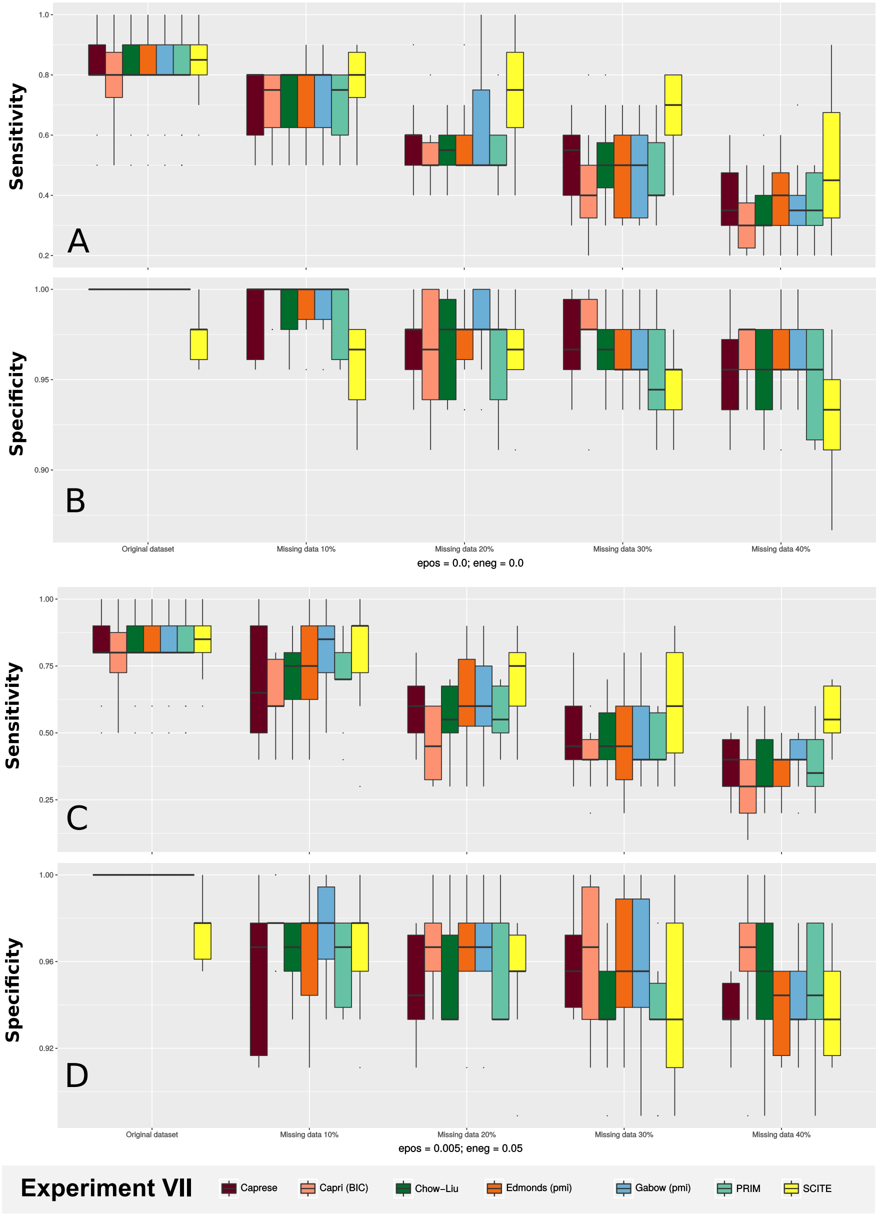

Supplementary Figure 12: Experiment VII. (missing data, SCS data). Sensitivity and specificity for with different proportions of missing entries, i.e., $r = (0, 0.1, 0.2, 0.3, 0.4)$, and different levels of noise: $(i)$ $\epsilon_+ = \epsilon_- = 0$ and $(ii)$ $\epsilon_+ = 0.005$, $\epsilon_- = 0.05$. The original dataset is generated from the medium tree in Supplementary Figure 7, with $n = 11$ nodes and $m = 75$ samples.

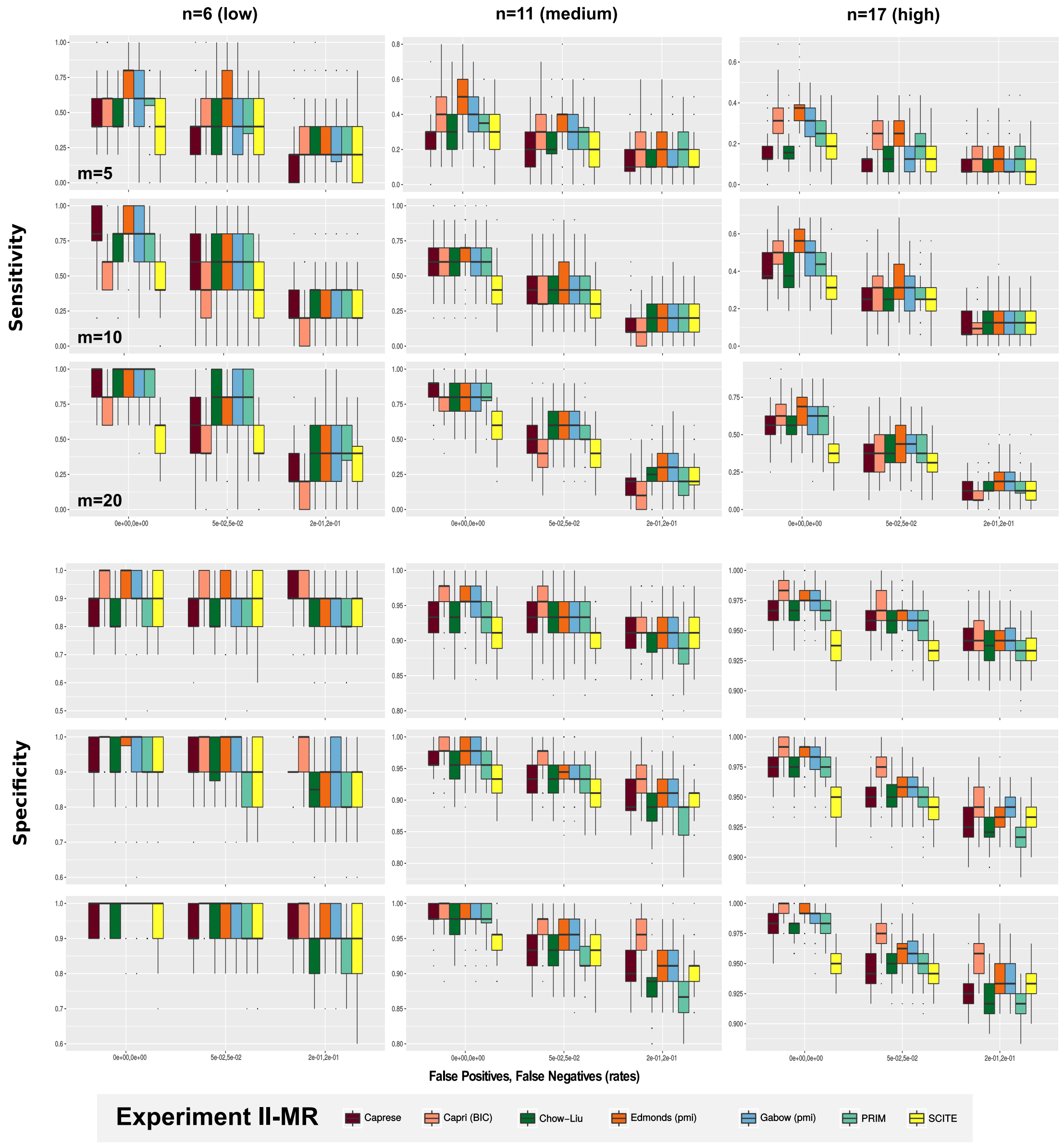

Supplementary Figure 13: Experiment II-MR. (branching evolution, fixed topologies, MR data). All algorithms are tested on datasets generated from the phylogenetic trees shown in Supplementary Figure 7, where we simulate multi-region data. Three levels of noise ($\epsilon_+ = \epsilon_- = 0.0, 0.05, 0.2$), and three sample sizes ($m = 5, 10, 20$) are tested.

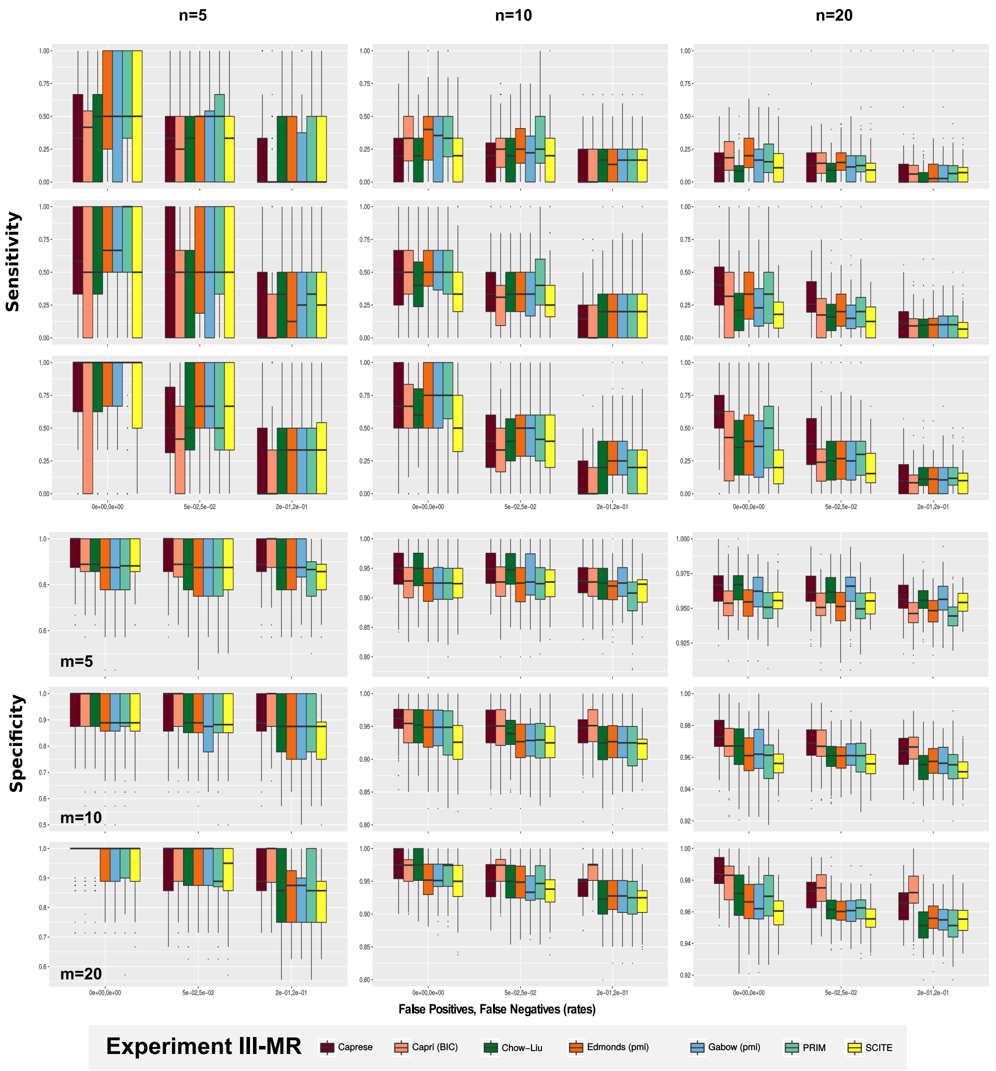

Supplementary Figure 14: Experiment III-MR (branching evolution, random topologies, MR data). The algorithms are tested on multi-region datasets generated from a number of random tree topologies, with $n = 5, 10, 20$ clones (100 distinct topologies for each case). Three levels of noise ($\epsilon_+ = \epsilon_- = 0.0, 0.05, 0.2$), and three sample sizes ($m = 5, 10, 20$) are tested.

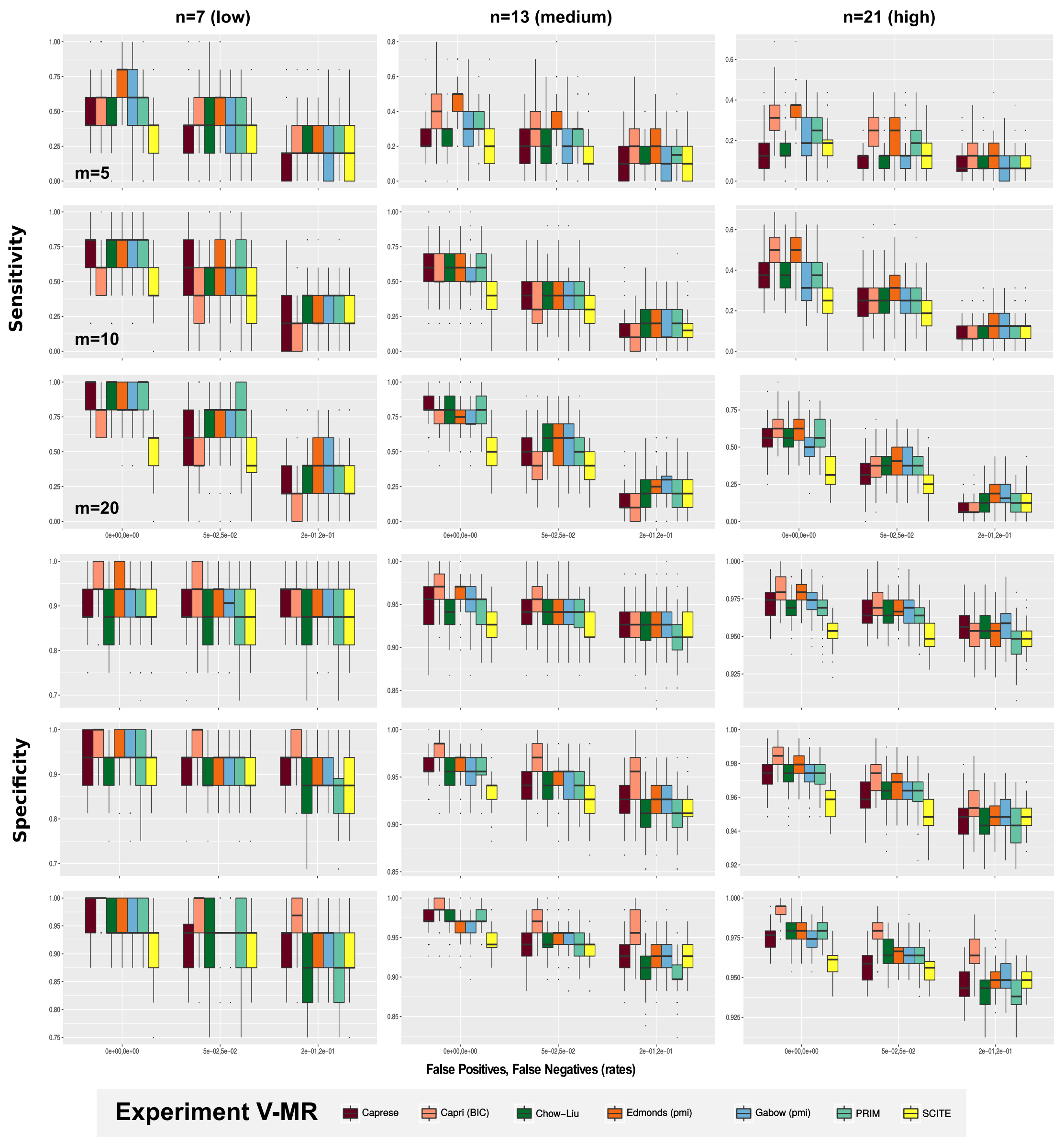

Supplementary Figure 15: Experiment V-MR. (confounding factors, fixed topologies, MR data). $n \times 10\%$ random (0/1) columns are added to the multi-region datasets generated from the trees shown in Supplementary Figure 7. Three levels of noise and sample size are tested.

#### 5.3 Multiple Independent Trajectories

**Experiment VI-MR** *We reproduce Experiment VI with multi-region data, and with datasets sampled from random forests (see Supplementary Figure 3).*

**Results (Supplementary Figure 16):** Gabow, Edmond and CAPRESE appear to be the most accurate algorithms in this scenario. The former achieves the best sensitivity and specificity in most settings. Prim is very efficient in retrieving the true relations in many cases, especially with low levels of noise, yet presenting a certain tendency toward overfitting. SCITE seems less accurate in most settings.

### 6 Parameters settings, computation time and scalability

#### 6.1 Parameter Settings

$n$ : **number of nodes** (i.e, genomic alterations/ clones).
We use

$$n = 5, 10, 15, 20$$

when we sample random models. In random tests, 100 trees are generated for each configuration of $n$ and $m$ (see below), and one dataset per tree is sampled. In some tests, we fix $n$ to $6, 11$ and $17$ (Supplementary Figure 7); we specify in the experiment description if that is the case.

$m$ : **number of samples** (i.e., cells, or regions sequenced).
When we perform a single-cell sampling, we scan the values

$$m = 10, 25, 50, 75, 100\,.$$

When we perform a multi-region sampling, we scan values in line with a reasonable number of biopsies that could be extracted from a solid tumor

$$m = 5, 7, 10, 20, 50\,.$$

$c$ : **number of signals from single-clones in a multi-region experiment**.
When we perform a multi-region sampling, we set the values

$$c = 3, 5, 8\,.$$

for the fixed topologies in Supplementary Figure 7, and

$$c = \frac{n}{2}$$

otherwise.

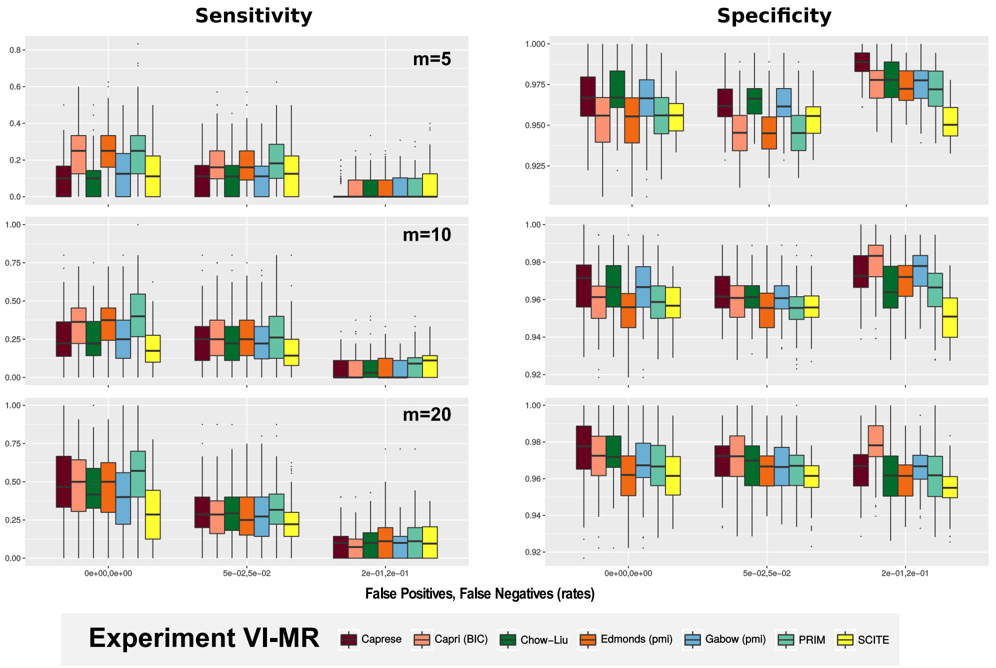

Supplementary Figure 16: Experiment VI-MR. (multiple independent trajectories, MR data). Multi-region datasets are generated from a number (i.e., 100 for each case) of random forest topologies, with $n = 20$ distinct clones. Tthree levels of noise ($\epsilon_+ = \epsilon_- = 0, 0.05, 0.2$) and three sample sizes ($m = 5, 10, 20$) are tested, recording sensitivity and specificity.

$\epsilon_+/\epsilon_-$ : **rates of FPs/FNs in data**, for the observed genotypes.
In single-cell sequencing we assume these to be

$$(\epsilon_+, \epsilon_-) = (0, 5, 10, 15, 20, 25, 30, 35) \times (10^{-3}, 10^{-2}) .$$

This corresponds to pairs of equal value that differ for an order of magnitude, e.g., ($\epsilon_+ = 0.015, \epsilon_- = 0.15$), consistently with the observation that false negative rates in single-cell data are much higher than false positives ones. Such values correspond to an overall error rate that is $2 \times \epsilon_+$ and $2 \times \epsilon_-$. For multi-region sequencing, these errors are symmetric

$$(\epsilon_+, \epsilon_-) = (0, 5, 10, 15, 20, 25, 30, 35) \times (10^{-2}, 10^{-2}) .$$

*topology* : **model structure** used according to the working scenarios (Supplementary Figure 3). This is either a tree, a forest or a DAG.

*probabilities* : **conditional probability tables** (CPTs), and **marginals** of a model.
When these are sampled at random we impose the constraint that for any pair of variables $X$ and $Y$ it holds

$$p(X \mid Y) \in [0.6, 0.9],$$

and for any marginal $p(X) > 0.001$. These values seem reasonable to avoid the introduction of biases in the sampling process. In some cases we assigned fixed values to the CPTs, which we report in the corresponding figures (e.g., in Supplementary Figure 7).

$p_\star = 0.05$ : **$\alpha$-level of the Mann-Whitney test** (p-value).

#### 6.2 Computation time and scalability

To assess and compare the computation time of the distinct techniques we used the Medium phylogenetic tree in Supplementary Figure 7 as generative topology, with $n = 11$ nodes, $m = 75$ samples, $\epsilon_+ = 0.005$, $\epsilon_- = 0.05$, and we repeated the inference for 100 distinct experiments, on a single core of a Lenovo Thinkpad t430s with an Intel i7 3520M 4-core 2.90GHz and 16Gb Ram.

In Supplementary Table 6 we see that CAPRESE is the fastest algorithm, because it does not bootstrap the data. It is followed by Prim, Gabow, Edmond, CAPRI and Chow-Liu, with almost identical running time (7× slower than CAPRESE). SCITE is remarkably slower (i.e., 25× slower than CAPRESE and more than 3× slower than the group of PRIM), whereas OncoNEM has the worst performance (i.e., 300× slower than CAPRESE, around 40× slower than CAPRI, and 12× slower than SCITE). For these reasons, we could not include also OncoNEM in more extensive experiments.

In order to assess TRaIT's scalability with increasingly larger single-cell datasets, we generated 100 random branching evolution topologies (as in Experiment III and Supplementary Figure 9), with $n = 20$ nodes, $\epsilon_+ = 0.005$, $\epsilon_- = 0.05$ and different values of sample size: $m \in (100, 500, 1000, 5000, 10000, 15000, 20000)$. We then timed both EDMONDS and CHOW-LIU, and evaluated performance. In Supplementary Table 7 we show median values and standard deviation for each distinct experimental settings. Simulations were performed on a Quad Core pc with Intel i7 - 8 thread 3.5 GHz and 16Gb Ram. As one can can see from the table, CHOW-LIU displays better median sensitivity with smaller sample size than EDMONDS, which in turn shows better median specificity. Both the algorithms display an approximately linear increase of the computational time with respect to the number of samples, and even for extremely large datasets (i.e., 20000 single cells) the median execution time is around 22 seconds per experiment.

### 7 Case studies

In addition to the Main Text, we show in Supplementary Figures 17 and 18 the inference from the triple-negative breast cancer SCS data, and in Supplementary Figure 20 the same for the colorectal cancer data. In Supplementary Figure 21 we show the fit with SCITE .

### 8 Noise model

Derivation of the noise model for both marginal and joint probabilities.

#### Marginal Probabilities

Let us call $\epsilon_+$ the probability of observing 1 when we had 0 (false positive) and $\epsilon_-$ the other way around (false negative). We remark that we assume these probability to be strictly in $[0, 0.5)$ with value 0.5 representing totally random entries. Then, for any event $\mathbf{x}_i$, we can write the expectation of the probability of observing it (here with the notation $n_i$), given its theoretical probability $p(\mathbf{x}_i)$ as follow.

$$n_i = p(\mathbf{x}_i) \cdot [1 - \epsilon_-] + [1 - p(\mathbf{x}_i)] \cdot \epsilon_+ \,,$$

and with some rearrangements,

$$n_i = p(\mathbf{x}_i) - p(\mathbf{x}_i) \cdot \epsilon_- + \epsilon_+ - p(\mathbf{x}_i) \cdot \epsilon_+ \,,$$
$$n_i = p(\mathbf{x}_i) \cdot [1 - \epsilon_+ - \epsilon_-] + \epsilon_+ \,,$$

from which, with $(\epsilon_+, \epsilon_-) \in [0, 0.5)$,

$$p(\mathbf{x}_i) = \frac{n_i - \epsilon_+}{1 - \epsilon_+ - \epsilon_-} \,.$$

#### Joint probabilities

Let us now consider the theoretical joint probability $p(\mathbf{x}_{i,j})$ (in what follow also called $p(\mathbf{x}_i, \mathbf{x}_j)$ to make the elements of the probability explicit) of any two pair of events and the respective observed marginal and joint probabilitis $n_i$, $n_j$ and $n_{i,j}$. Then, the expectation of $n_{i,j}$ can be written as follow.

$$\begin{aligned} n_{i,j} \quad = \quad & p(\mathbf{x}_i, \mathbf{x}_j) \cdot [1 - \epsilon_-] \cdot [1 - \epsilon_-] \\ & + \; p(\mathbf{x}_i, \overline{\mathbf{x}_j}) \cdot [1 - \epsilon_-] \cdot \epsilon_+ \\ & + \; p(\overline{\mathbf{x}_i}, \mathbf{x}_j) \cdot \epsilon_+ \cdot [1 - \epsilon_-] \\ & + \; p(\overline{\mathbf{x}_i}, \overline{\mathbf{x}_j}) \cdot \epsilon_+ \cdot \epsilon_+ \,, \end{aligned}$$

and being,

$$\begin{aligned} p(\mathbf{x}_i, \overline{\mathbf{x}_j}) &= p(\mathbf{x}_i) - p(\mathbf{x}_i, \mathbf{x}_j) \,, \\ p(\overline{\mathbf{x}_i}, \mathbf{x}_j) &= p(\mathbf{x}_j) - p(\mathbf{x}_i, \mathbf{x}_j) \,, \\ p(\overline{\mathbf{x}_i}, \overline{\mathbf{x}_j}) &= 1 - p(\mathbf{x}_i) - p(\mathbf{x}_j) + p(\mathbf{x}_i, \mathbf{x}_j) \,, \end{aligned}$$

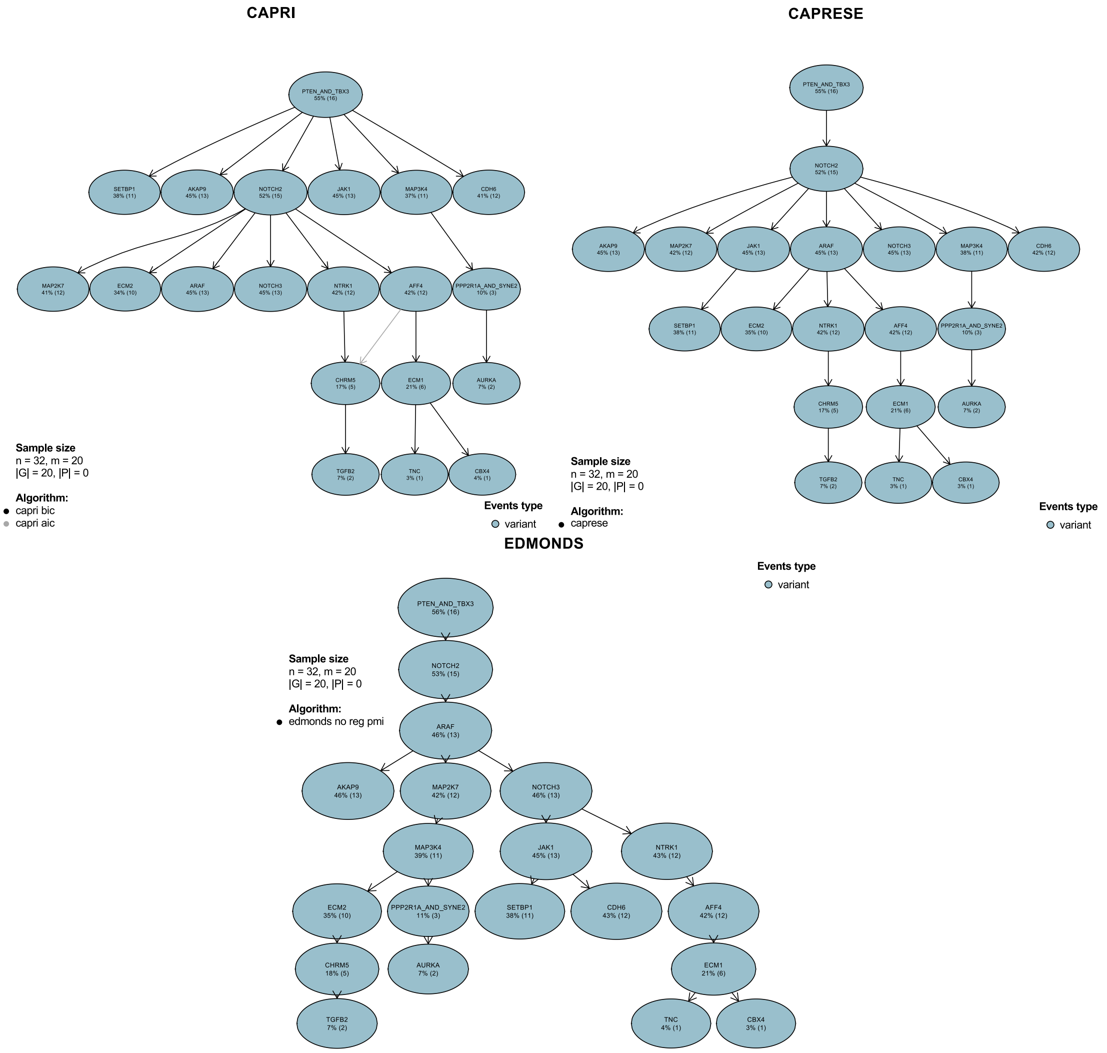

Supplementary Figure 17: Analysis of the data from [23] (single-nucleus exome sequencing of 16 tumor cells from a triple-negative breast cancer) with CAPRI, CAPRESE and Edmond.

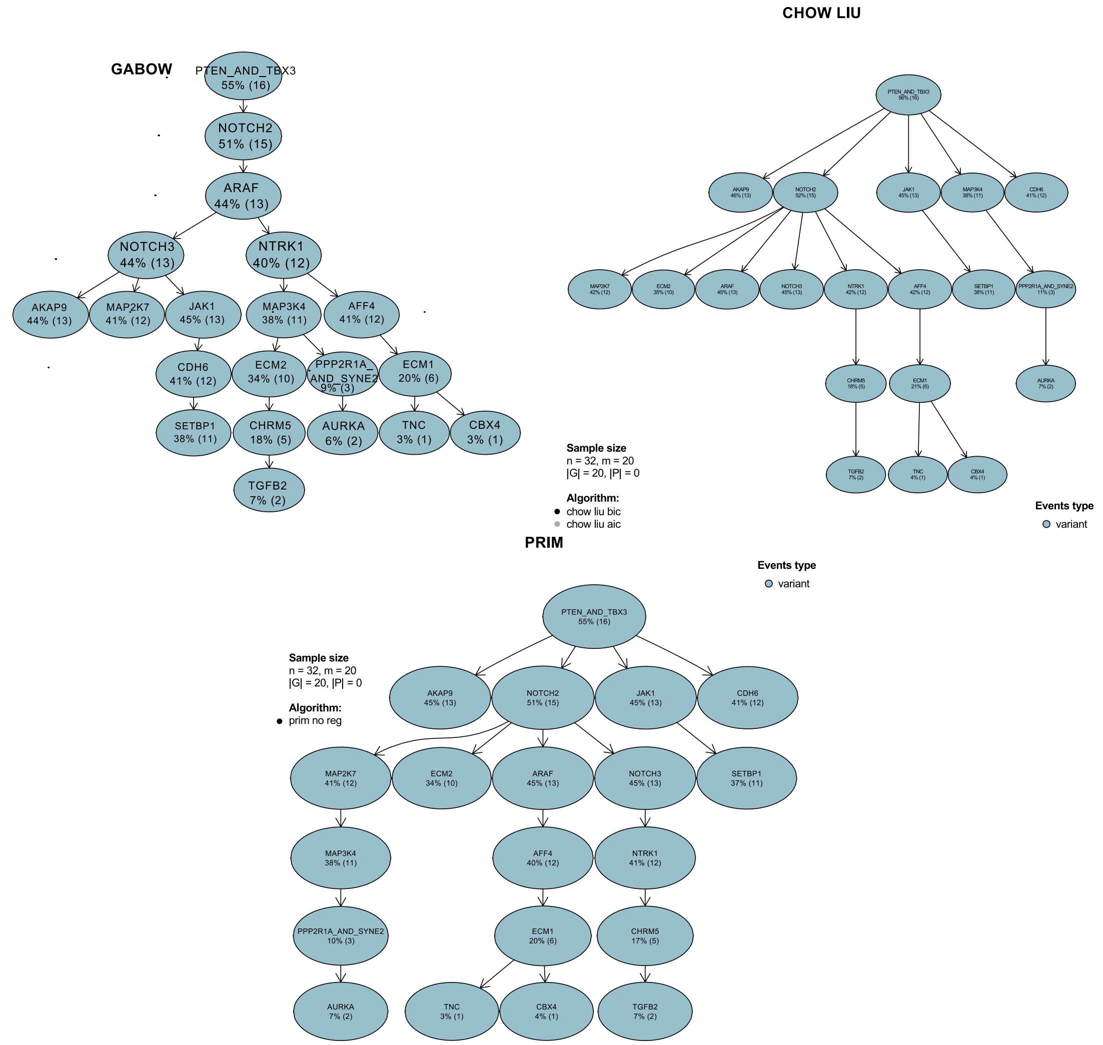

Supplementary Figure 18: Analysis of the data from [23] (single-nucleus exome sequencing of 16 tumor cells from a triple-negative breast cancer) with Gabow, Chow-Liu and PRIM.

with some rearrangements,

$$\begin{aligned}
n_{i,j} &= p(\mathbf{x}_i, \mathbf{x}_j) \cdot (1 + \epsilon_+^2 + \epsilon_-^2 + 2 \cdot \epsilon_+ \cdot \epsilon_- - 2 \cdot \epsilon_+ - 2 \cdot \epsilon_-) \\
&\quad + [p(\mathbf{x}_i) + p(\mathbf{x}_j)] \cdot \epsilon_+ \cdot [1 - \epsilon_+ - \epsilon_-] + \epsilon_+^2 \,, \\
n_{i,j} &= p(\mathbf{x}_i, \mathbf{x}_j) \cdot (1 - \epsilon_+ - \epsilon_-)^2 + [p(\mathbf{x}_i) + p(\mathbf{x}_j)] \cdot \epsilon_+ \cdot [1 - \epsilon_+ - \epsilon_-] + \epsilon_+^2 \,, \\
n_{i,j} &= p(\mathbf{x}_i, \mathbf{x}_j) \cdot (1 - \epsilon_+ - \epsilon_-)^2 + \frac{n_i + n_j - 2 \cdot \epsilon_+}{1 - \epsilon_+ - \epsilon_-} \cdot \epsilon_+ \cdot [1 - \epsilon_+ - \epsilon_-] + \epsilon_+^2 \,, \\
n_{i,j} &= p(\mathbf{x}_i, \mathbf{x}_j) \cdot (1 - \epsilon_+ - \epsilon_-)^2 + (n_i + n_j) \cdot \epsilon_+ - 2 \cdot \epsilon_+^2 + \epsilon_+^2 \,,
\end{aligned}$$

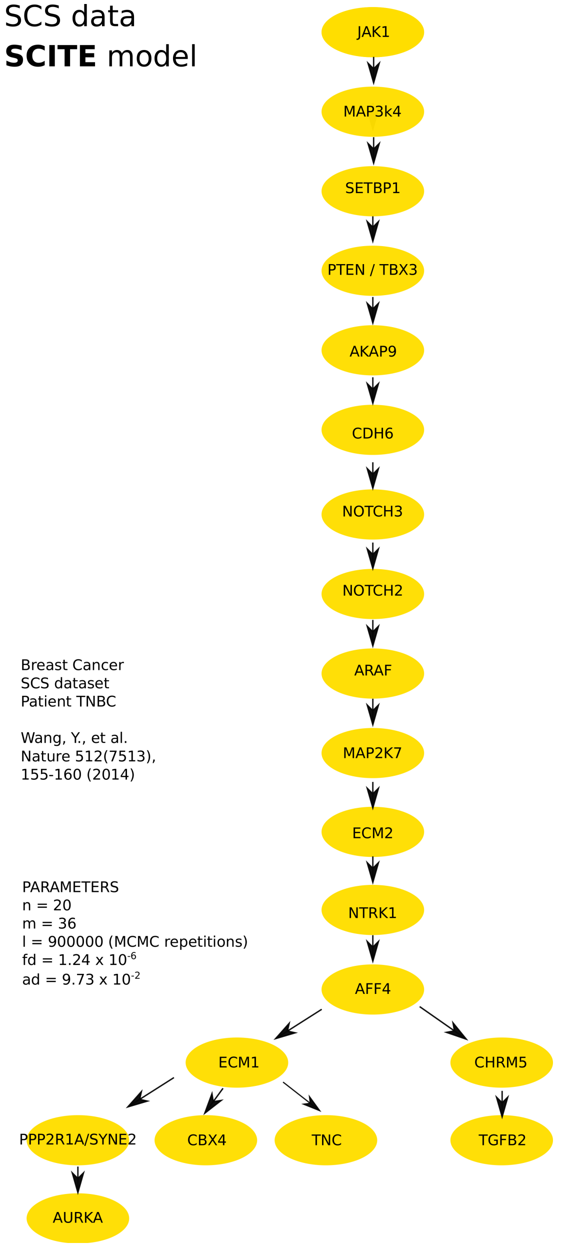

Supplementary Figure 19: Analysis of the data from [23] (single-nucleus exome sequencing of 16 tumor cells from a triple-negative breast cancer) with SCITE. False discovery rate of $1.26 \times 10^{-6}$ and allelic dropout of $9.73 \times 10^{-2}$ are provided as input parameters. Posterior estimates are computed with 900.000 Montecarlo steps, and 10000 equivalent-scoring trees returned. Here we show one of the top-scoring.

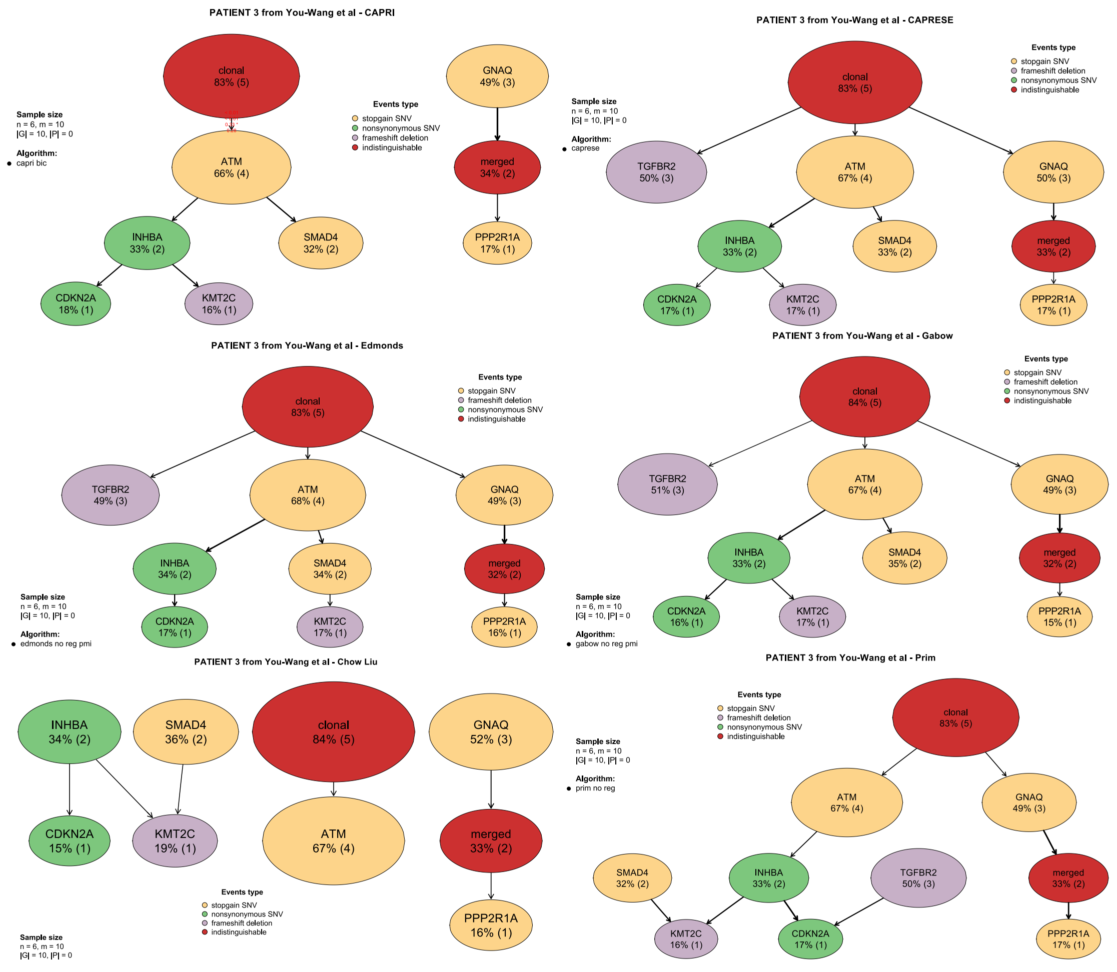

Supplementary Figure 20: Analysis by TRaIT's algorithm of the colorectal cancer MR data from [24].

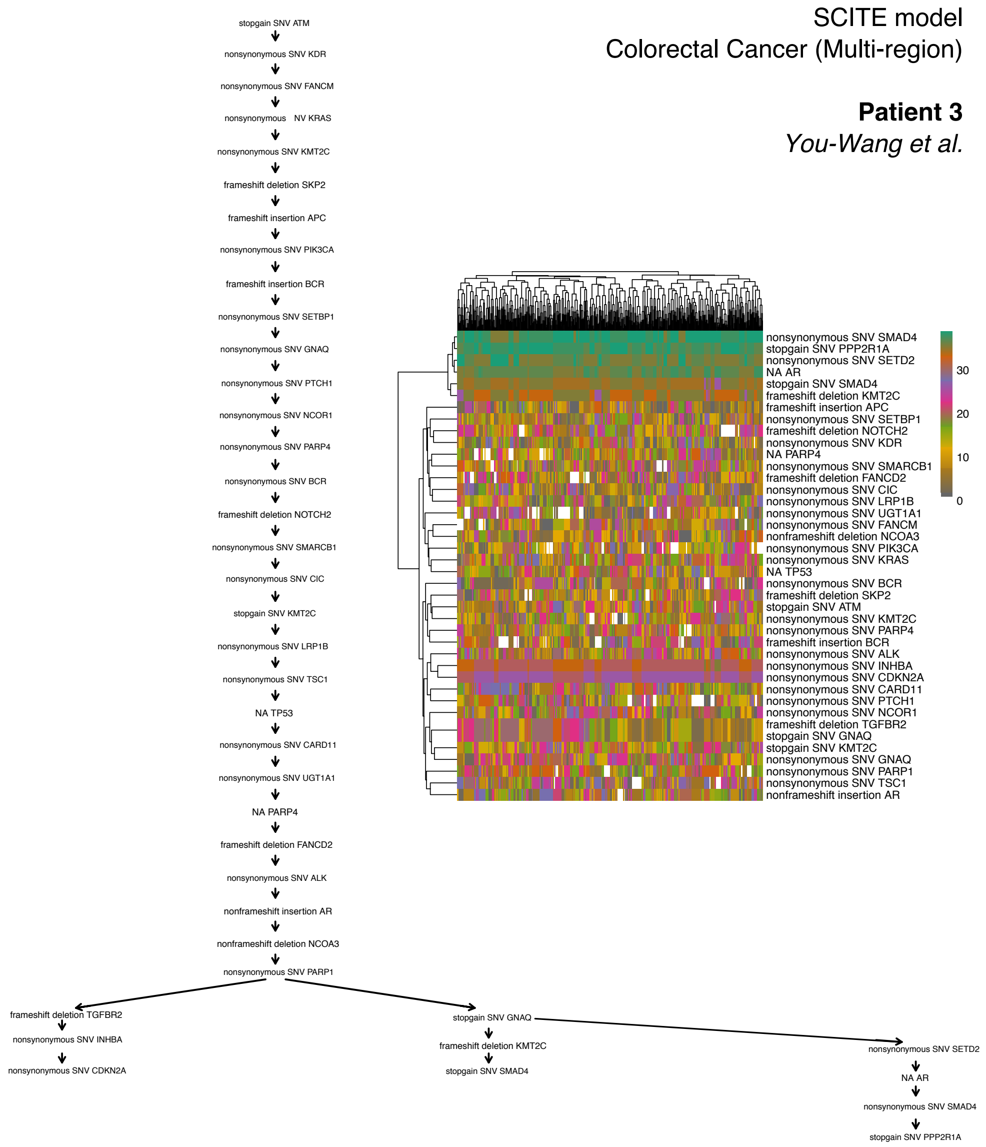

Supplementary Figure 21: Analysis by SCITE of the colorectal cancer MR data from [24]. The events occurring in the same group of samples are indistinguishable in our framework. SCITE retrieves from them a long linear chain of consecutive events, whose ordering is not unique. The posterior is in fact multi-modal, as shown via an heatmap of top-scoring models. There each column is a model, each row an event, an teh parents are coloured. Roots are white. The presence of non-uniform colours shows the posterior uncertainty for to the linear chain of events that cannot be distinguished.

A – Original input matrix

| | Mutation of X | Focal deletion of X, or deletion of a region including X | Mutation of Y | Mutation of Z |
|---|---|---|---|---|
| Sample 1 | 1 | 0 | 1 | 0 |
| Sample 2 | 1 | 0 | 0 | 0 |
| Sample 3 | 1 | 0 | 1 | 1 |
| Sample 4 | 0 | 1 | 0 | 0 |
| Sample 5 | 0 | 1 | 1 | 0 |
| Sample 6 | 0 | 1 | 1 | 0 |

B – Reshaped matrix

| | Aggregate event: mutation of X [OR] deletion of X | Mutation of Y | Mutation of Z |
|---|---|---|---|
| Sample 1 | 1 | 1 | 0 |
| Sample 2 | 1 | 0 | 0 |
| Sample 3 | 1 | 1 | 1 |
| Sample 4 | 1 | 0 | 0 |
| Sample 5 | 1 | 1 | 0 |
| Sample 6 | 1 | 1 | 0 |

C – Model

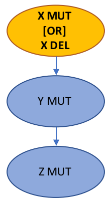

Supplementary Figure 22: Back mutation example. **A.** The input matrix includes 6 single cells and 4 genomic events: mutation of gene X, deletion of a region including X, and mutations of Y and Z. In this scenario cells 4 - 6 are evolutionary subsequent cells 1–3, and suffer a back mutation on X. **B.** The input matrix is reshaped by merging the mutation and deletion of X, creating a new event. **C.** The output model will include such aggregate event as root, prior to mutations of Y and Z, hence preserving the overall temporal coherence.

from which, with $(\epsilon_+, \epsilon_-) \in [0, 0.5)$,

$$p(\mathbf{x}_i, \mathbf{x}_j) = \frac{n_{i,j} - \epsilon_+ \cdot (n_i + n_j - \epsilon_+)}{(1 - \epsilon_+ - \epsilon_-)^2}\,.$$

### 9 Modeling back mutations

Our framework does not explicitly account for *back mutations*. These confound the inference when a previously acquired genomic alteration is lost during the evolutionary history of a tumor, due, e.g., to loss of heterozygosity or general chromosomal deletions. In phylogenetic jargon this is a violation of the no-back mutation assumption, and cannot be handled in general cases.

However, when both SNVs and CNAs data are provided, one can attempt at merging the events in a pre-processing phase. Consider for instance a mutation on a gene X acting as a tumour suppressor, and a chromosomal deletion spanning through the mutated gene (or a wider chromosomal region including it). If one cane phase the deletion to the strand where X is mutated, a single-cell dataset could look as in Supplementary Figure 22. If there is a lineage relation between cells (samples 1–3 are ancestral to 4–6, which we do not know), and we merge the two distinct events before performing the inference, then we can retrieve a progression model where we time the event “inactivation of X”. In more general scenarios of aneuploidy, or when cells are siblings and the events are not phased, creating a merged events is not a solution to the back-mutation problem.

### References

[1] Alexander Davis and Nicholas E Navin. Computing tumor trees from single cells. *Genome biology*, 17(1):113, 2016.

[2] Katharina Jahn, Jack Kuipers, and Niko Beerenwinkel. Tree inference for single-cell data. *Genome biology*, 17(1):86, 2016.

[3] Edith M Ross and Florian Markowetz. Onconem: inferring tumor evolution from single-cell sequencing data. *Genome biology*, 17(1):69, 2016.

[4] Andrew Roth, Andrew McPherson, Emma Laks, Justina Biele, Damian Yap, Adrian Wan, Maia A Smith, Cydney B Nielsen, Jessica N McAlpine, Samuel Aparicio, et al. Clonal genotype and population structure inference from single-cell tumor sequencing. *Nature methods*, 13(7):573–576, 2016.

[5] Loes Olde Loohuis, Giulio Caravagna, Alex Graudenzi, Daniele Ramazzotti, Giancarlo Mauri, Marco Antoniotti, and Bud Mishra. Inferring tree causal models of cancer progression with probability raising. *PloS one*, 9(10):e108358, 2014.

[6] Daniele Ramazzotti, Giulio Caravagna, Loes Olde Loohuis, Alex Graudenzi, Ilya Korsunsky, Giancarlo Mauri, Marco Antoniotti, and Bud Mishra. Capri: efficient inference of cancer progression models from cross-sectional data. *Bioinformatics*, 31(18):3016–3026, 2015.

[7] Giulio Caravagna, Alex Graudenzi, Daniele Ramazzotti, Rebeca Sanz-Pamplona, Luca De Sano, Giancarlo Mauri, Victor Moreno, Marco Antoniotti, and Bud Mishra. Algorithmic methods to infer the evolutionary trajectories in cancer progression. *Proceedings of the National Academy of Sciences*, page 201520213, 2016.

[8] Daniele Ramazzotti. *A Model of Selective Advantage for the Efficient Inference of Cancer Clonal Evolution*. PhD thesis, arXiv preprint arXiv:1602.07614, 2017.

[9] Daniele Ramazzotti, Marco S Nobile, Paolo Cazzaniga, Giancarlo Mauri, and Marco Antoniotti. Parallel implementation of efficient search schemes for the inference of cancer progression models. In *Computational Intelligence in Bioinformatics and Computational Biology (CIBCB), 2016 IEEE Conference on*, pages 1–6. IEEE, 2016.

[10] Daniele Ramazzotti, Alex Graudenzi, Giulio Caravagna, and Marco Antoniotti. Modeling cumulative biological phenomena with suppes-bayes causal networks. *Evolutionary Bioinformatics*, 14:1176934318785167, 2018.

[11] Daphne Koller and Nir Friedman. *Probabilistic graphical models: principles and techniques*. MIT press, 2009.

[12] Patrick Suppes. *A probabilistic theory of causality*. North-Holland Publishing Company Amsterdam, 1970.

[13] David JC MacKay. *Information theory, inference and learning algorithms*. Cambridge university press, 2003.

[14] Mohammed El-Kebir, Layla Oesper, Hannah Acheson-Field, and Benjamin J Raphael. Reconstruction of clonal trees and tumor composition from multi-sample sequencing data. *Bioinformatics*, 31(12):i62–70, June 2015.

[15] Victoria Popic, Raheleh Salari, Iman Hajirasouliha, Dorna Kashef-Haghighi, Robert B West, and Serafim Batzoglou. Fast and scalable inference of multi-sample cancer lineages. *Genome Biology*, 16(1):795–17, May 2015.

[16] Jack Edmonds. Optimum branchings. *Journal of Research of the National Bureau of Standards B*, 71(4):233–240, 1967.

[17] Robert Endre Tarjan. Finding optimum branchings. *Networks*, 7(1):25–35, 1977.

[18] Robert Clay Prim. Shortest connection networks and some generalizations. *Bell Labs Technical Journal*, 36(6):1389–1401, 1957.

[19] Harold N Gabow. Path-based depth-first search for strong and biconnected components. *Information Processing Letters*, 74(3-4):107–114, 2000.

[20] C Chow and Cong Liu. Approximating discrete probability distributions with dependence trees. *IEEE transactions on Information Theory*, 14(3):462–467, 1968.

[21] Barbara L Parsons. Many different tumor types have polyclonal tumor origin: evidence and implications. *Mutation Research/Reviews in Mutation Research*, 659(3):232–247, 2008.

[22] Ke Yuan, Thomas Sakoparnig, Florian Markowetz, and Niko Beerenwinkel. Bitphylogeny: a probabilistic framework for reconstructing intra-tumor phylogenies. *Genome biology*, 16(1):36, 2015.

[23] Yong Wang, Jill Waters, Marco L Leung, Anna Unruh, Whijae Roh, Xiuqing Shi, Ken Chen, Paul Scheet, Selina Vattathil, Han Liang, et al. Clonal evolution in breast cancer revealed by single nucleus genome sequencing. *Nature*, 512(7513):155–160, 2014.

[24] You-Wang Lu, Hui-Feng Zhang, Rui Liang, Zhen-Rong Xie, Hua-You Luo, Yu-Jian Zeng, Yu Xu, La-Mei Wang, Xiang-Yang Kong, and Kun-Hua Wang. Colorectal cancer genetic heterogeneity delineated by multi-region sequencing. *PloS one*, 11(3):e0152673, 2016.

| Error Rates | | Specificity | | Sensitivity | |
|---|---|---|---|---|---|
| $\epsilon_+$ | $\epsilon_-$ | Average | SD | Average | SD |
| 0.003 | 0.07 | 0.984 | 0.020 | 0.729 | 0.172 |
| 0.004 | 0.07 | 0.984 | 0.020 | 0.725 | 0.179 |
| 0.005 | 0.07 | 0.985 | 0.020 | 0.730 | 0.176 |
| 0.006 | 0.07 | 0.984 | 0.020 | 0.723 | 0.179 |
| 0.007 | 0.07 | 0.984 | 0.020 | 0.724 | 0.175 |
| 0.003 | 0.06 | 0.984 | 0.020 | 0.729 | 0.172 |
| 0.004 | 0.06 | 0.984 | 0.020 | 0.725 | 0.179 |
| 0.005 | 0.06 | 0.985 | 0.020 | 0.730 | 0.176 |
| 0.006 | 0.06 | 0.984 | 0.020 | 0.725 | 0.178 |
| 0.007 | 0.06 | 0.984 | 0.020 | 0.726 | 0.175 |
| 0.003 | 0.05 | 0.984 | 0.020 | 0.729 | 0.172 |
| 0.004 | 0.05 | 0.984 | 0.020 | 0.725 | 0.179 |
| 0.005 | 0.05 | 0.985 | 0.020 | 0.730 | 0.176 |
| 0.006 | 0.05 | 0.985 | 0.020 | 0.727 | 0.178 |
| 0.007 | 0.05 | 0.984 | 0.020 | 0.726 | 0.175 |
| 0.003 | 0.04 | 0.984 | 0.020 | 0.729 | 0.172 |
| 0.004 | 0.04 | 0.984 | 0.020 | 0.726 | 0.178 |
| 0.005 | 0.04 | 0.985 | 0.020 | 0.730 | 0.176 |
| 0.006 | 0.04 | 0.985 | 0.020 | 0.727 | 0.178 |
| 0.007 | 0.04 | 0.984 | 0.020 | 0.726 | 0.175 |
| 0.003 | 0.03 | 0.984 | 0.020 | 0.729 | 0.172 |
| 0.004 | 0.03 | 0.984 | 0.020 | 0.726 | 0.178 |
| 0.005 | 0.03 | 0.985 | 0.020 | 0.730 | 0.176 |
| 0.006 | 0.03 | 0.985 | 0.020 | 0.727 | 0.178 |
| 0.007 | 0.03 | 0.984 | 0.020 | 0.726 | 0.175 |

Supplementary Table 4: Experiment IV. (noise-robustness). Average sensitivity and specificity (and standard deviation) of Gabow on datasets generated from the medium phylogenetic tree in Supplementary Figure 7, with $\epsilon_+ = 5 \times 10^-3$ and $\epsilon_- = 5 \times 10^-2$. We test combinations of input for $\epsilon_+$ and $\epsilon_-$ in the following ranges: $\epsilon_+ = (3, 4, 5, 6, 7) \times 10^{-3}$ and $\epsilon_- = (3, 4, 5, 6, 7) \times 10^{-2}$.

| Error Rates | | Specificity | | Sensitivity | |
|---|---|---|---|---|---|
| $\epsilon_+$ | $\epsilon_-$ | Average | SD | Average | SD |
| 0.003 | 0.07 | 0.959 | 0.030 | 0.708 | 0.181 |
| 0.004 | 0.07 | 0.958 | 0.032 | 0.696 | 0.202 |
| 0.005 | 0.07 | 0.960 | 0.029 | 0.702 | 0.188 |
| 0.006 | 0.07 | 0.962 | 0.027 | 0.709 | 0.175 |
| 0.007 | 0.07 | 0.958 | 0.031 | 0.693 | 0.197 |
| 0.003 | 0.06 | 0.959 | 0.030 | 0.699 | 0.180 |
| 0.004 | 0.06 | 0.957 | 0.032 | 0.692 | 0.203 |
| 0.005 | 0.06 | 0.962 | 0.028 | 0.707 | 0.182 |
| 0.006 | 0.06 | 0.959 | 0.030 | 0.696 | 0.190 |
| 0.007 | 0.06 | 0.958 | 0.030 | 0.691 | 0.198 |
| 0.003 | 0.05 | 0.957 | 0.029 | 0.696 | 0.174 |
| 0.004 | 0.05 | 0.955 | 0.032 | 0.686 | 0.201 |
| 0.005 | 0.05 | 0.962 | 0.027 | 0.710 | 0.181 |
| 0.006 | 0.05 | 0.960 | 0.029 | 0.698 | 0.183 |
| 0.007 | 0.05 | 0.958 | 0.031 | 0.682 | 0.207 |
| 0.003 | 0.04 | 0.956 | 0.029 | 0.698 | 0.183 |
| 0.004 | 0.04 | 0.954 | 0.031 | 0.680 | 0.203 |
| 0.005 | 0.04 | 0.962 | 0.027 | 0.718 | 0.177 |
| 0.006 | 0.04 | 0.960 | 0.030 | 0.701 | 0.187 |
| 0.007 | 0.04 | 0.958 | 0.030 | 0.684 | 0.194 |
| 0.003 | 0.03 | 0.956 | 0.029 | 0.691 | 0.195 |
| 0.004 | 0.03 | 0.956 | 0.030 | 0.693 | 0.192 |
| 0.005 | 0.03 | 0.960 | 0.027 | 0.703 | 0.184 |
| 0.006 | 0.03 | 0.961 | 0.028 | 0.703 | 0.179 |
| 0.007 | 0.03 | 0.956 | 0.031 | 0.672 | 0.195 |

Supplementary Table 5: Experiment IV. (noise-robustness). Average sensitivity and specificity (and standard deviation) of SCITE on datasets generated from the medium phylogenetic tree in Supplementary Figure 7, with $\epsilon_+ = 5 \times 10^-3$ and $\epsilon_- = 5 \times 10^-2$. We test combinations of input for $\epsilon_+$ and $\epsilon_-$ in the following ranges: $\epsilon_+ = (3, 4, 5, 6, 7) \times 10^{-3}$ and $\epsilon_- = (3, 4, 5, 6, 7) \times 10^{-2}$.

| | **Average execution time** | **Total execution time (100 experiments)** |
|---|---|---|
| **CAPRESE** | 0.03s | 19.67s |
| **PRIM** | 1.35s | 150.12s |
| **GABOW** | 1.36s | 151.56s |
| **EDMONDS** | 1.36s | 152.30s |
| **CAPRI** | 1.36s | 153.70s |
| **CHOW-LIU** | 1.39s | 154.34s |
| **SCITE** | 4.43s | 505.70s |
| **OncoNEM** | 59.23s | 6037.22s |

Supplementary Table 6: Average and total execution times for 100 distinct experiments, with the parameters described in Section 6.

**Sensitivity**

| | CHOW-LIU | | EDMONDS | |
|---|---|---|---|---|
| **Sample Size ($m$)** | Median | St. Dev. | Median | St. Dev. |
| 100 | 0.78 | 0.27 | 0.6 | 0.2 |
| 500 | 1 | 0.16 | 0.74 | 0.25 |
| 1000 | 1 | 0.07 | 0.89 | 0.15 |
| 5000 | 1 | 0.17 | 1 | 0.18 |
| 10000 | 1 | 0.14 | 1 | 0.14 |
| 15000 | 1 | 0.17 | 1 | 0.17 |
| 20000 | 1 | 0.22 | 1 | 0.22 |

**Specificity**

| | CHOW-LIU | | EDMONDS | |
|---|---|---|---|---|
| **Sample Size ($m$)** | Median | St. Dev. | Median | St. Dev. |
| 100 | 0.97 | 0.04 | 0.98 | 0.01 |
| 500 | 0.95 | 0.12 | 0.99 | 0.01 |
| 1000 | 0.96 | 0.12 | 0.99 | 0.01 |
| 5000 | 0.97 | 0.17 | 1 | 0 |
| 10000 | 0.98 | 0.13 | 1 | 0 |
| 15000 | 0.99 | 0.15 | 1 | 0 |
| 20000 | 0.98 | 0.14 | 1 | 0 |

**Computation time (sec)**

| | CHOW-LIU | | EDMONDS | |
|---|---|---|---|---|
| **Sample Size ($m$)** | Median | St. Dev. | Median | St. Dev. |
| 100 | 2.35 | 0.42 | 2.29 | 0.41 |
| 500 | 1.89 | 0.68 | 1.85 | 0.59 |
| 1000 | 2.21 | 1.34 | 2.19 | 0.82 |
| 5000 | 6.76 | 2.78 | 6.78 | 0.59 |
| 10000 | 12.22 | 2.54 | 12.43 | 0.94 |
| 15000 | 16.79 | 2.45 | 16.94 | 0.17 |
| 20000 | 22.08 | 2.61 | 22.24 | 0.21 |

Supplementary Table 7: Median sensitivity, specificity and computation time, along with the corresponding standard deviation, for CHOW-LIU and EDMONDS. We used 100 distinct experiments of SCS data, with the following parameter settings: $n = 20$ nodes, $\epsilon_+ = 0.005$, $\epsilon_- = 0.05$, $m \in (100, 500, 1000, 5000, 10000, 15000, 20000)$.

.
